# Spatial Heterogeneity of Brain Lipids in SIV-infected Macaques Treated with Antiretroviral Therapy

**DOI:** 10.1101/2022.09.26.508302

**Authors:** Cory J. White, Andrew M. Gausepohl, Hannah N. Wilkins, Colten D. Eberhard, Benjamin C. Orsburn, Dionna W. Williams

## Abstract

Human immunodeficiency virus (HIV) infection continues to promote neurocognitive impairment, mood disorders, and brain atrophy even in the modern era of viral suppression. Brain lipids are vulnerable to HIV-associated energetic strain and contribute to HIV-associated neurologic dysfunction due to alterations in lipid breakdown and structural lipid composition. HIV neuropathology is region dependent, yet there has not been comprehensive characterization of the spatial heterogeneity of brain lipids during infection that may impact neurologic function. To address this gap, we evaluated the spatial lipid distribution using matrix laser desorption/ionization imaging mass spectrometry (MALDI-IMS) across four brain regions (parietal cortex, midbrain, thalamus, and temporal cortex), as well as kidney for a peripheral tissue control, in a virally suppressed simian immunodeficiency virus (SIV)-infected rhesus macaque. We assessed lipids indicative of fat breakdown [acylcarnitines (CARs)] and critical structural lipids [phosphatidylcholines (PCs) and phosphatidylethanolamines (PEs)] across fatty acid chain lengths and degrees of unsaturation. CARs with very long-chain, polyunsaturated fatty acids (PUFAs) were more abundant across all brain regions than shorter chain, saturated or monounsaturated species. We observed distinct brain lipid distribution patterns for CARs and PCs. However, no clear expression patterns emerged for PEs. Surprisingly, kidney was nearly devoid of ions corresponding to PUFAs common in brain. PE’s and PC’s with PUFAs had little intensity and less density than other species and, only one CAR species was observed in kidney at high intensity. Overall, our study provides substantial evidence for persistent bioenergetic changes to the brain despite viral suppression, including region-dependent mobilization of CARs for oxidation and disparities in the presence of key phospholipids necessary for maintaining proper brain structure and function. These data indicate that region-specific interventions to restore proper lipid metabolism are essential for treating HIV neurologic disease in the era of antiretroviral therapy.

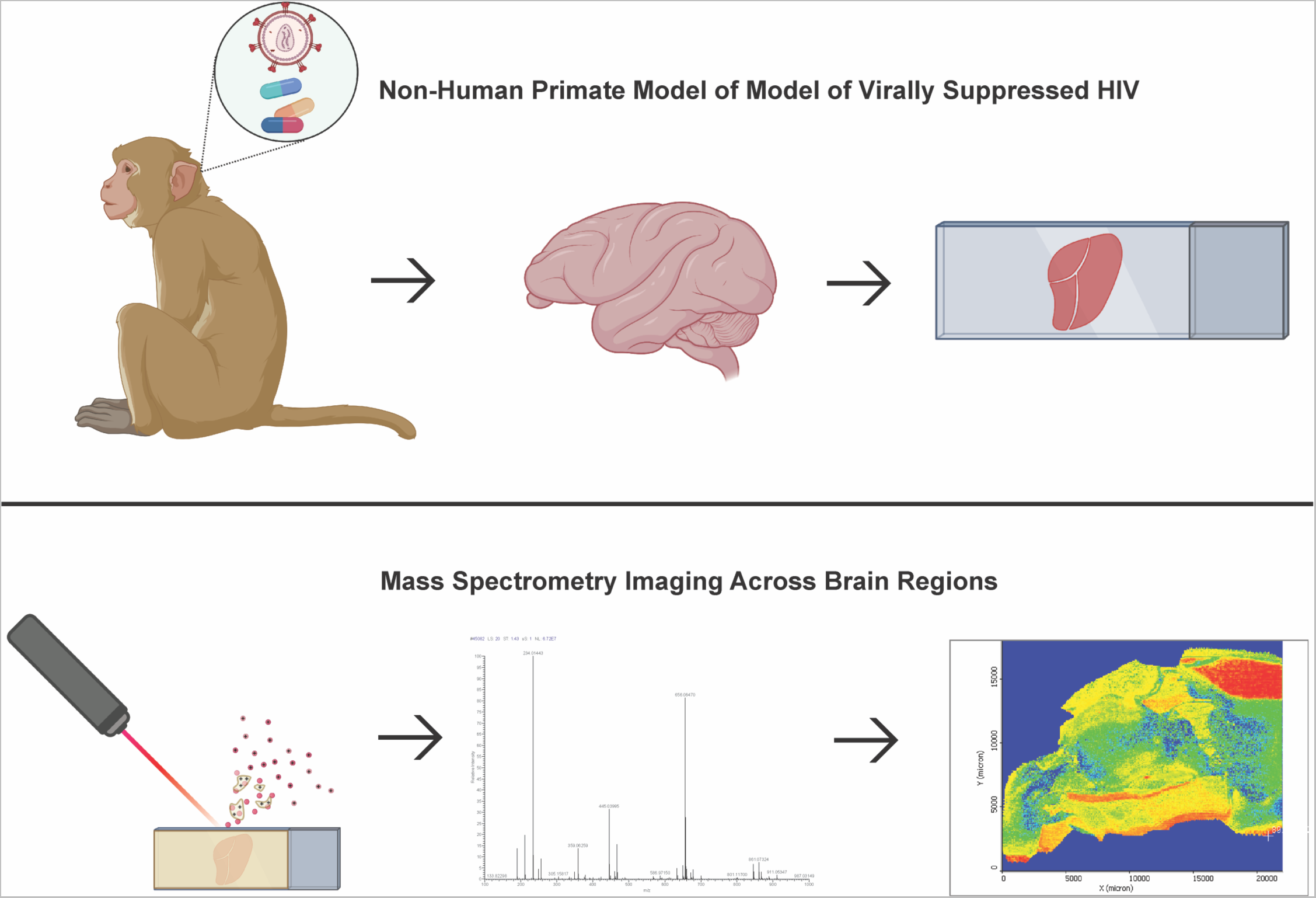
Graphical Abstract.

## INTRODUCTION

Human immunodeficiency virus (HIV) continues to pose a significant risk to neurologic health, despite the increased duration and quality of life afforded by antiretroviral therapy (ART)-mediated viral suppression ^1^. While the severity of neurologic disease has decreased in the ART era, virally suppressed people with HIV can now age with the virus, and are more susceptible to neurocognitive impairment, mood disorders, and brain atrophy, compared to uninfected individuals. Changes in the metabolic signature of the brain are substantial drivers of neurologic dysfunction. Lipid imbalance is particularly important in the CNS as the brain is approximately 60% lipid (dry weight) and has a unique composition, compared to other tissues, that is highly enriched in critical omega-3 and omega-6 fatty acids ^2, 3^ As such, lipid dysregulation contributes substantially to neurologic dysfunction ^4, 5^ as they are necessary for maintaining cellular homeostasis, proper structure, propagating action potentials, use as signaling molecules, and as a source of energy ^5–7^.

It is not surprising, then, that disruptions in brain lipid metabolism are associated with the development of neurocognitive and mood disorders in people with HIV ^8–14^. Post-mortem examination showed that the brains of people with HIV that experienced cognitive decline were enriched in sphingomyelin containing very long-chain polyunsaturated fatty acids (PUFAs) in the parietal cortex, medial frontal gyrus, and cerebellum ^9^. In primary astrocyte cultures, the HIV protein, Tat, decreased mRNA of brain cholesterol biosynthesis genes and abundance of brain cholesterol ^8^. Further, in primary astrocytes, Tat increased mRNA of oxidation genes ^15^. Another HIV protein, gp120, altered lipid composition in hippocampal neurons, where it increased very long-chain PUFAS incorporated into sphingolipids ^9^. Similarly, brain tissue of transgenic rats that overexpressed regulatory and accessory HIV proteins were enriched in very long-chain PUFAs incorporated into glycerophospholipids ^10, 16^.

Although these *in vivo* and *ex vivo* studies clearly demonstrate the contribution of lipid dysregulation to central nervous system dysfunction during HIV, prior studies have not characterized spatial differences in lipid abundance across brain regions in a model of HIV. Previous studies have profiled total lipid abundance using dissected brain regions from people with HIV without consideration of the spatial heterogeneity inherent to the brain lipid profile ^9, 10^. However, lipid distribution is not uniform across brain regions ^7, 17^, due to varying energetic demands and functional roles, as well as regional differences in structural lipid composition ^18, 19^. Similarly, region-dependent neuropathological hallmarks of HIV are well recognized, where certain brain regions are preferentially vulnerable to HIV-induced damage while others remain spared ^20–31^. For example, areas of the cortex and the basal ganglia are more prone to white matter atrophy while areas of cortex and thalamus are prone to gray matter atrophy ^23, 31^. No studies have characterized the spatial distributions of brain lipids using a model of HIV, despite strong evidence of region-dependent HIV neuropathology and the existing region-specific differences in the brain’s lipid spatial profile under normal conditions necessary to meet energetic demands, supply structural needs, and maintain metabolic homeostasis.

While powerful methods have been used for lipid metabolism determinations in the brain during HIV, they were unable to characterize the spatial lipid distribution. For example, liquid/gas chromatography uses homogenized tissue to perform quantitative abundance of specific lipids. However, it does not provide spatial information about the tissue lipid composition. Conversely, magnetic resonance spectroscopy can spatially image small metabolites in tissue, but cannot discern larger metabolites, including most common lipids. To overcome these limitations, matrix assisted laser desorption ionization imaging mass spectrometry (MALDI-IMS) has emerged as a powerful technique with the capacity to perform spatial imaging within tissues while providing the specificity required to analyze individual lipid species. Indeed, this tool combines amongst the best aspects of several other techniques commonly used to evaluate brain lipid metabolism ^7, 32–34^. MALDI-IMS generates density maps for hundreds of ions that simultaneously characterizes spatial density, relative quantification, and discernment of individual lipid species. Due to their propensity to ionize well, various lipid classes, including membrane lipids, simple fatty acids and their derivatives, and signaling lipids are great candidates for characterization using MALDI-IMS ^32, 35^.

In this study we leverage the innovative and informative MALDI-IMS method in conjunction with our virally suppressed simian immunodeficiency virus (SIV)-infected rhesus macaque model to evaluate the spatial heterogeneity of brain lipids in a model of modern HIV infection in the absence of comorbid disease or dietary and activity differences, which is a substantial confounding factor when performing human studies. We focused on key classes of lipids related to lipid utilization [acylcarnitines (CARs)] and cellular structure/homeostasis [phosphatidylcholines and phosphatidylethanolamines (PCs and PEs)] and evaluated their distribution in four brain regions: parietal cortex, midbrain, thalamus, and temporal cortex. Kidney was also assessed to serve as a peripheral control with known HIV pathology ^36^. We identified distinct lipid distribution patterns across diverse brain regions that varied according to fatty acid chain length and degree of unsaturation. Unexpectedly, we identified prominent gaps in representation of ions corresponding to some lipid species in brain regions, wherein they were essentially undetectable. These findings demonstrate both the breadth and spatial pattern of species of membrane phospholipids and intermediates of fat breakdown found in the virally suppressed brain. Our study demonstrates the need for targeted therapeutics to restore region-specific changes in lipid metabolism linked to neurologic decline in people living with HIV.

## EXPERIMENTAL SECTION

### Animal Ethics Statement

All procedures were performed in accordance with the *NIH’s Guide for the Care and Use for Laboratory Animals,* the *US Department of Agriculture Animal Welfare Act*, and under the approval of the Johns Hopkins Medical School Animal Care and Use Committee.

### SIV-infected Rhesus Macaque Model

We used a well-established SIV-infected rhesus macaque (*Macaca mulatta*) model optimal for studying brain pathology, immunological compromise, viral replication, and the impact of therapeutics on HIV outcome ^37–52^. The present study was conducted using samples harvested from one juvenile 4-year-old, 6.1 kg SIV-infected, ART-treated rhesus macaque male ^37^.

The rhesus macaque was inoculated intravenously using SIVmac251 as previously described ^37, 38^. Two weeks after SIV infection, the rhesus macaque was treated daily with intraperitoneal injections of combined ARTs consisting of TFV (20 mg/kg), DTG (25 mg/kg), and FTC (50 mg/kg). The ARTs were donated from Gilead and ViiV Healthcare. This model achieved peak viremia at 14 days post inoculation and accomplished complete viral suppression by day 42 post inoculation (28 days after ART initiation) ^37^.

The macaque was monitored to ensure there were no signs of pain or distress. Although not necessary for this rhesus macaque, we followed criteria for humane euthanasia prior to planned endpoint if any of the following were observed: a) weight loss of greater than 15%, b) CD4+ T-lymphocytes count less than 5% of baseline level, c) clinical signs of neurological disease, d) intractable diarrhea, and opportunistic infection.

### Necropsy and Harvest

At approximately 130 days post inoculation, the juvenile SIV-infected, ART-treated rhesus macaque male was euthanized in accordance with federal guidelines and institutional policies ^37^. Euthanasia occurred with an overdose of sodium pentobarbital while under ketamine sedation (15- to 20-mg/kg intramuscular injection) before perfusion with phosphate-buffered saline (PBS) (Gibco) to remove blood from tissues ^37, 38, 53, 54^. The necropsy was performed following protocols established within the Retrovirus Laboratory of the Department of Molecular and Comparative Pathobiology at Johns Hopkins Medicine, in accordance with the American Veterinary Medical Association guidelines for euthanasia of animals ^37, 38, 53, 54^. After harvest, the whole brain was placed in cold 2.5% agarose, and 4mm coronal sections were obtained. Brain regions (parietal cortex, temporal cortex, thalamus, and midbrain) were further cut from 4mm coronal sections. The fresh kidney was cut into four longitudinal sections. Brain regions (parietal cortex, temporal cortex, thalamus, and midbrain) and kidney were fresh frozen using liquid nitrogen in optimum cutting temperature medium (OCT) (Sakura Finetek Inc, Torrance, CA) and stored at -80°C until cryosectioning ^38^.

### Materials and Chemicals for MALDI

The MALDI matrix, a-cyano-4-hydroxycinnamic acid (CHCA), was obtained from MilliporeSigma (St. Louis, MO). The CHCA matrix is widely suitable for lipids ^55^. All solvents and other chemicals used were either reagent or high-performance liquid chromatography (HPLC) grade, and purchased from Fisher Scientific (Hampton, NH), unless otherwise specified.

### Tissue Sections and Matrix Application

The OCT-embedded brain and kidney were sectioned to 10 μm onto Superfrost Plus Microscope Slides (Fisher Scientific, Pittsburgh, Pennsylvania) at -20°C using a Leica CM3050S cryostat (Leica Biosystems, Buffalo Grove, IL). Matrix [CHCA, 10 mg/ml in ACN:H2O (50:50, v/v)] with 0.03% (v/v) TFA was applied to each slide using a TM-Sprayer (HTX Technologies, LLC, Chapel Hill, NC) at a flow rate of 100 mL/min. The TM-Sprayer was operated at an air pressure (N2) of 10 psi, spray nozzle velocity of 1200 mm/min, track spacing at 2 mm, and spray nozzle temperature at 80°C. For homogeneous deposition of matrix onto each slide, we used 16 passes (matrix deposition cycles) ^56, 57^.

### MALDI Imaging

The MALDI-MSI data were acquired using a LTQ Orbitrap XL (Thermo Fisher Scientific, Bremen, Germany), equipped with a Fourier transform mass spectrometer (FTMS) with the MALDI ion source fitted with a direct beam N2 laser (l 5 337.7 nm). Data acquisition was performed using the positive ion mode of the MS instrument at a mass range of m/z 100–1000 Da. Mass resolution of the FTMS analyzer was 60,000. The CHCA matrix was used at 10 mg/ml for ionization of lipids and a laser energy of 7.8 mJ. The camera from the MALDI instrument was used to define regions of interest and generate MALDI position files. Ion images had a spatial resolution of 50 μm. Mass spectrometry data processing and analysis were done using Xcalibur 3.0 (ThermoFisher Scientific), and ion images were generated using ImageQuest 1.0.1 software (Thermo Fisher Scientific, San Jose, CA). To confirm the capacity to measure lipids, exact mass peaks corresponding to H^+^ adducts from standards of C18:1 (Δ-cis) carnitine oleoyl L-carnitine (Avanti Polar Lipids, Inc., Alabaster, AL) and 16:0-18:1 1-palmitoyl-2-oleoyl-sn-glycero-3-phosphoethanolamine (Avanti Polar Lipids, Inc., Alabaster, AL) were confirmed in positive mode.

### Selection of Ions

Representative lipid ions were selected for known m/z values of multiple species with varying acyl-chains for three lipid classes: CARs, PCs, and PEs using Xcalibur 3.0 (ThermoFisher Scientific) ^58^. Ion values were determined for H^+^, K^+^, and Na^+^ adducts from each neutral lipid using the LIPID MAPS Lipidomics Gateway mass (m/z) calculation tool for lipid classes. For CARs, ions were imaged for H^+^, Na^+^, and K^+^ adducts of 43 species. For both PCs and PEs respectively, ions were imaged for H^+^, Na^+^, and K^+^ adducts of 74 species each. In all, a grand total of 573 ions were imaged for each individual tissue. Representative images were selected after determination of uniform expression patterns from all three adducts. Phospholipid ion notation is as such to include all possible structural isomers. For example PE(32:1) includes the structural isomers [PE(16:1/16:0), PE(18:1,14:0), etc.].

## RESULTS

### Lipid Spatial Distribution Determination

Brain lipids are necessary for maintaining homeostasis, proper structure, propagating action potentials, use as signaling molecules, and as a source of energy ^5–7^. Rather than being homogenously present, they are instead enriched in a region-specific manner to rapidly meet energetic demands, supply structural needs, and maintain metabolic homeostasis ^14, 59, 60^. Perturbations in brain lipids contribute to neurologic dysfunction during HIV, even with successful ART. However, an understanding of the mechanisms by which this occurs remains incomplete as there has not been an evaluation of the spatial localization of lipids in brain in a virally suppressed contemporary model of HIV.

To address this gap in knowledge, we evaluated brain lipid distribution by MALDI-IMS from HIV-relevant brain regions (midbrain, parietal cortex, thalamus, temporal cortex) using a well-established, virally suppressed SIV-infected, ART-treated rhesus macaque model ^23, 37^. We also performed MALDI-IMS on kidney as a point of reference for lipid distribution in peripheral tissues with known HIV-relevance as renal disorders remain prevalent even among virally suppressed people living with HIV ^61^. To examine lipid regional distribution, 10-µm coronal slices of parietal cortex, thalamus, temporal cortex, midbrain, and kidney were imaged with a 50-µm resolution (**Fig. 1A**; **Fig. 1B**) using a m/z acquisition range of 100-1000 Da (**Fig 1A**; **Fig. 1B**). We focused on three main classes of lipids of key relevance to fatty acid oxidation (CARs) and cellular structure/homeostasis (PCs and PEs). For CARs, ions were imaged for H^+^, Na^+^, and K^+^ adducts of 43 species. For both PCs and PEs respectively, ions were imaged for H^+^, Na^+^, and K^+^ adducts of 74 species each. In all, a grand total of 573 ions were imaged for each individual tissue.

**Figure 1.**
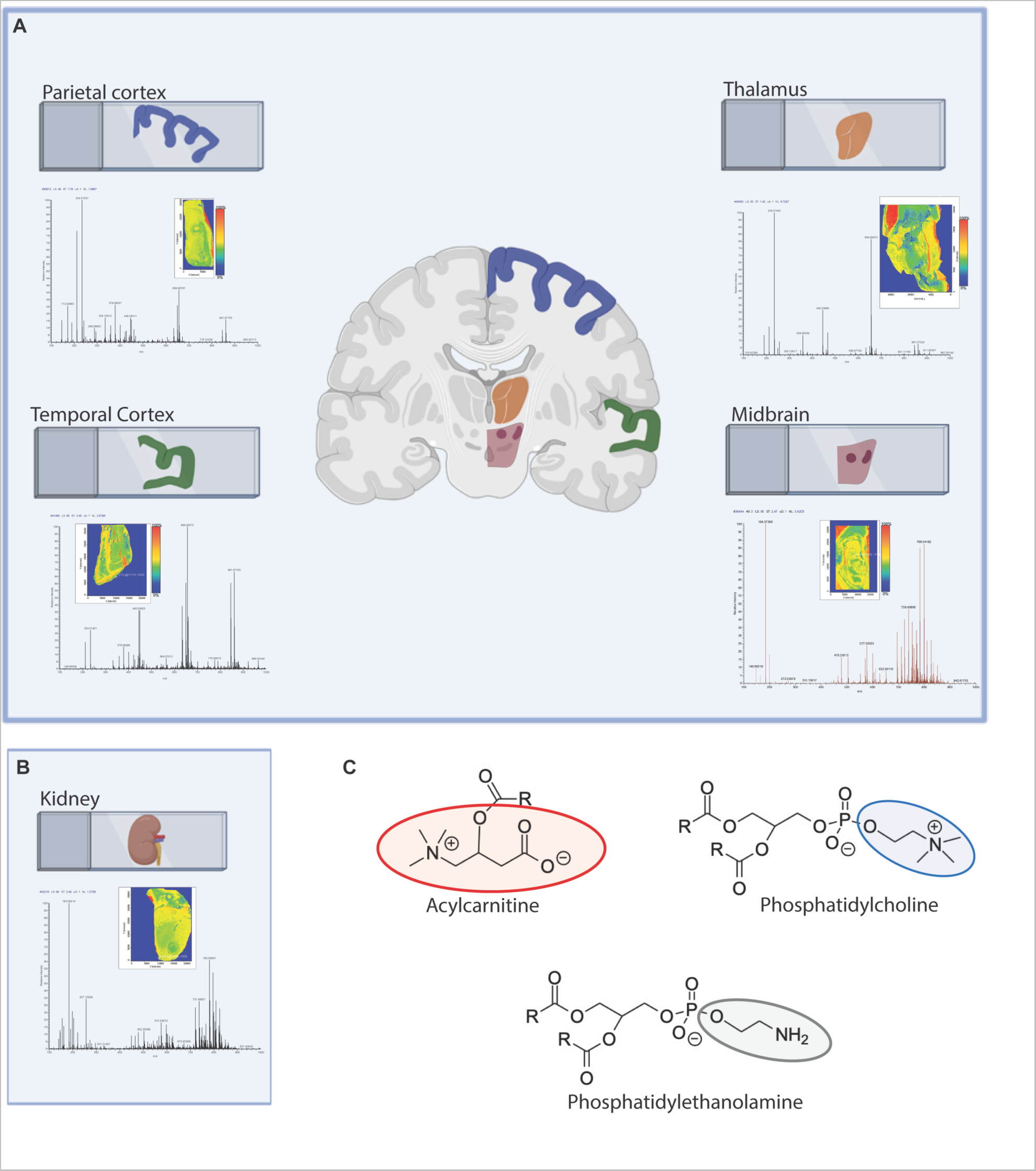
MALDI MS Imaging of SIV-infected, ART-treated macaque tissues using CHCA matrix and positive mode of the LTQ Orbitrap XL instrument. Representative full scan spectra and images of total ion counts of SIV-infected, ART-treated rhesus macaque A) brain regions: (parietal cortex, midbrain, thalamus, and temporal cortex) and B) kidney. Red signifies the highest intensity and blue the lowest of each m/z. C) CAR (red), PC (blue), and PE (gray) structures. R = acyl (fatty chain) group.

### CARs Have Unique Spatial Patterns and are Well Represented in the Brain, While Minimally Present in Kidney

We first evaluated 43 CARs, which have chain lengths ranging from 2-26 carbons, with zero to six degrees of unsaturation ^62^. There was a striking pattern wherein very long-chain CARs were highly represented across brain regions, but minimally represented in kidney. This is evidenced by the very long chain CARs, CAR20:0, CAR20:1, CAR22:3, and CAR22:4, that were highly represented across all imaged brain regions but minimally represented in kidney (**Fig. 2**). Surprisingly CAR18:3, which contains the essential linolenic acid, was minimally represented in the brain but was the only CAR prominently represented in kidney (**Fig. 2**)

**Figure 2.**
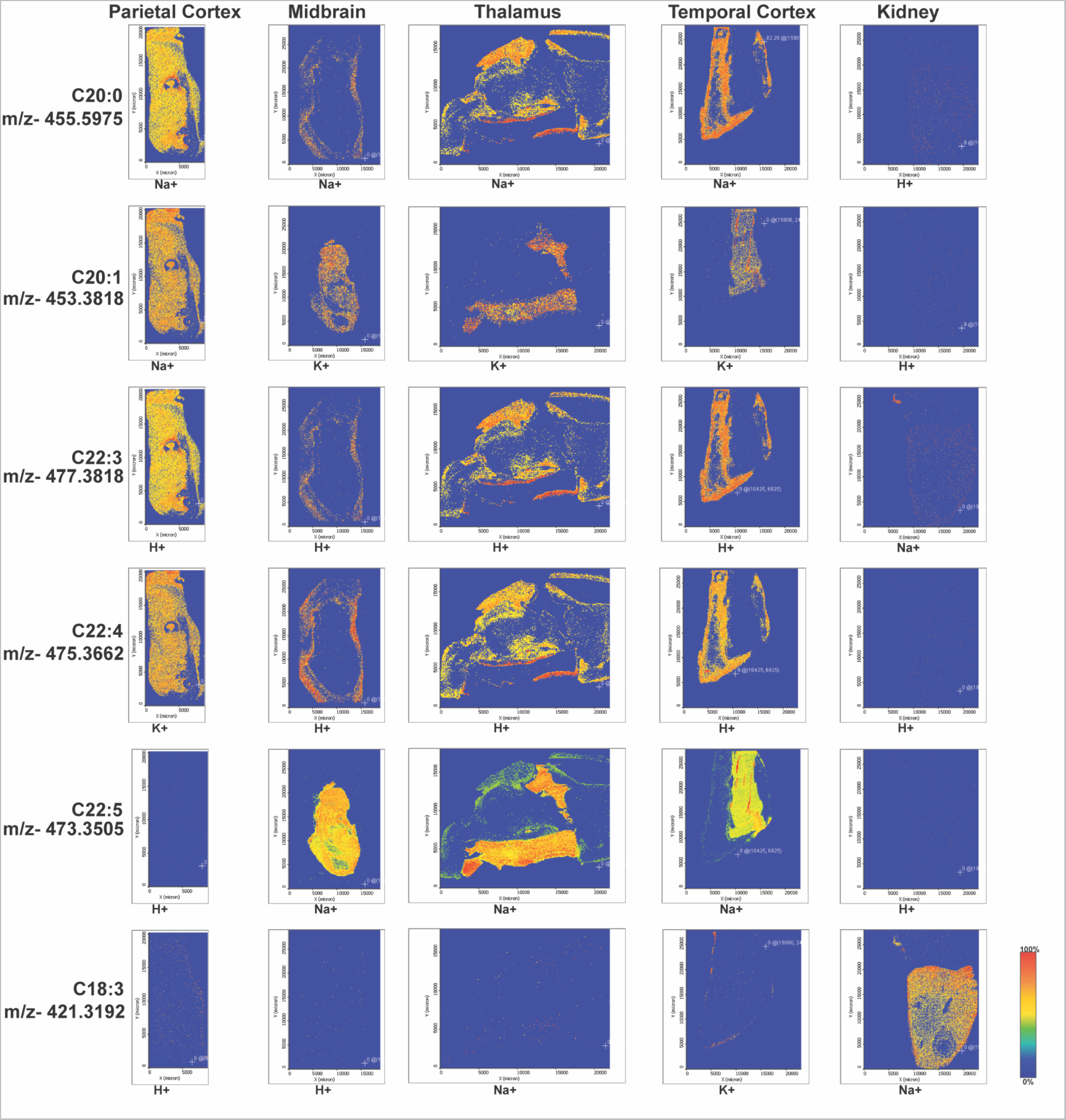
Very long-chain CARs are prevalent in the brain, but not kidney, where spatial distribution varies by species composition. Representative MALDI-IMS of SIV-infected, ART-treated rhesus macaque brain regions [parietal cortex (P. ctx), midbrain, thalamus, and temporal cortex (T. ctx)] and kidney of A) very long-chain polyunsaturated CAR(22:4) species, B) polyunsaturated CAR(18:3) species, C) varying very long-chain un-, mono-, and polyunsaturated CAR species demonstrating differences in spatial distribution. Red signifies the highest intensity and blue the lowest of each m/z.

We determined that the prominent very long-chain CARs had a high level of regional specificity both across, and within, the four brain regions (**Fig. 2**). CAR20:0, CAR22:3, and CAR22:4 were highly represented and homogenously distributed in the parietal cortex, sparse in the midbrain and temporal cortex (localizing only to the perimeter of the tissue), and were uniform in some regions with several localized clusters of intensity in the thalamus, with incomplete, but prevalent representation **(Fig. 2).** A different regional specificity was seen for CAR22:5, which was absent from the parietal cortex, yet was abundant in both midbrain, thalamus, and temporal cortex. CAR22:5 had a unique localization where it had the highest abundance in the innermost portion of tissue but was absent at the edges. In contrast, CAR20:1 was homogenously present in the parietal cortex, and localized only to the central portions of midbrain, thalamus, and temporal cortex (**Fig. 2**).

### PCs Containing Long-chain Fatty Acids are Prominently Represented and Have Consistent Spatial Patterns Within Brain and Kidney

We next evaluated PCs, focusing on species with biologically abundant medium-, long-, and very long-chain acyl-groups (**Fig. 3**). The parietal cortex had the lowest representation of PCs, where those of all chain lengths were minimally detected (**Fig. 3A**). Conversely, in the midbrain, the PCs were highly represented and, in most cases, detectable throughout the entire tissue. In both thalamus and temporal cortex, medium-chain PCs were highly represented, and longer chain length PCs were less represented.

**Figure 3.**
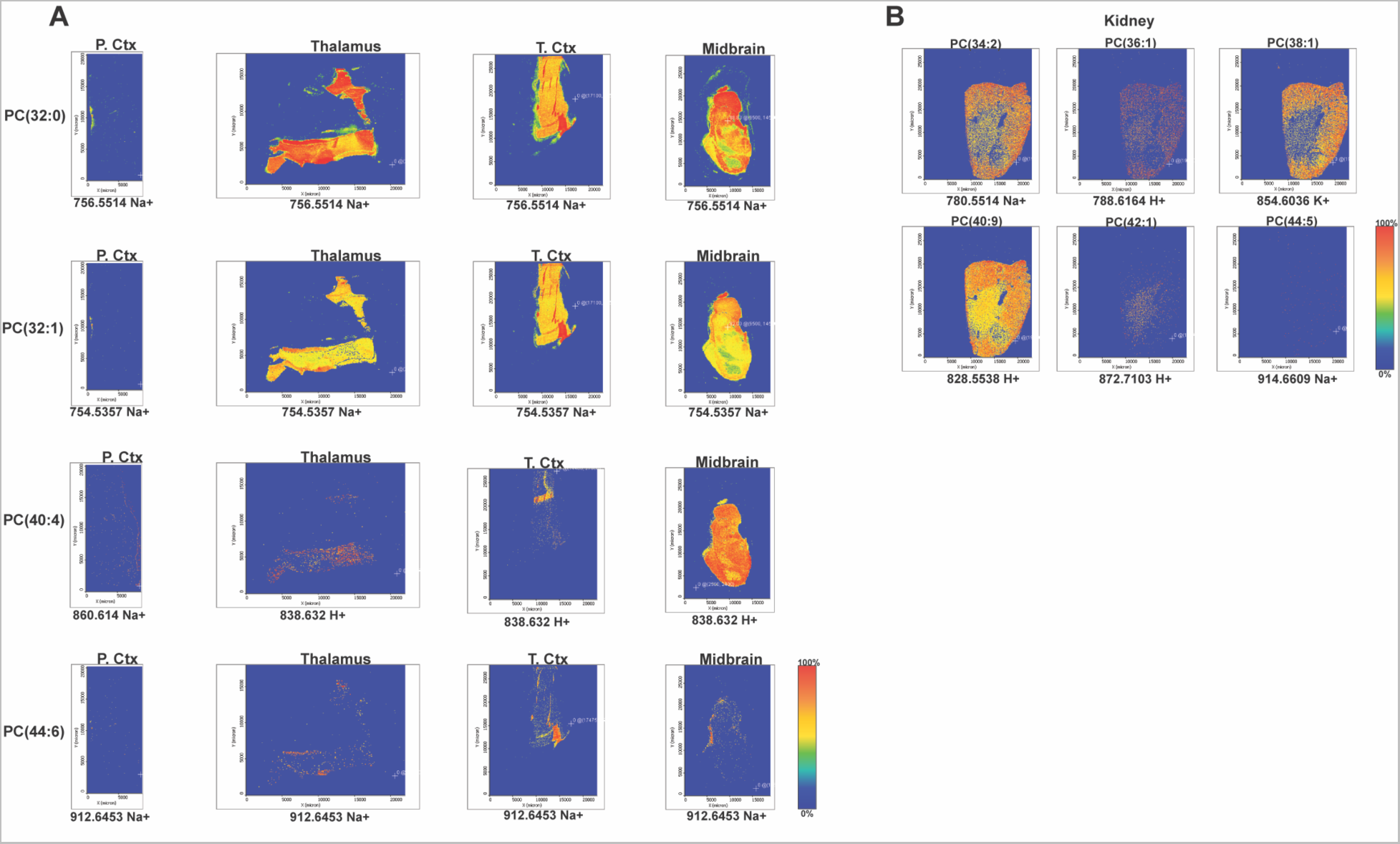
Lower chain-length PCs are more abundant in both kidney and across brain regions. Representative MALDI-IMS of SIV-infected, ART-treated rhesus macaque brain regions of A) m/z values shared by representative PCs of similar chain length in A) the following brain regions: parietal cortex (P. ctx), thalamus, temporal cortex (T. ctx), and midbrain as well as B) in kidney. Red signifies the highest intensity and blue the lowest of each m/z.

Of all four brain regions, the temporal cortex had the most unique and variable PC localization patterns.

Both PC(32:0) and PC(32:1) were abundant in a large, central portion of the temporal cortex, while PC(40:4) localized only to a relatively small area of the tissue. Similarly, PC(44:6) was mainly observed at a higher density at another relatively small area of the temporal cortex (**Fig. 3A**). Interestingly, in the thalamus, the PCs had a spatial distribution that was comparable to that of some of the CARs (CAR20:1 and CAR22:5), with greatest representation of ions being restricted to the uniquely patterned internal zone (**Fig. 3A**; **Fig. 2)**.

Of interest, there was one notable observation that occurred across brain regions, namely that very-long chain PCs were less represented (**Fig. 3A**). PCs containing very long-chain fatty acids (ex. PC(40:4) and PC(44:6)) had lower abundance overall in brain than PCs containing long-chain fatty acids (ex. PC(32:0) and PC(32:1)). One exception to this was observed in midbrain which showed high representation of ions corresponding to PCs containing long-chain fatty acids (PC(32:0) and PC(32:1)), as well as ions corresponding to PC(40:4), which contains very long-chain fatty acids, in a large, central portion of the tissue (**Fig. 3A**).

Conversely, all selected ions corresponding to PCs were minimally represented in parietal cortex (**Fig. 3A**) In the kidney, the spatial pattern of ions corresponding to PCs within the tissue varied more than what was observed in most imaged brain regions. In brain, PC ions for most species were concentrated in distinct regions of tissues (**Fig. 3A**). However, the kidney lacked any consistent trend in distribution pattern. For example, PC(36:1) was sparse and uniform throughout the entire section of kidney. PC(34:2) was more densely represented on the outer part of the kidney with a greater intensity than PC(36:1) and slightly less dense on the inner part of the kidney, while PC(38:1) was even less dense in the inner part of the kidney.

Exceptionally, PC(40:9), which could contain very long-chain PUFAs, was more densely represented in both the interior and exterior than other imaged ions (**Fig. 3B**). Additionally, PC(42:1) showed the opposite trend where the interior of the kidney had a higher representation of ions than the exterior, and PC(44:5) was sparsely, but uniformly represented (**Fig. 3B**) The spatial representation of PC ions overall was still more dense in PC ions containing long-chain fatty acids compared to those with very long-chain fatty acids similar to the brain (**Fig. 3A;3B**).

### PE Species are Heterogeneously Represented Within Brain Regions and in Kidney

The spatial distributions of PE species showed less consistent patterning than that of the CARs and PCs in both brain and kidney (**Fig. 4A;4B**). As with PCs, the smaller chain PEs were more represented in three brain regions (thalamus, temporal cortex, and midbrain) and in kidney than very-long chain PEs, while larger chain PEs were less represented in both tissues (**Fig. 4A;4B**). Interestingly, PEs were not present in the parietal cortex (**Fig. S1**).

**Figure 4.**
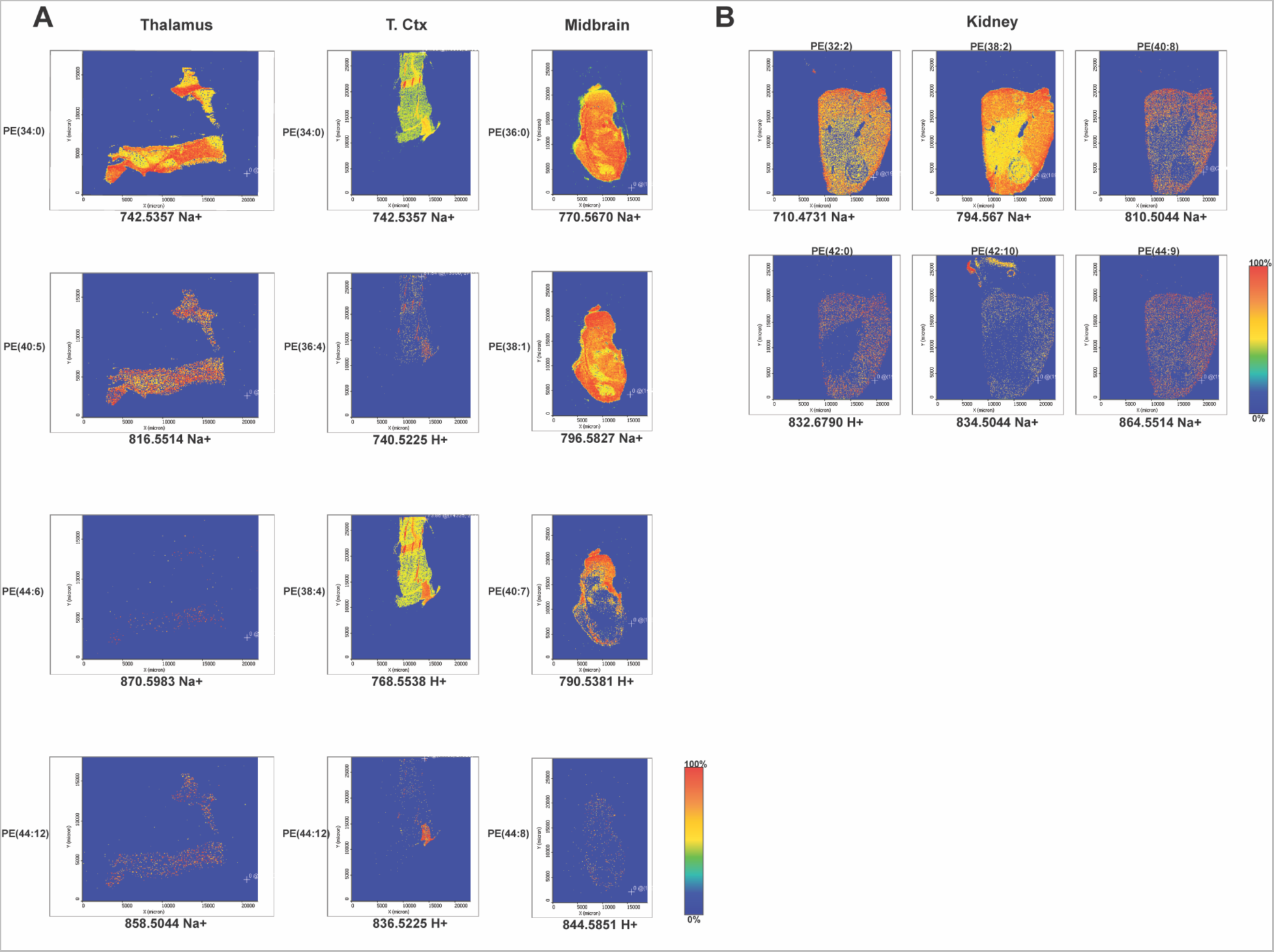
Patterns in PE spatial distribution across brain regions or within a respective tissue are less clear. Representative MALDI-IMS of SIV-infected, ART-treated rhesus macaque brain regions of A) m/z values shared by representative PEs of similar chain length in A) the following brain regions: thalamus, midbrain, and temporal cortex (T. ctx) as well as B) in kidney. Red signifies the highest abundance and blue the lowest of each m/z.

## DISCUSSION

Our study is among the first to evaluate the spatial patterns of major lipid classes related to fatty acid oxidation (CARs) and membrane phospholipid homeostasis (PCs and PEs) between and within regions of the virally suppressed brain using our non-human primate model of HIV. Using ion-specific MALDI-IMS in conjunction with our non-human primate model of HIV treated with a standard daily ART regimen, our study displays the wide breadth of lipid variation across the brain under the context of modern HIV infection. We identified distinct lipid distribution patterns within and between brain regions (and in kidney) for CARs and PCs based on differences in acyl-chain length and number of unsaturations but found no consistent patterns for PEs. CARs comprised of long-chain PUFAs were more abundant across all brain regions compared to kidney. However, the distribution pattern within each brain region differed greatly by species. For example, some species were represented along the peripheries of regions and others were restricted to the interior portions of the same region. PCs containing common saturated and monounsaturated long-chain fatty acids were most represented compared to other sized species across brain regions and in kidney. These PC species were also consistently observed in defined interior portions of brain regions including thalamus, temporal cortex, and midbrain. While PEs species containing common long-chain acyl-groups overall were also more prevalently represented, like midbrain, there were no distinct patterns of ion representation within brain regions or kidney. Surprisingly, the parietal cortex was particularly absent in many species of PCs and PEs observed in the three other brain regions.

Our findings show consistent representation of many CARs species despite viral suppression. This is of major significance because CARs are a transient intermediate of fatty acid oxidation. Fatty acid oxidation occurs in the brain minimally under normal conditions but has been reported to increase due to states of duress such as infection ^7^. Our studies support that the HIV-infected brain undergoes a considerable amount of fatty acid oxidation. Further, while PCs containing very long-chain PUFAs are less prominent than PCs containing common saturated and unsaturated long-chain fatty acids, they are still well represented across most brain regions. Previous work determined brain tissue from people with HIV that experienced severe cognitive decline had increased very long-chain fatty acids in sphingolipids, a class of structural membrane lipids, despite viral suppression ^9^. It is possible that elevations in very long-chain PUFAs contribute to, or are indicative of, cognitive decline. Determining mechanisms by which HIV/SIV and viral suppression impact these fatty acids is necessary to identify therapeutic strategies to restore changes to brain lipid homeostasis that contribute to HIV-associated cognitive decline.

### HIV Viral Proteins, Chronic Inflammation, and ARTs Impair Brain Lipid Metabolic Homeostasis

One potential driver of changes in cellular mechanisms that control lipid metabolic homeostasis are HIV viral proteins. HIV viral proteins, which are expressed despite suppression of viral replication with ARTs, greatly contribute to neurologic dysfunction. These viral proteins contribute to neurotoxicity manifesting as dendritic injuries ^63–65^, neurogenic deficits ^66^, and neuroendocrine defects ^67^. Neurotoxic HIV proteins include gp120, gp41, Tat, Nef, Rev, and Vpr ^68, 69^. Viral proteins alter brain bioenergetics in numerous ways, including Ca^2+^ dysregulation ^70–72^, modulated fatty acid oxidation ^15^, and electron transport chain complex abnormalities ^73^. They also disrupt metabolic coupling in the brain ^15^, elevate reactive oxygen species ^74, 75^, and exacerbate neuroinflammation ^63, 76^. Previous studies showed that HIV viral proteins increased fatty acid oxidation genes *in vitro* and disrupted the balance of membrane lipids ^10, 15^.

Chronic inflammation is another major contributor to CNS dysfunction that persists in people living with HIV, even with successful ART ^77–79^. Inflammation in the periphery, perivascular space, and within the brain parenchyma contribute to CNS dysfunction during HIV ^80–84^. Chronic inflammation promotes neuropathology, including accelerated brain aging, increased risk of developing psychiatric disorders, and cognitive decline ^85^. In the short term, the neuroimmune response is beneficial by promoting pathogen clearance, inducing angiogenesis, and increasing wound healing ^86^. In addition to canonical cytokines and chemokines, classes of specialized pro-resolving lipid mediators, including lipoxins, protectins, resolvins, and meresins, also promote neuroinflammation ^87^. Our study identified high abundance of the C22:5 fatty acid prominently found in specialized proresolving mediators known to alleviate neuroinflammation in specific areas of midbrain, thalamus, and temporal cortex (**Fig. 2**). This is important because previous studies determined that HIV viral proteins increased the abundance of C22:5 acid in total lipid, PE, and triglycerides ^10^. It is possible that the high abundance of C22:5 exists to counter the chronic neuroinflammation that occurs after HIV infection.

ART may also play a direct role in lipid dysregulation in brain as it adversely impacts metabolism. Systemically, it is well established that ART causes a mean weight gain of 3-7 kg within the first year of initiating treatment ^88–91^. Further, specific ART drugs or ART classes can have varying impacts on circulating lipid levels, blood pressure, and risks for developing metabolic syndromes, including diabetes mellitus. For example, classical non-nucleoside reverse transcriptase inhibitors and protease inhibitors were known for increasing the likelihood of developing dyslipidemia ^92^. Even though newer antiretrovirals do not elevate circulating lipids to the same extent, tenofovir prodrugs, including tenofovir alafenamide, increase cholesterol species ^93^. This occurs despite the known impact of HIV on diminishing total cholesterol levels ^88, 94–97^.

Conversely other ARTs, including raltegravir, dolutegravir, and bictegravir, have either a beneficial or minimal impact on the cholesterol profile after changing regimens from older protease inhibitors ^88, 98^. Hypertension, another disorder related to lipid dysregulation, is also more prevalent in individuals receiving ART. There is a 1.68-fold higher risk of developing hypertension in people living with HIV on ART in comparison to those who remain untreated ^88, 99^. However, it is not well characterized whether many of these changes to metabolic parameters occur due to the antiretrovirals directly, or because of the weight gain associated with many ARTs^88^. With the clear role of ART in promoting lipodystrophy and metabolic dysfunction systemically, it is likely that similar perturbations also occur in brain.

### MALDI-IMS Has Previously Been Used to Assess Changes in Spatial Densities of Brain Lipids Due to Other Neurologic Diseases

This study is amongst the first to spatially characterize brain lipids using a model of virally suppressed HIV infection using MALDI-IMS. While our study was limited to tissue from one rhesus macaque due to sampling availability and the precious resource they represent, our findings represent a wide breadth of membrane phospholipids and fatty acid oxidation intermediates with varying spatial density patterns both between and within brain regions. MALDI-IMS has been used previously to characterize the spatial densities of gangliosides, glycerophospholipids, sphingolipids, neuromodulatory lipids, and several fatty acid derivatives under normal conditions throughout the brain ^35, 60, 100–102^. These spatial characterization studies most commonly observed notable variations in lipid representation and density in gray versus white matter. However, density patterns amongst species are quite variable between gray and white matter. A particular lipid may be ubiquitous throughout both gray and white matter, completely segregated, or at a higher density in one with some presence remaining in the other ^60, 102^. Additional characterization of regional lipid localization remains of great importance as the brain is a spatially heterogenous organ comprised of regions with differing cellular compositions, functional roles, energetic demands, and predispositions to damage ^14, 59, 60^.

MALDI-IMS has also been used to assess spatial changes in brain lipid homeostasis during disease, including traumatic brain injuries, strokes, multiple sclerosis, Parkinson’s disease, Alzheimer’s disease, epilepsy, inborn genetic errors of metabolism, and gliomas ^32, 103–108^. For example, MALDI-IMS of coronal brain sections using a mouse model of orthotopic mouse gliomas showed increased unsaturated PCs and lyso-PCs and decreased saturated PCs as well as alterations in the balance of sphingomyelins found in tumors, as compared to wild-type mice ^103^. Our study provides a basis for the variation in lipid composition and density that exists within the virally suppressed brain. However, comparisons in spatial profile of lipids across infection status and ART use via MALDI-IMS have not yet been investigated.

### Brain Metabolic Imaging Using Other Modalities

While MALDI-IMS is not widely used to evaluate brain lipids during HIV, other modalities of brain imaging have been used to assess other metabolites. These methods were primarily limited to including positron emission tomography (PET) and magnetic resonance spectroscopy (MRS) as they can be performed clinically with minimal or moderate levels of invasiveness. Using ^18^F fluoro-deoxygluose PET (^18^F-FDG-PET), glucose uptake is significantly decreased in cerebellum and across whole brain in humans due to chronic HIV infection. Glucose uptake is further diminished in people with both a history of chronic HIV infection and chronic cocaine use in basal ganglia, cerebellum, cortex, and across whole brain ^109^. Using MRS, people with HIV have been observed to have decreased glutamate and glutamine in parietal and frontal cortex gray matter^22^. Based on the known impact of HIV on lipid-relevant gene expression, oxygen consumption, and the shifting energetic demands under duress as well as the spatial variation in HIV viral load and brain atrophy observed in people with HIV, it is imperative to comprehensively evaluate the spatial impact of HIV infection to brain lipids^14, 110^.

### Conclusions

It is important to evaluate the spatial heterogeneity inherent to brain metabolism to comprehensively evaluate the impact of HIV, and other disease states, on CNS function ^31, 111^. While other studies have investigated spatial differences in glucose and other small metabolites, our findings are the first to characterize the great variability in lipid spatial densities, which are both highly abundant and with a unique composition in brain compared to other tissues, found within and across brain regions using a model of HIV infection. These data inform the need for researchers to consider the spatial changes to brain lipids as contributors to or consequences of HIV neuropathology. For future studies, brain lipid spatial profiles should be evaluated in uninfected and SIV-infected macaques, as well as those receiving differing ART regimens ^14^. Additional considerations for future studies include the contribution of genetic variation, developmental stage/age, sex, diet composition, energy expenditure, and metabolic comorbidities, and environmental stressors to lipid enrichment and regional distribution in the brain. While limited to only one animal, our study lays the framework necessary to perform MALDI-IMS lipid evaluation in brain in the context of HIV and ART that may inform future personalized clinical interventions to maintain neurologic health in virally suppressed individuals.

## Acknowledgements

We thank our colleagues in the Johns Hopkins School of Medicine Retrovirus Laboratory in the Department of Molecular and Comparative Pathobiology for graciously providing historic tissue samples, as well as their efforts towards the husbandry and veterinary care of non-human primate models, analysis of infection and viral suppression of the animal, necropsy and sample collection, including Suzanne E. Queen, Erin N. Shirk, Brandon T. Bullock, Dr. Kelly A. Metcalf Pate, Dr. Lisa M. Mangus, Dr. Janice E. Clements, and Dr. Joseph L. Mankowski. In particular, we would like to thank Dr. Clements for providing the funding that supported procurement and goals of the original research study in which this historical animal was involved. We are also thankful for Gilead and ViiV for graciously providing antiretrovirals used in our non-human primate model.

Research reported in this publication was supported by the National Institutes of Health under award number R00 DA044838, R01 DA052859, and U01 DA058527 (DWW), R01 GM103853 (BCO), R01 AG064908 (BCO), and K00 NS118713 (CJW). Additionally, HNW was supported by T32 GM144272 granted to the Biochemistry, Cellular & Molecular Biology Graduate Program at Johns Hopkins and CDE was supported by T32 GM135083 granted to the Pharmacology and Molecular Sciences Graduate Program at Johns Hopkins. This work was supported, in part, by pilot funding provided by parent funding under the JHU NIMH Center for Novel Therapeutics for HIV-associated Cognitive Disorders P30MH075673 to Justin C. McArthur. The authors also acknowledge mentorship to DWW from the Johns Hopkins University Center for AIDS Research (P30AI094189). The content is solely the responsibility of the authors and does not necessarily represent the official views of the National Institutes of Health. Graphical images in figures were created using BioRender.

## Author Contributions

D.W.W., C.J.W., H.N.W., and C.D.E. performed experiments, A.M.G. and C.J.W. analyzed data, D.W.W. and C.J.W. were responsible for conceptualization of the study design, H.N.W., C.D.E., and B.C.O. developed the methodologies, C.J.W. and D.W.W. wrote the original draft with review and editing from all authors.

---

**Figure S1.** PEs were not observed within parietal cortex.

MALDI-IMS of SIV-infected, ART-treated rhesus macaque parietal cortex of m/z values shared by PEs in linked PowerPoint file. Red signifies the highest abundance and blue the lowest of each m/z.

