## Supplemental Figure 1 for "Spatial Heterogeneity of Brain Lipids in SIV-infected Macaques Treated with Antiretroviral Therapy"

### Slide 1
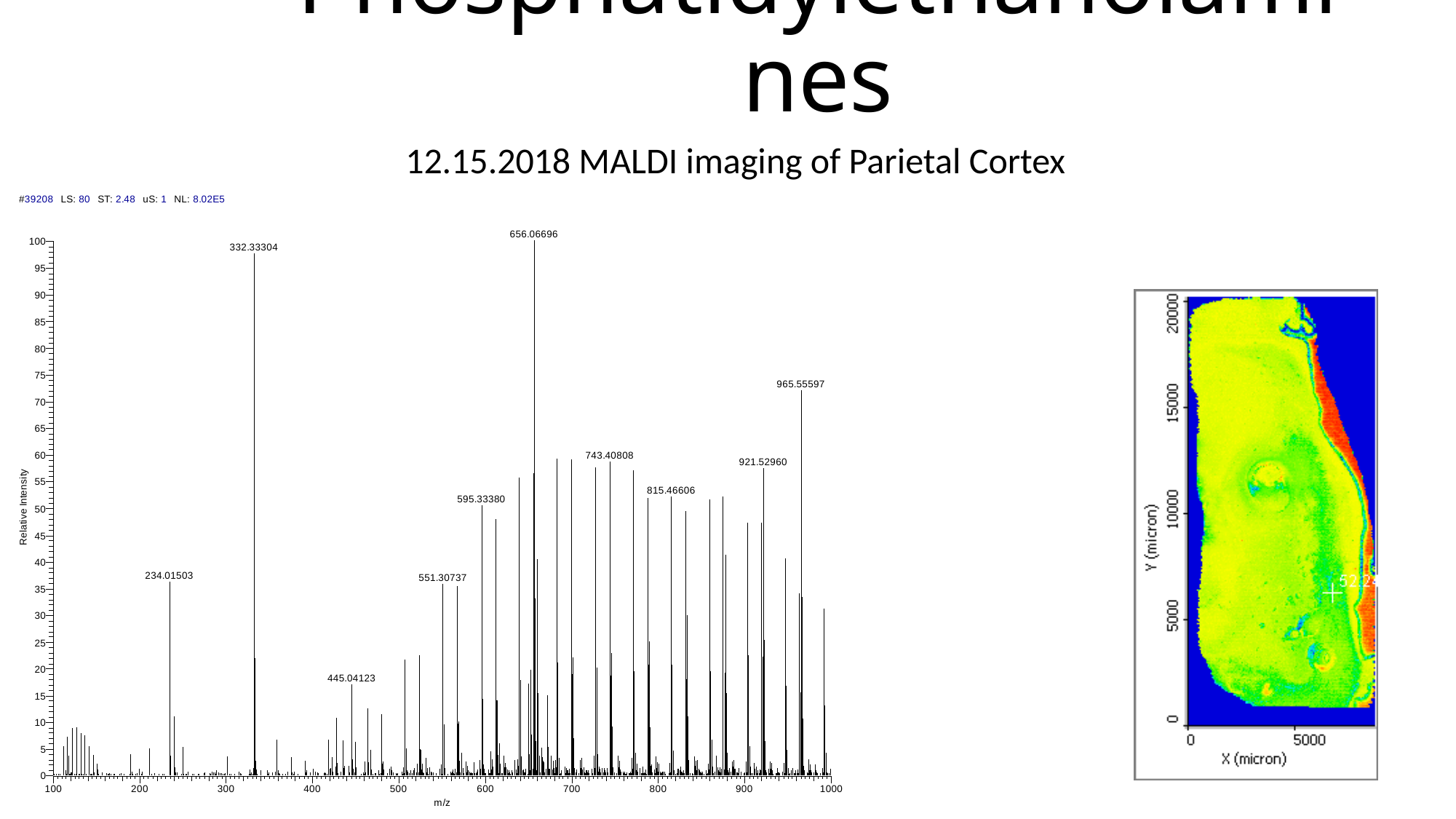

#### Phosphatidylethanolamines
12.15.2018 MALDI imaging of Parietal Cortex

### Slide 2
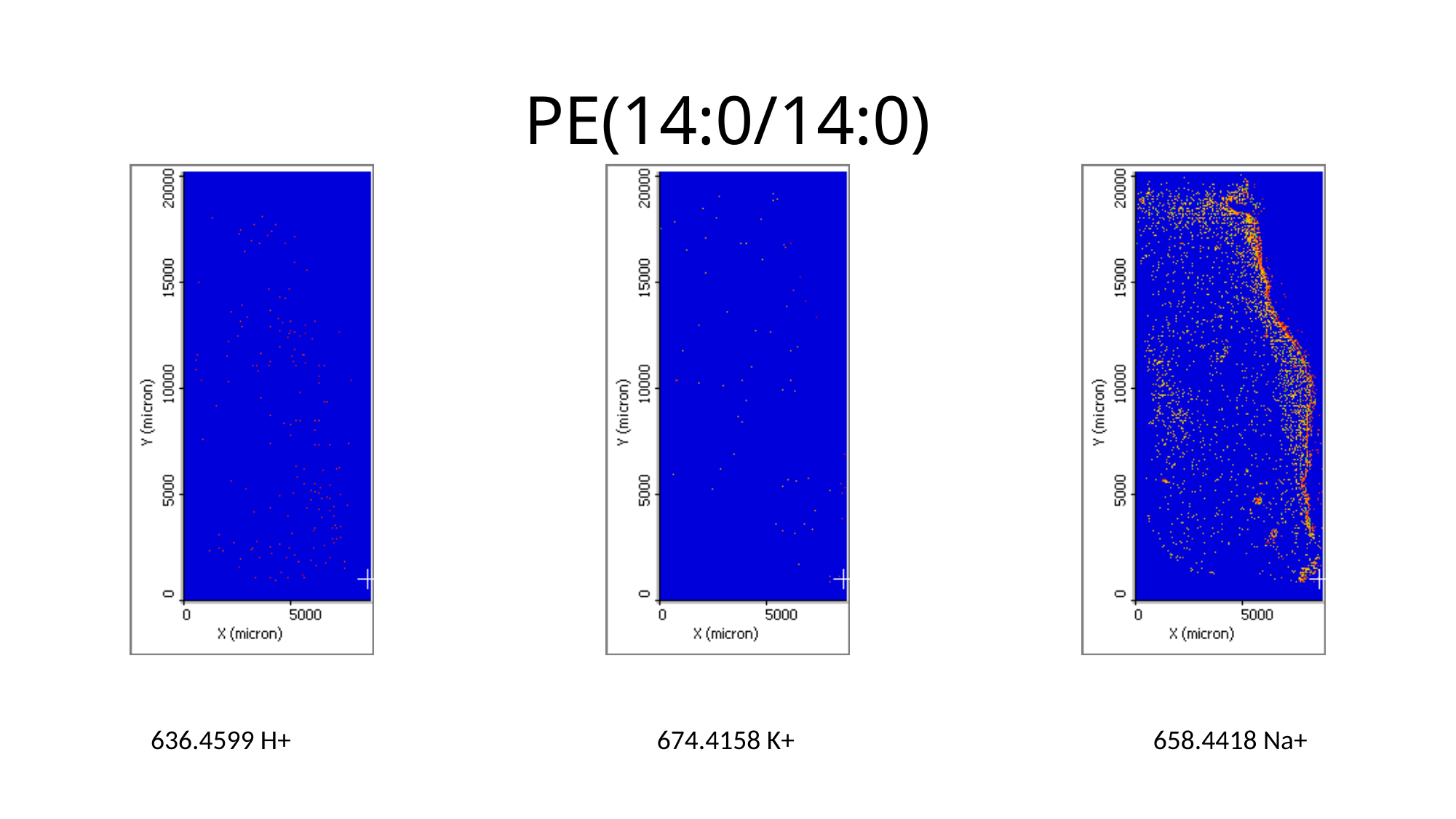

# PE(14:0/14:0)
636.4599 H+
674.4158 K+
658.4418 Na+

### Slide 3
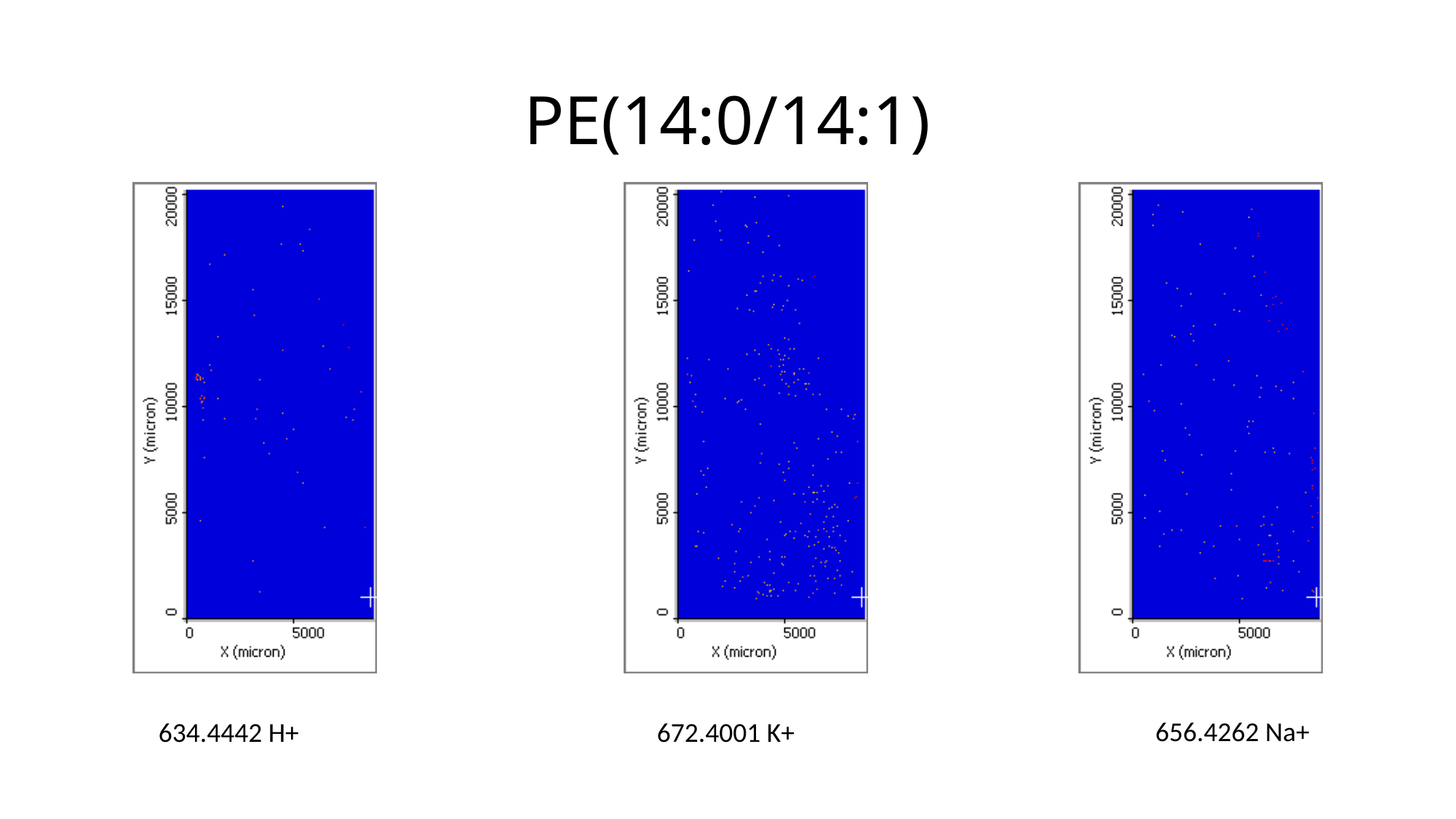

# PE(14:0/14:1)
656.4262 Na+
634.4442 H+
672.4001 K+

### Slide 4
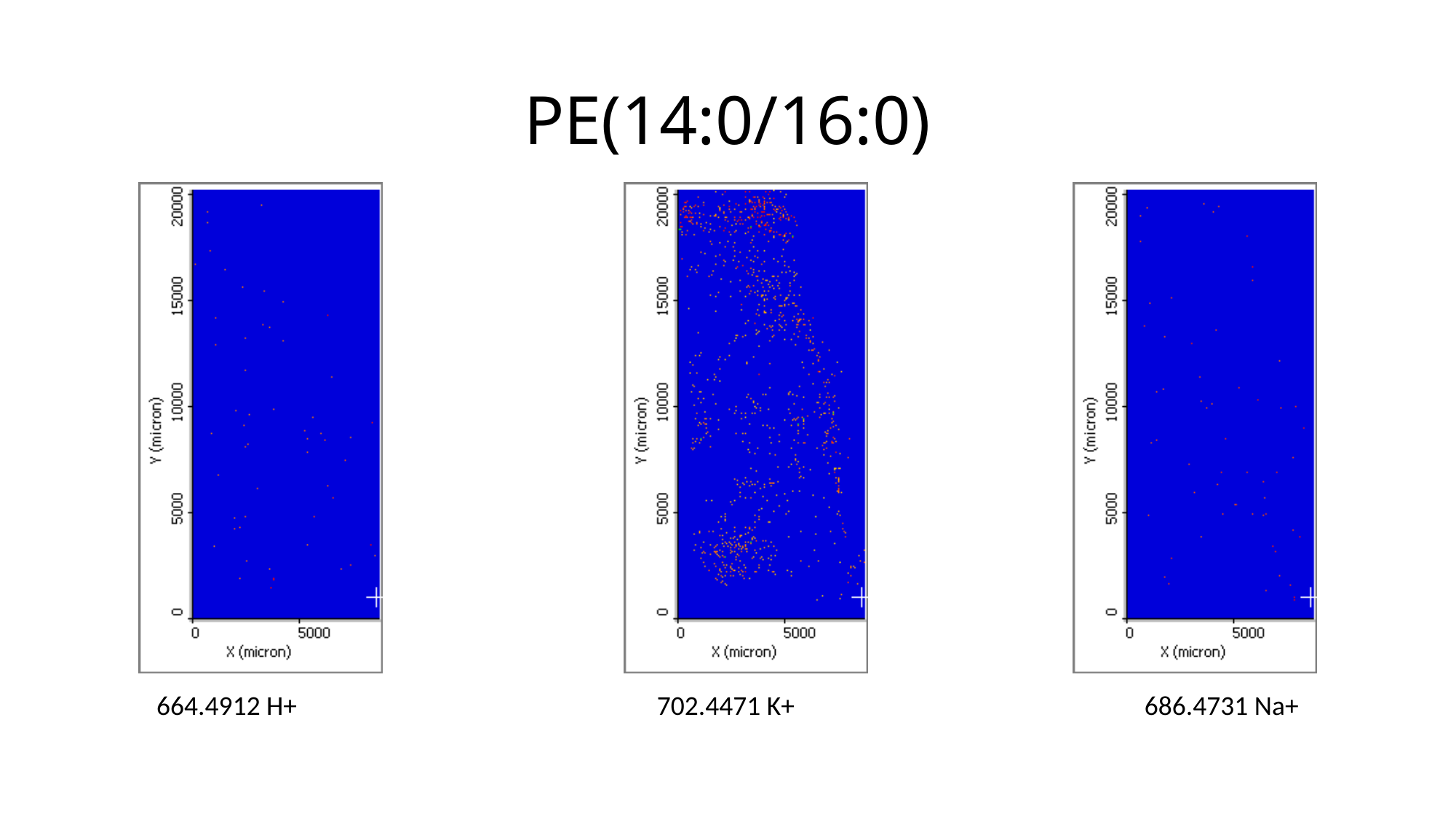

# PE(14:0/16:0)
664.4912 H+
702.4471 K+
686.4731 Na+

### Slide 5
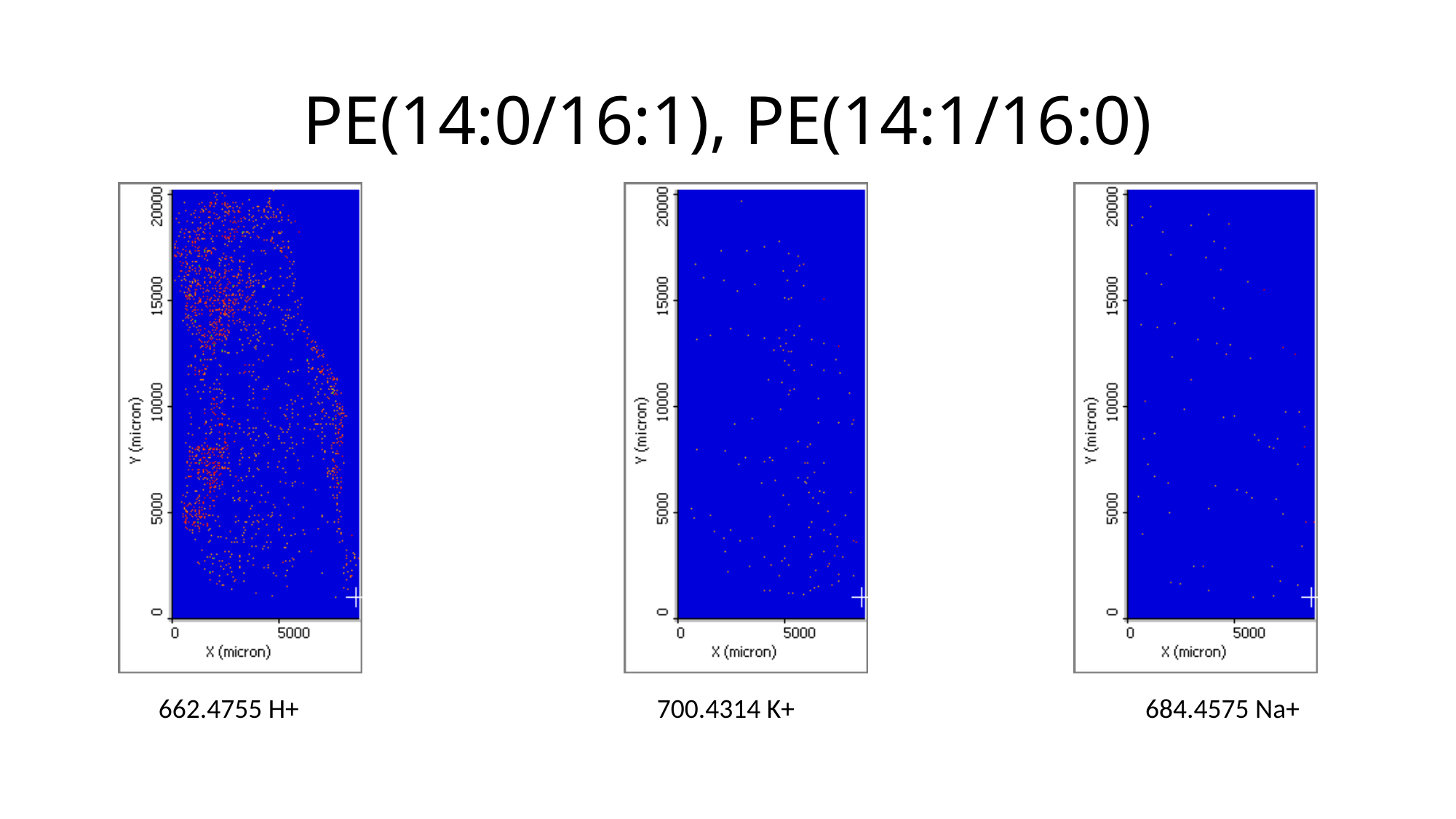

# PE(14:0/16:1), PE(14:1/16:0)
662.4755 H+
700.4314 K+
684.4575 Na+

### Slide 6
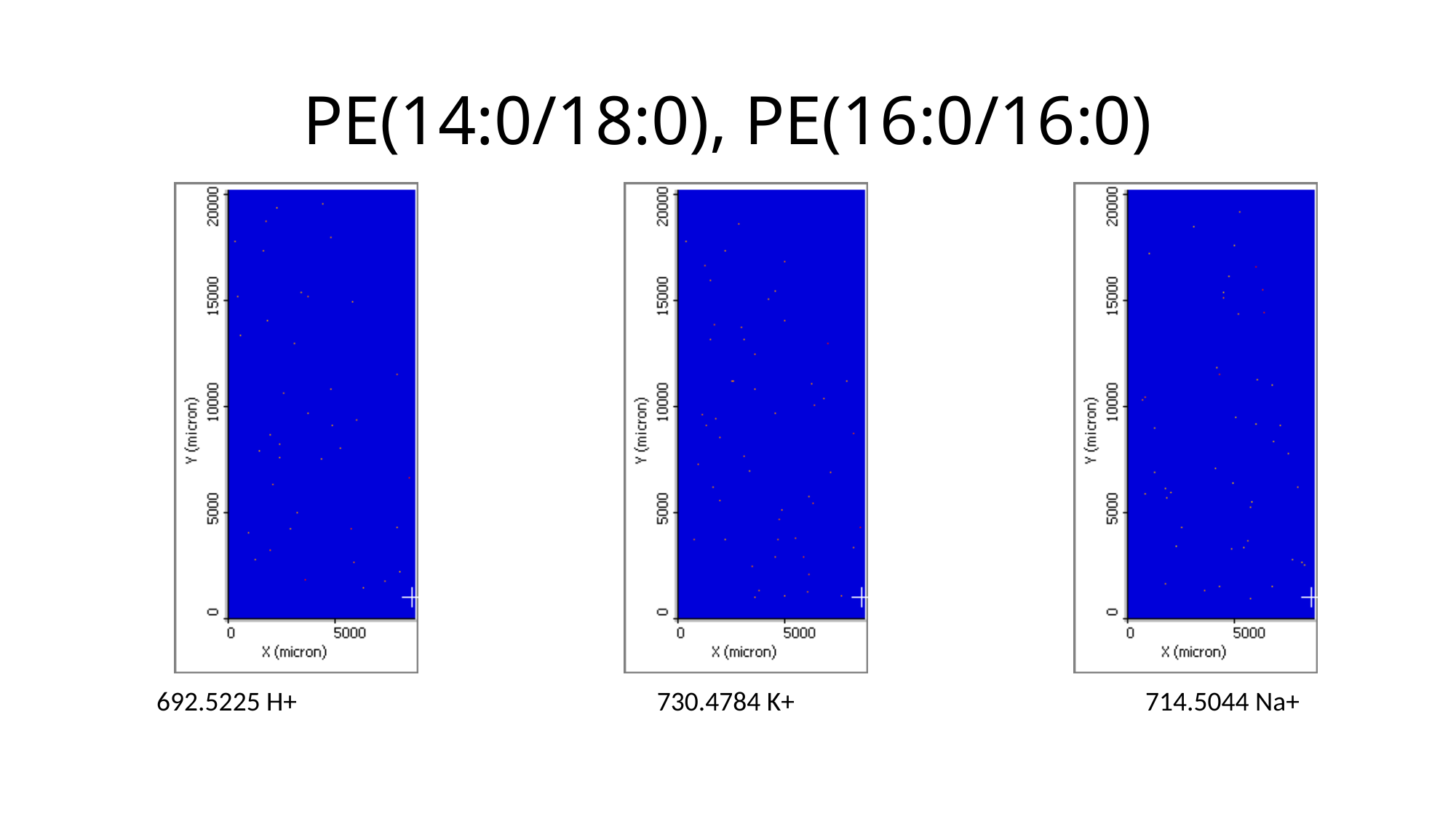

# PE(14:0/18:0), PE(16:0/16:0)
692.5225 H+
730.4784 K+
714.5044 Na+

### Slide 7
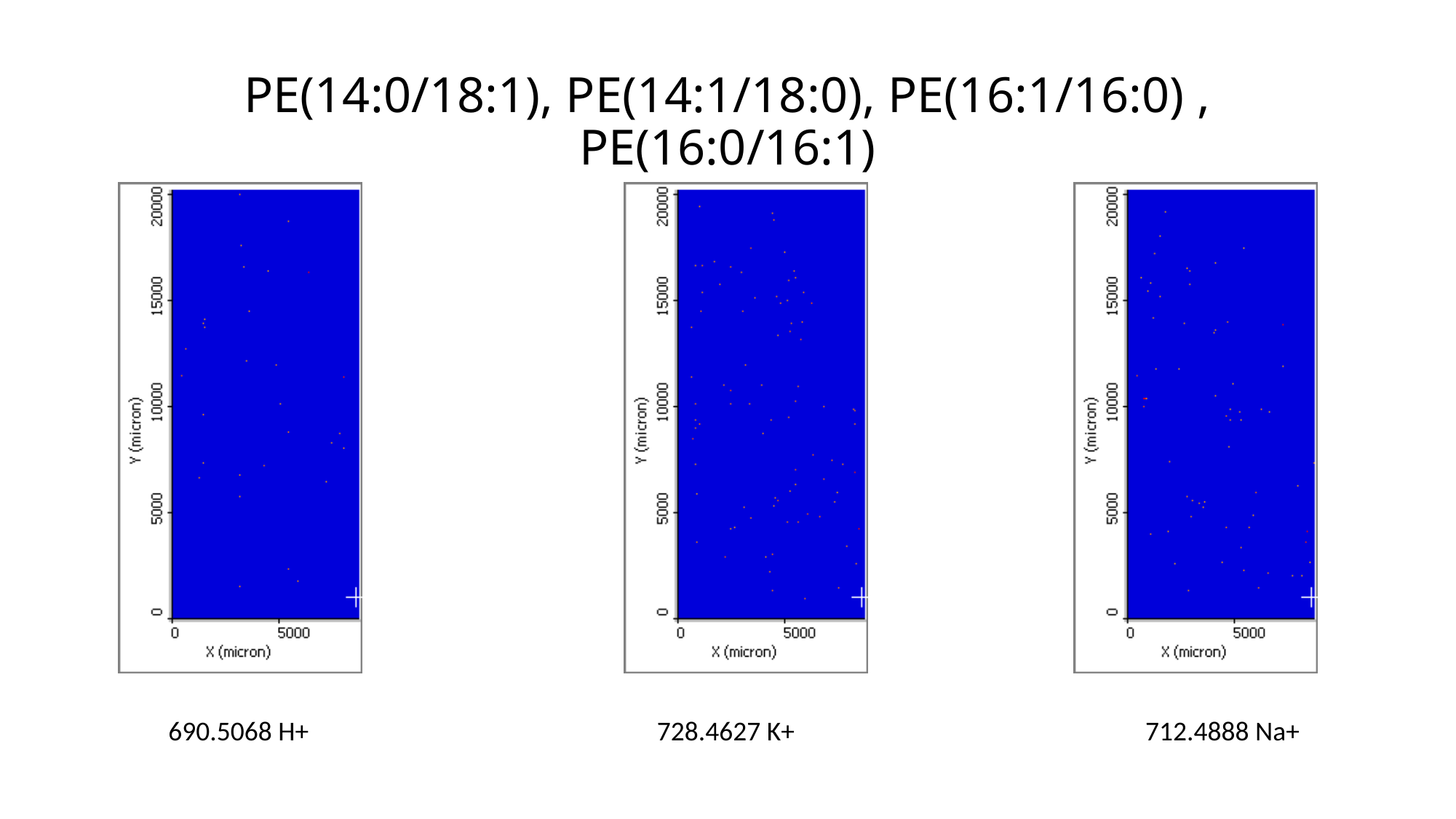

# PE(14:0/18:1), PE(14:1/18:0), PE(16:1/16:0) , PE(16:0/16:1)
690.5068 H+
728.4627 K+
712.4888 Na+

### Slide 8
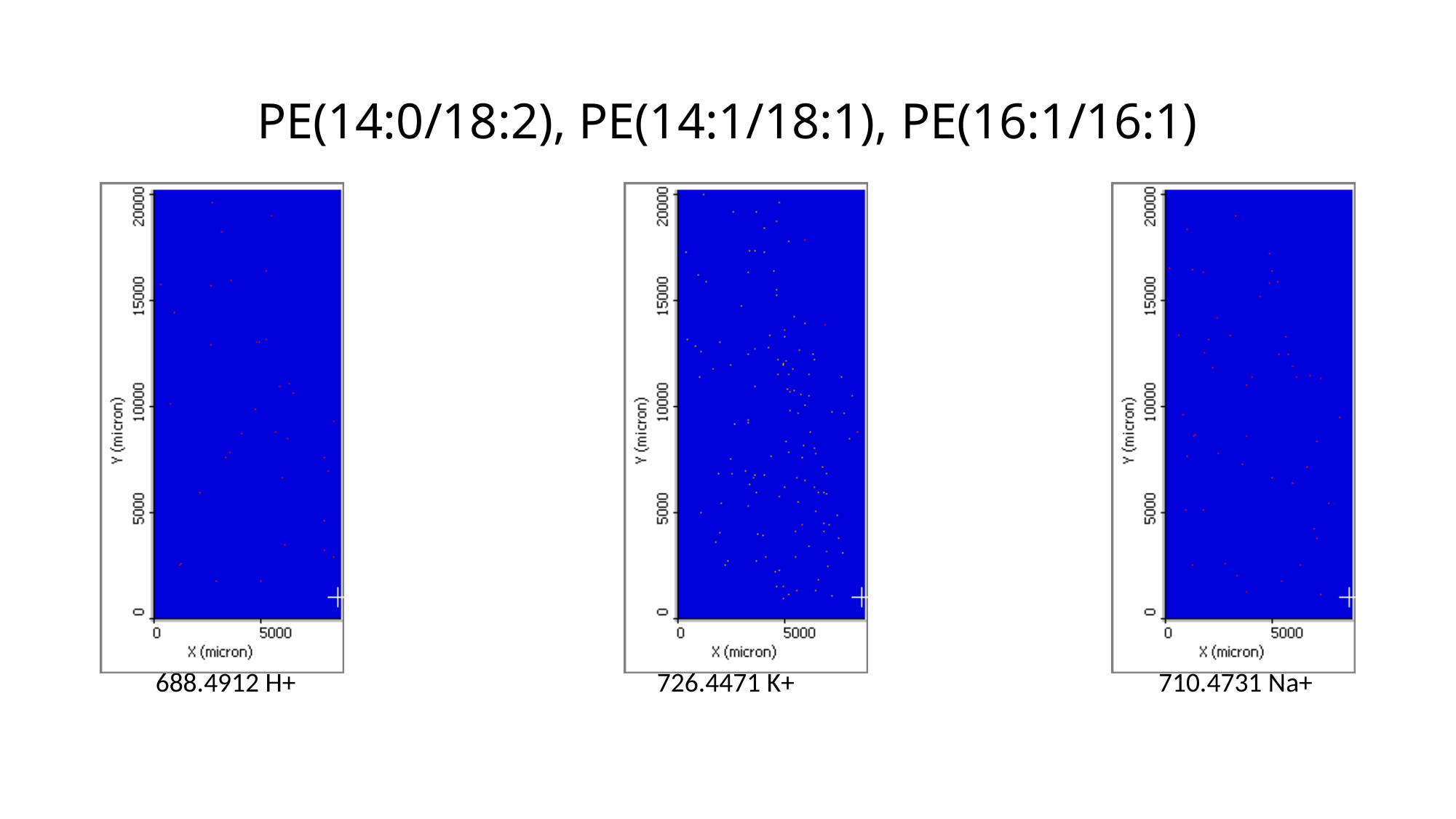

# PE(14:0/18:2), PE(14:1/18:1), PE(16:1/16:1)
688.4912 H+
726.4471 K+
710.4731 Na+

### Slide 9
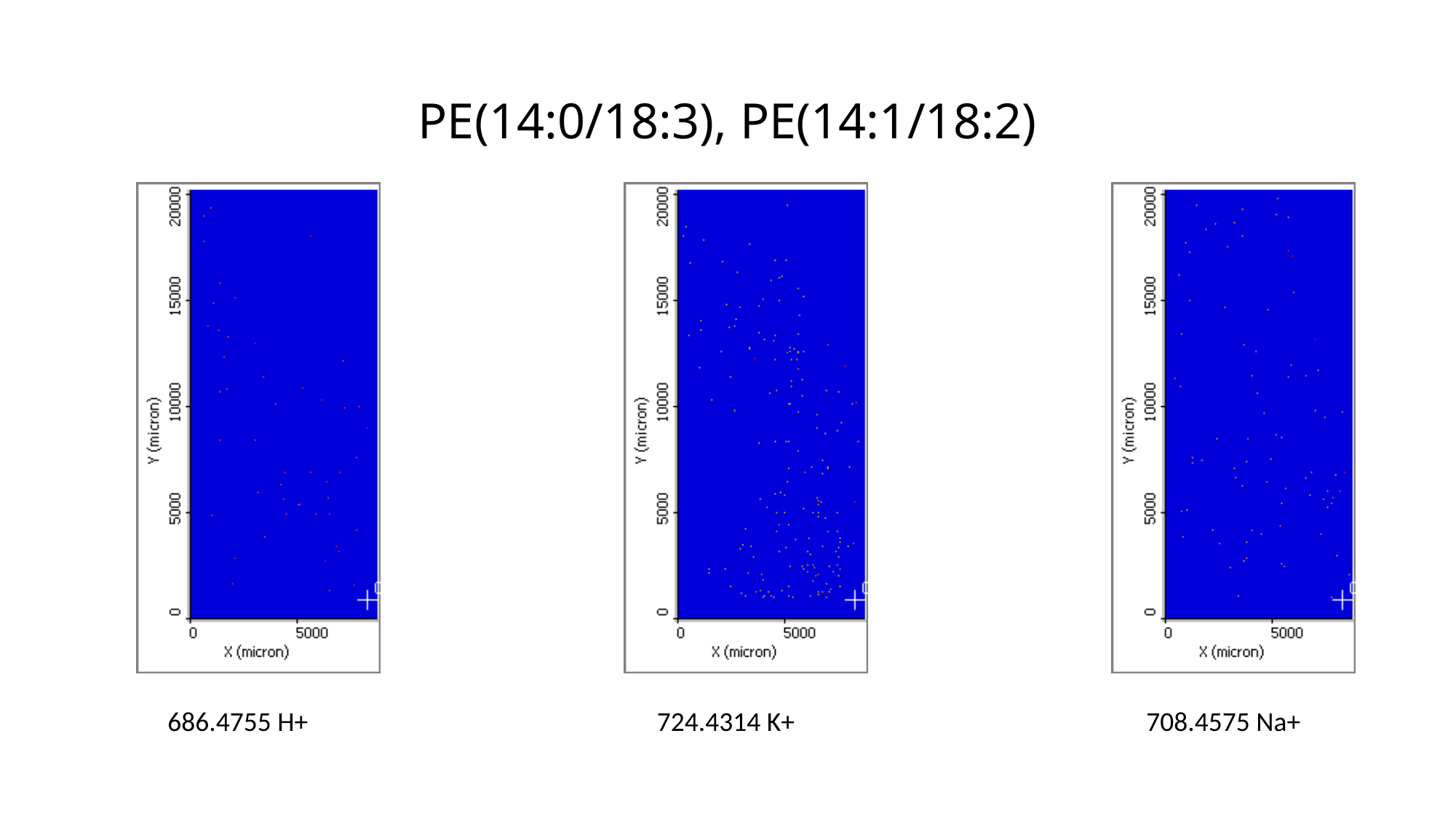

# PE(14:0/18:3), PE(14:1/18:2)
686.4755 H+
724.4314 K+
708.4575 Na+

### Slide 10
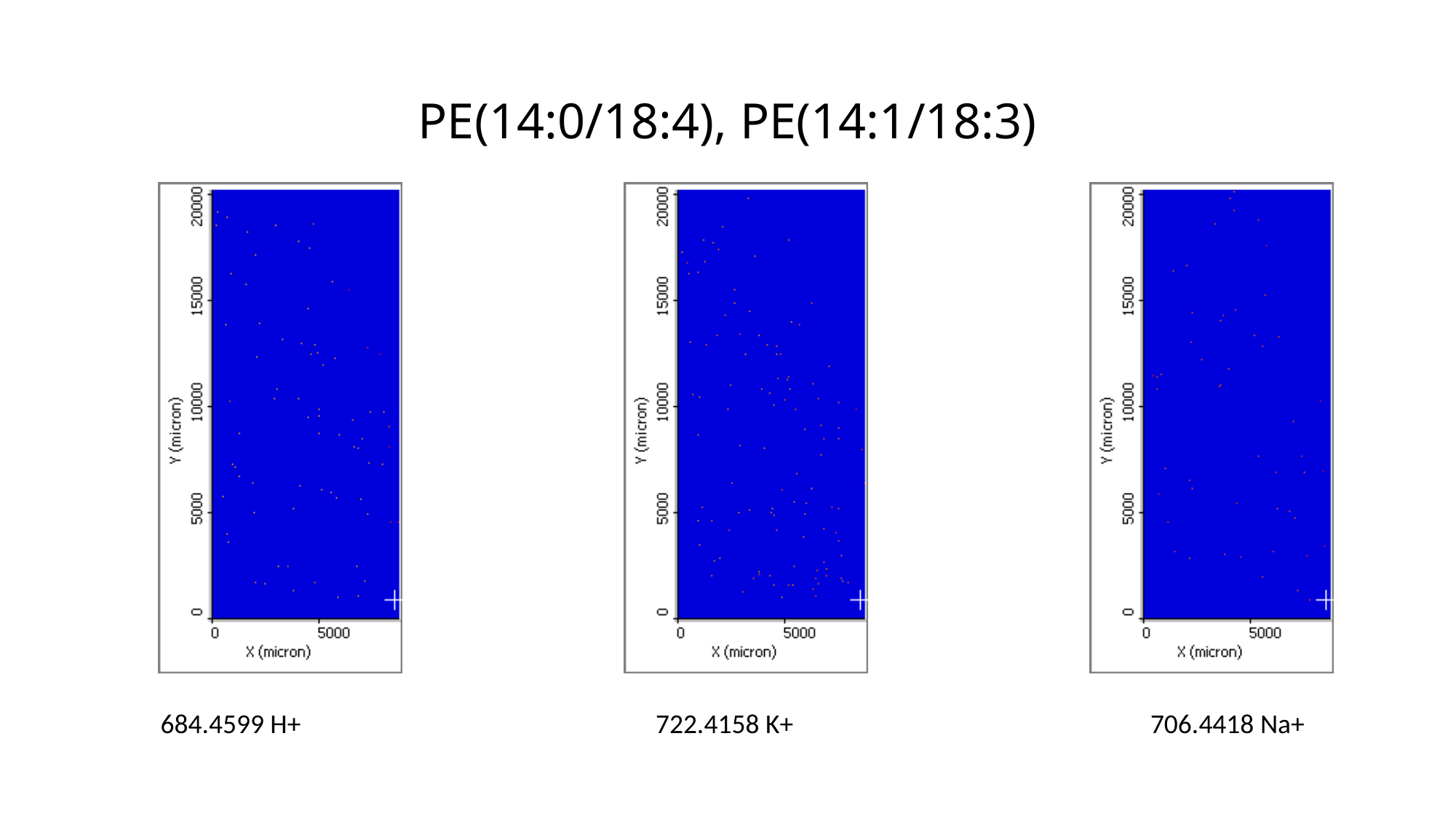

# PE(14:0/18:4), PE(14:1/18:3)
684.4599 H+
722.4158 K+
706.4418 Na+

### Slide 11
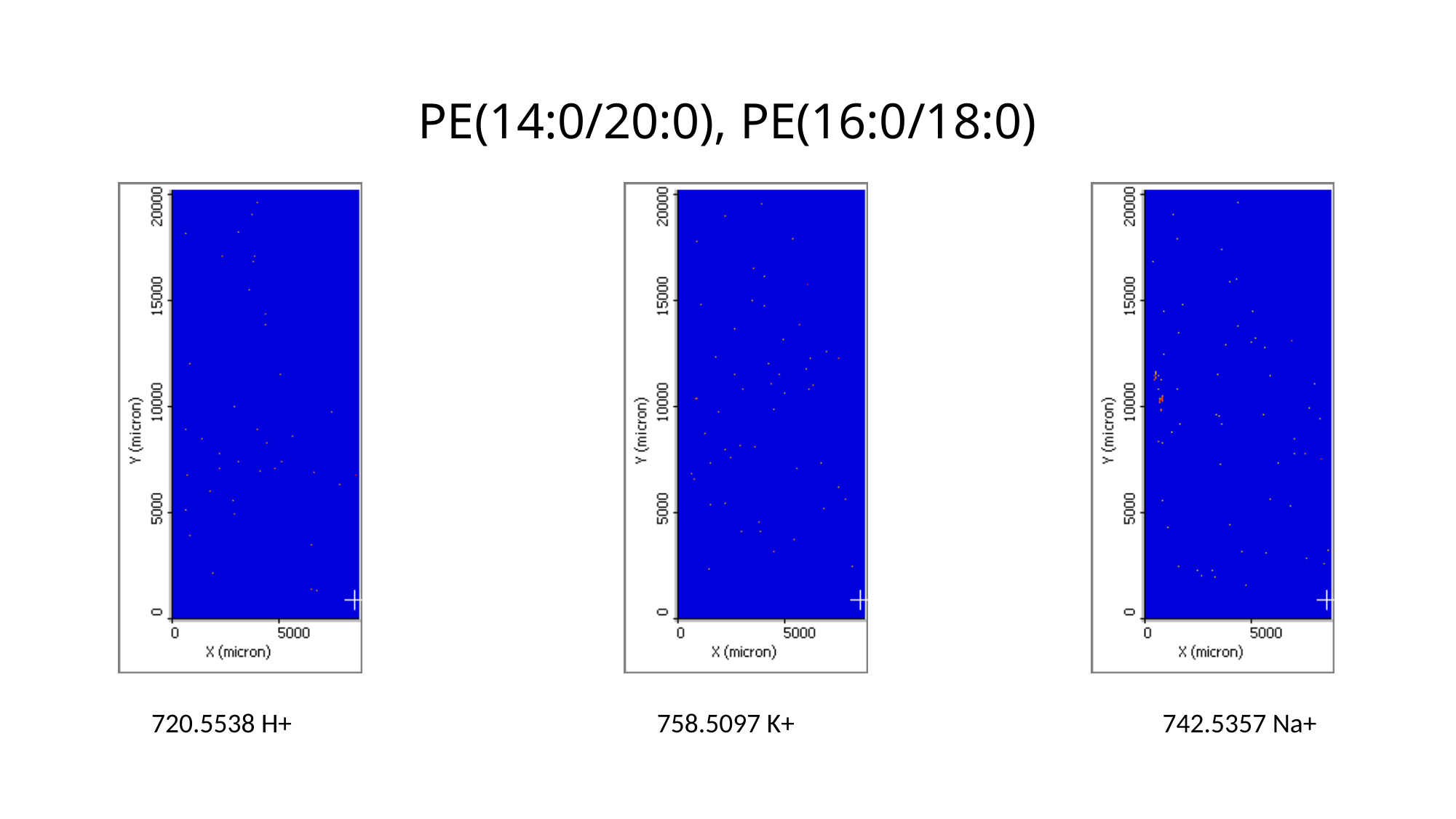

# PE(14:0/20:0), PE(16:0/18:0)
720.5538 H+
758.5097 K+
742.5357 Na+

### Slide 12
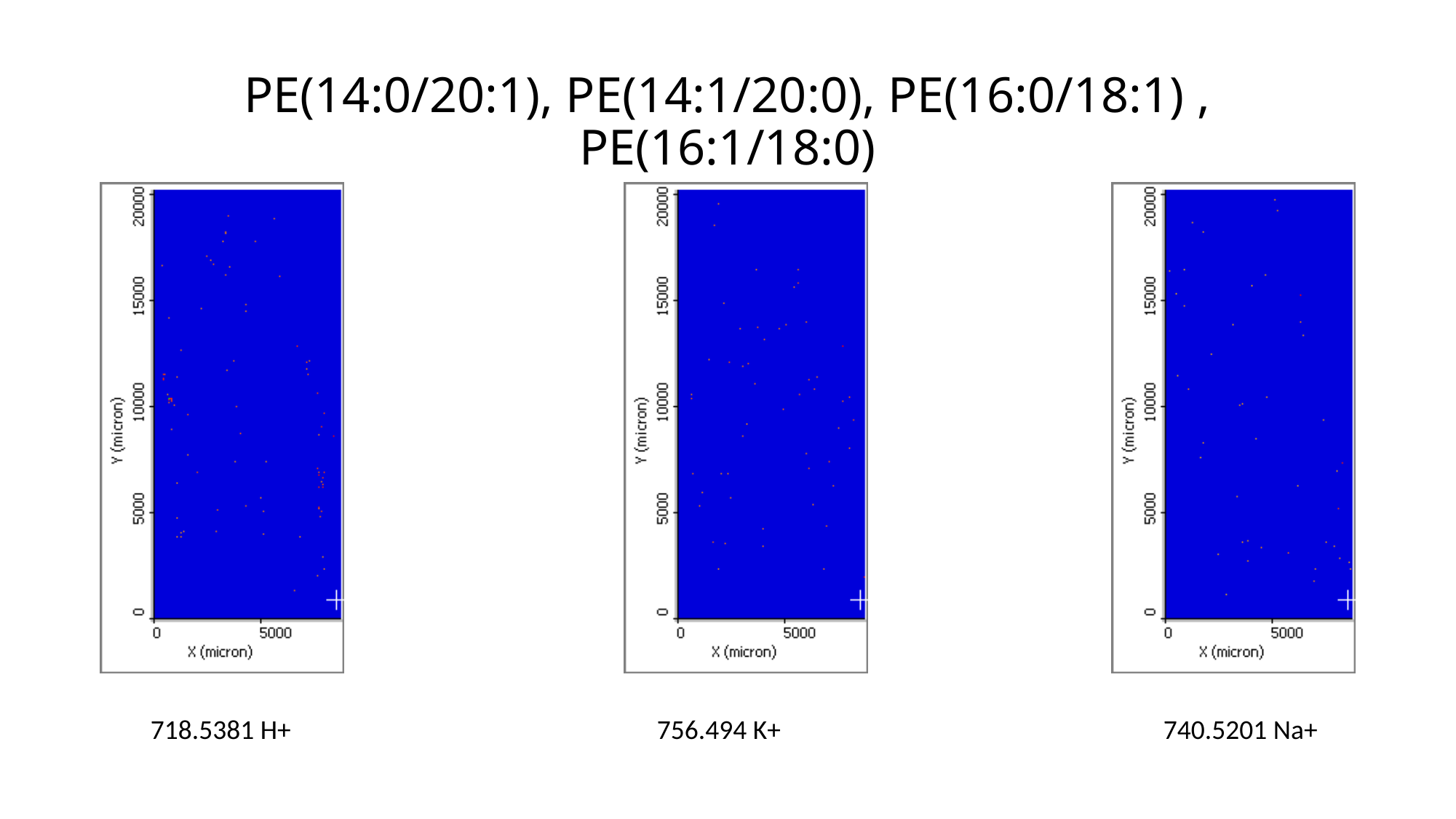

# PE(14:0/20:1), PE(14:1/20:0), PE(16:0/18:1) , PE(16:1/18:0)
718.5381 H+
756.494 K+
740.5201 Na+

### Slide 13
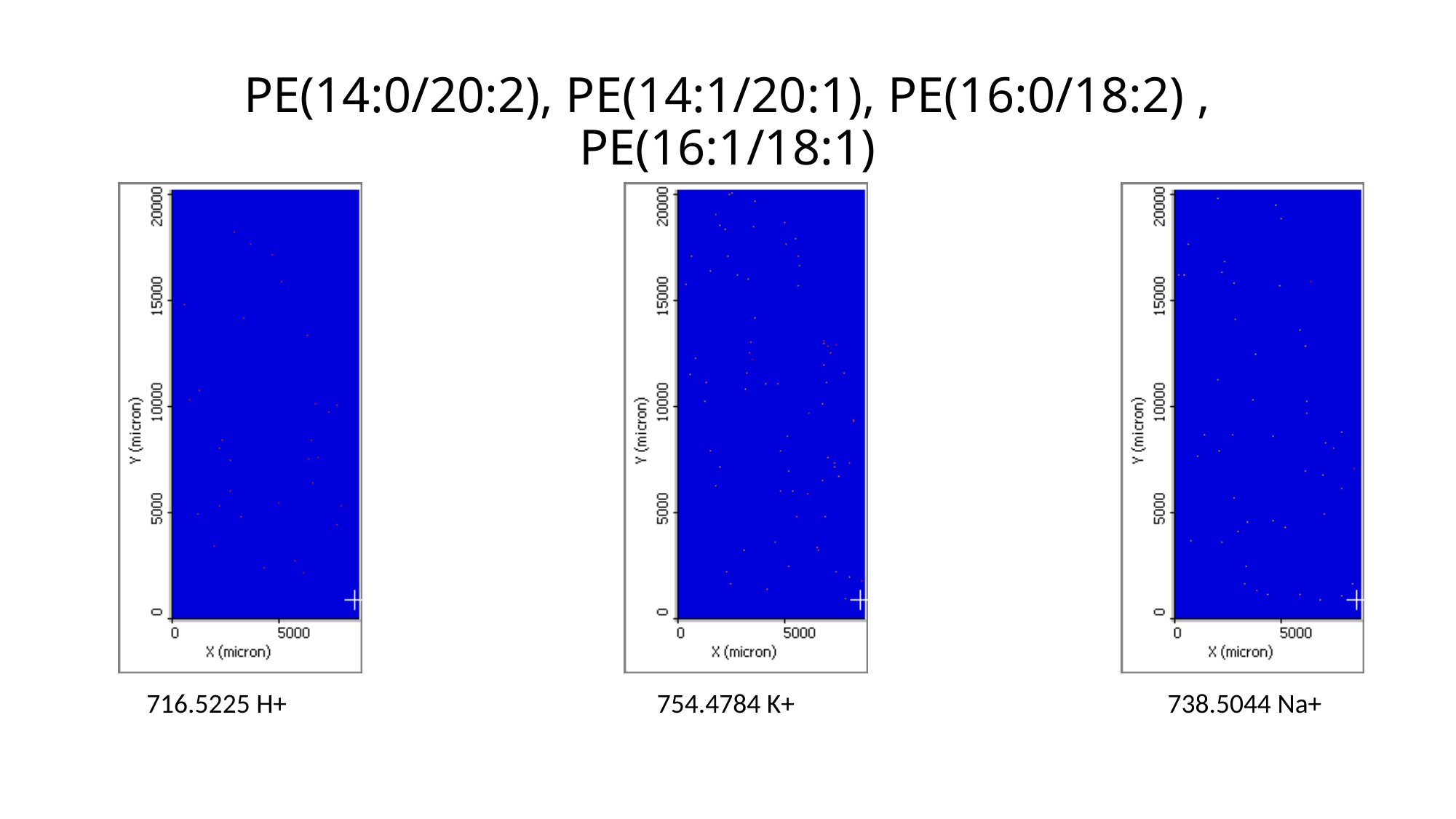

# PE(14:0/20:2), PE(14:1/20:1), PE(16:0/18:2) , PE(16:1/18:1)
716.5225 H+
754.4784 K+
738.5044 Na+

### Slide 14
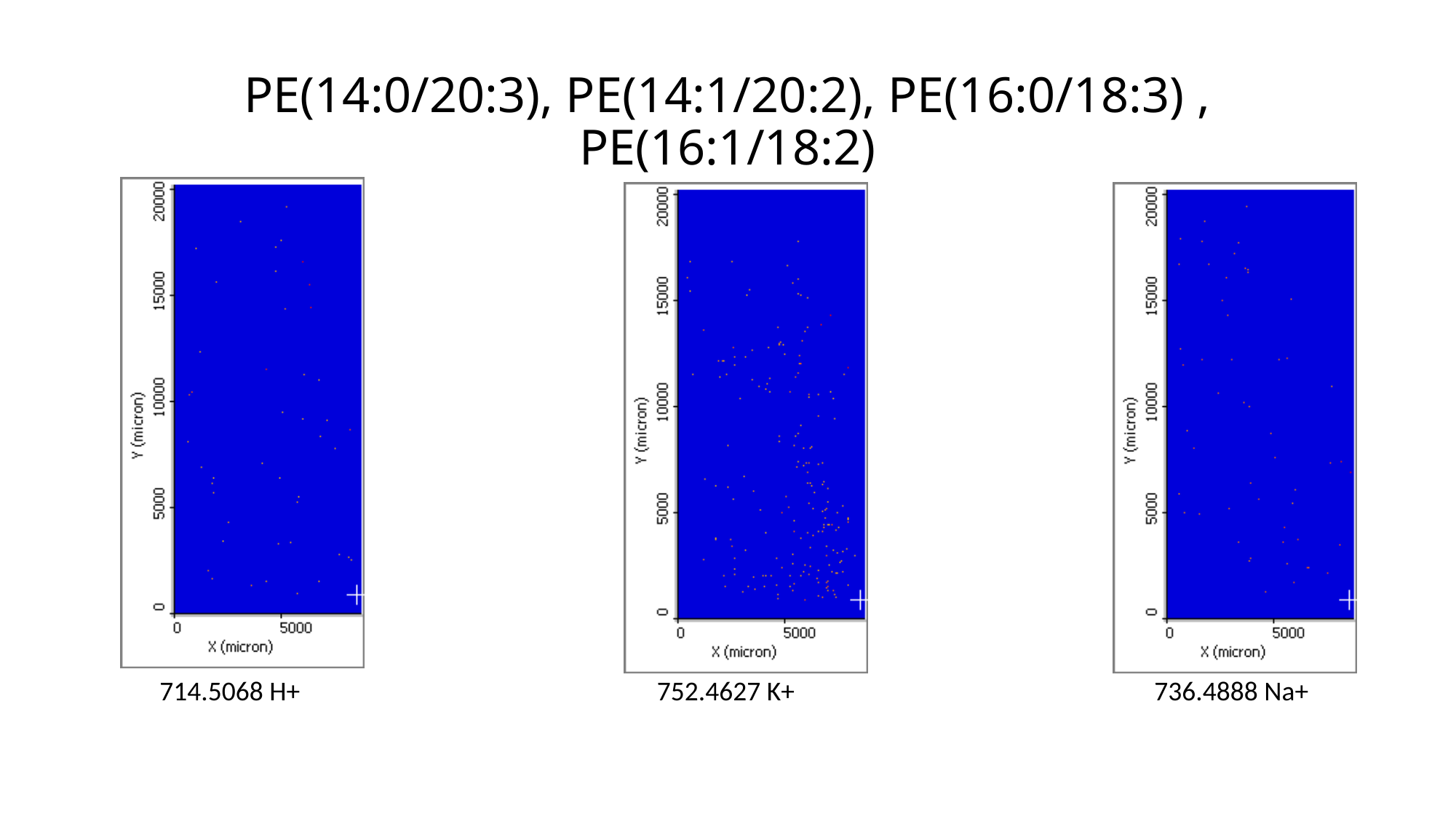

# PE(14:0/20:3), PE(14:1/20:2), PE(16:0/18:3) , PE(16:1/18:2)
714.5068 H+
752.4627 K+
736.4888 Na+

### Slide 15
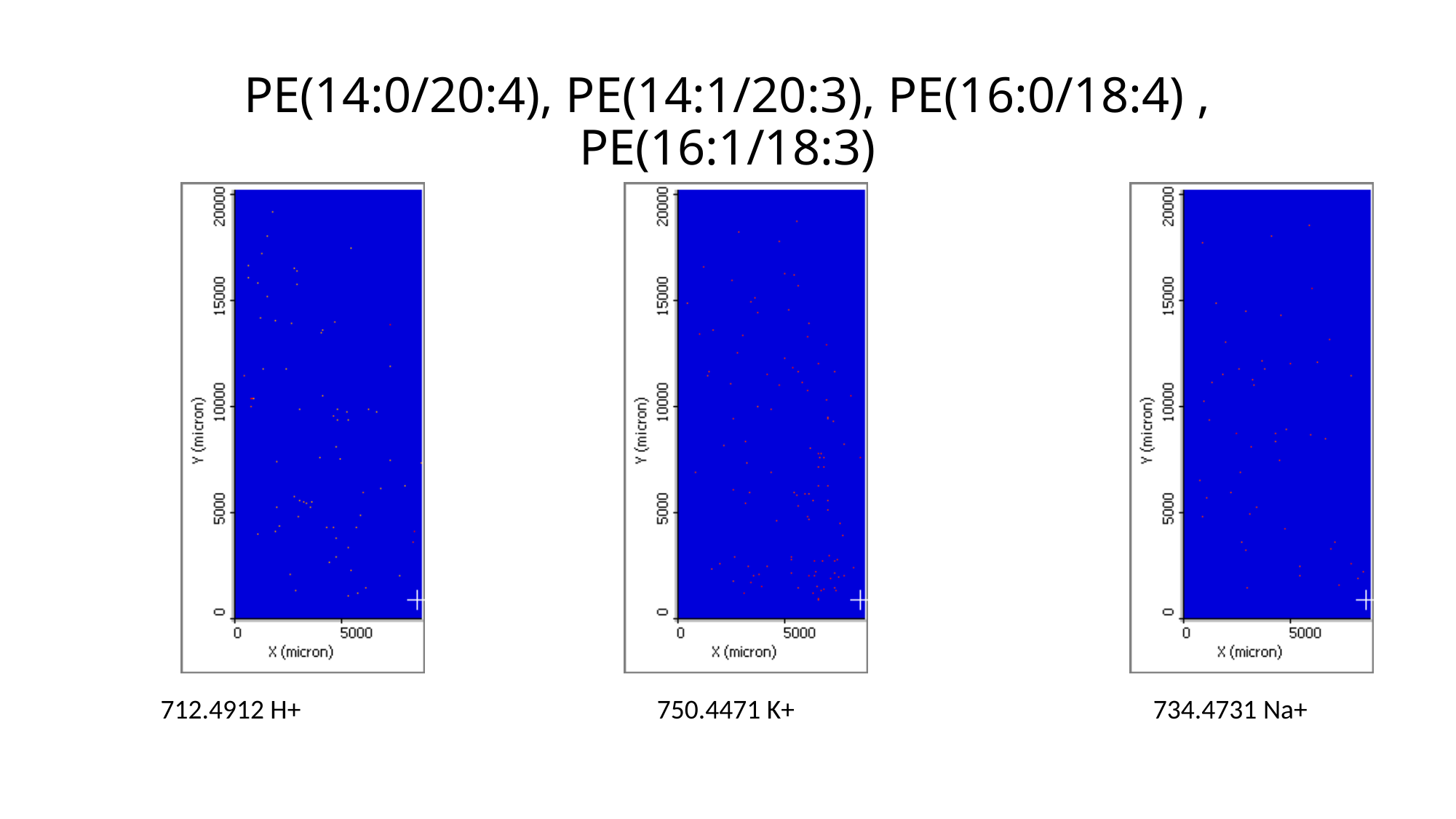

# PE(14:0/20:4), PE(14:1/20:3), PE(16:0/18:4) , PE(16:1/18:3)
712.4912 H+
750.4471 K+
734.4731 Na+

### Slide 16
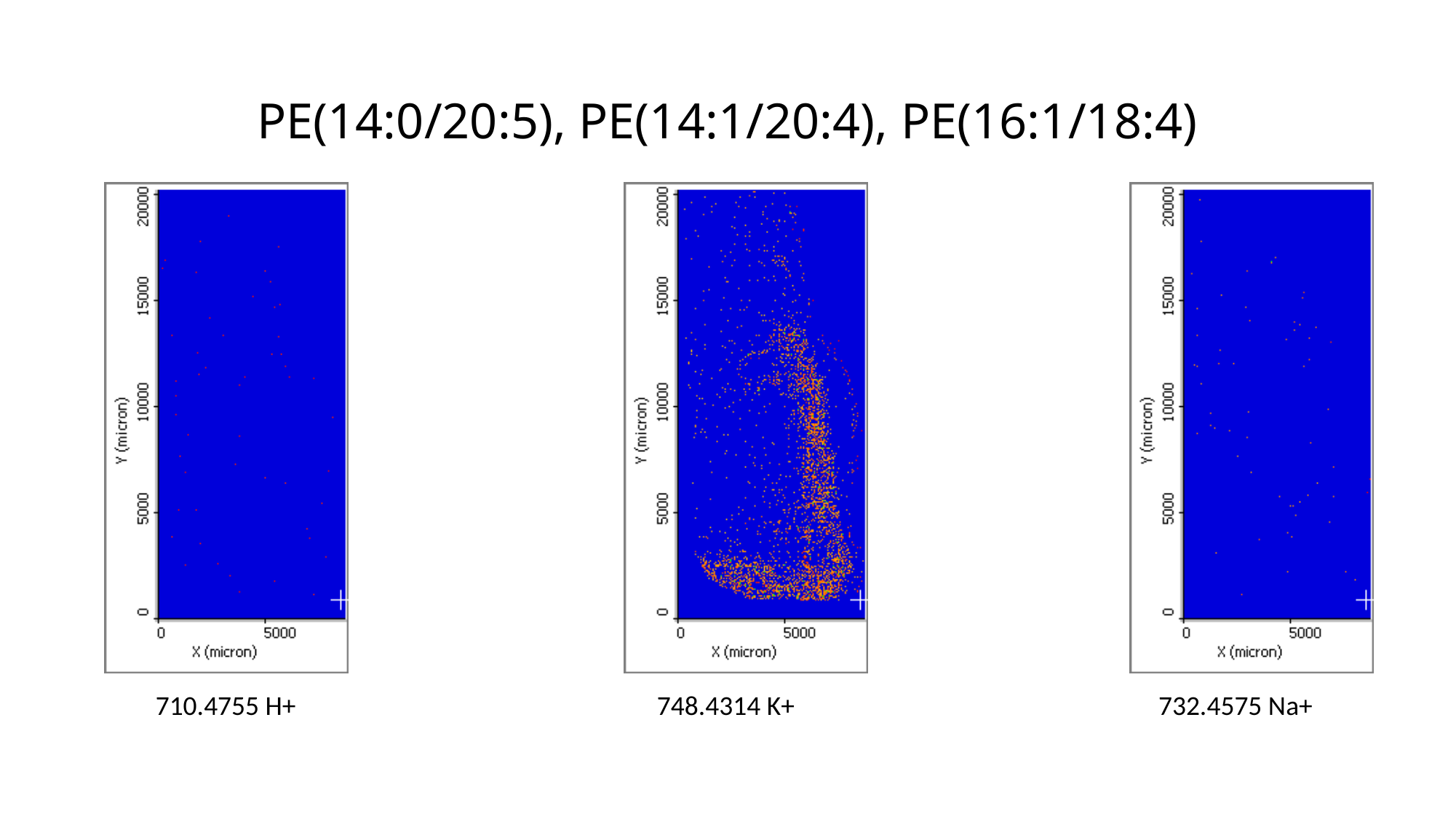

# PE(14:0/20:5), PE(14:1/20:4), PE(16:1/18:4)
710.4755 H+
748.4314 K+
732.4575 Na+

### Slide 17
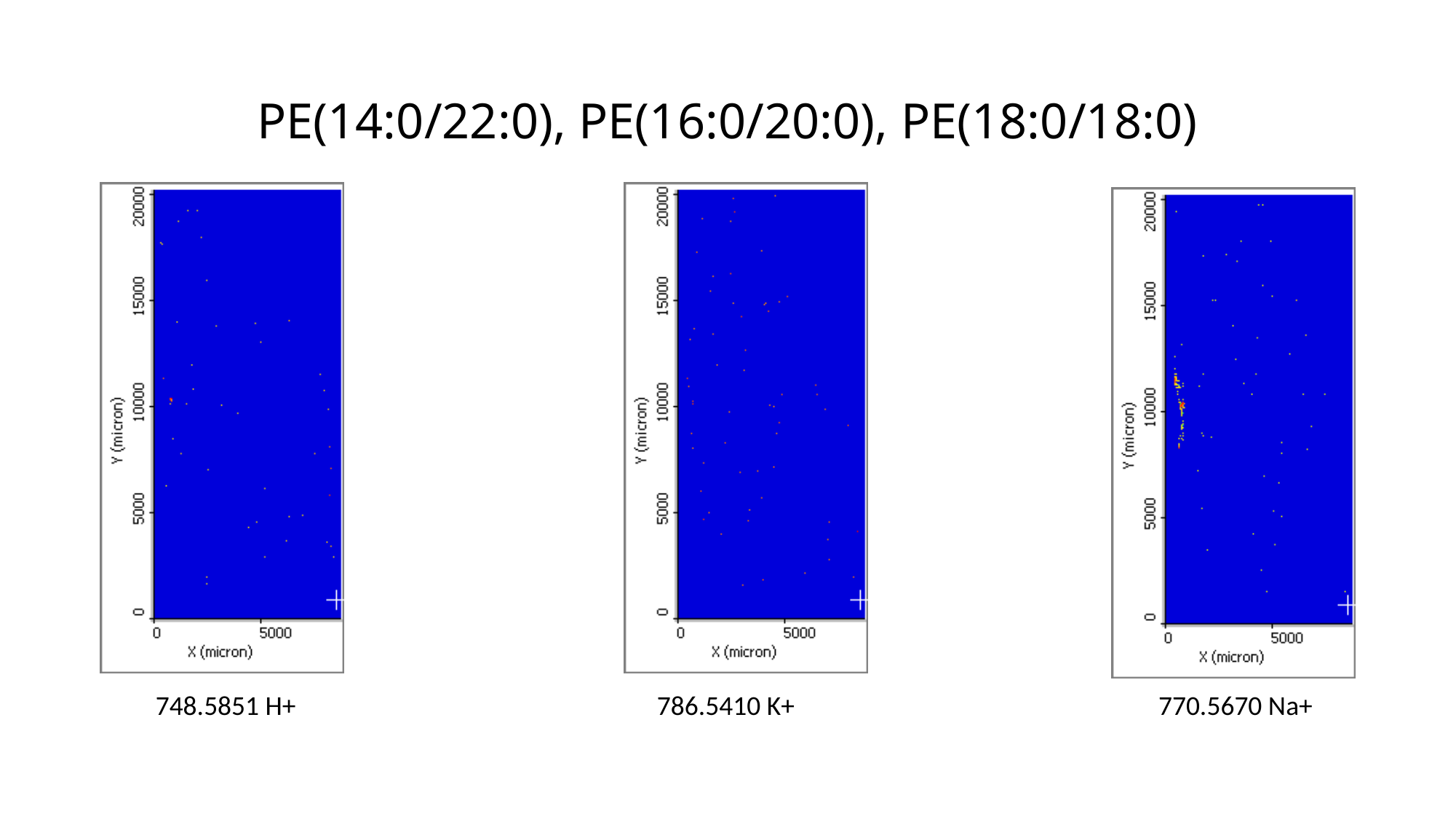

# PE(14:0/22:0), PE(16:0/20:0), PE(18:0/18:0)
748.5851 H+
786.5410 K+
770.5670 Na+

### Slide 18
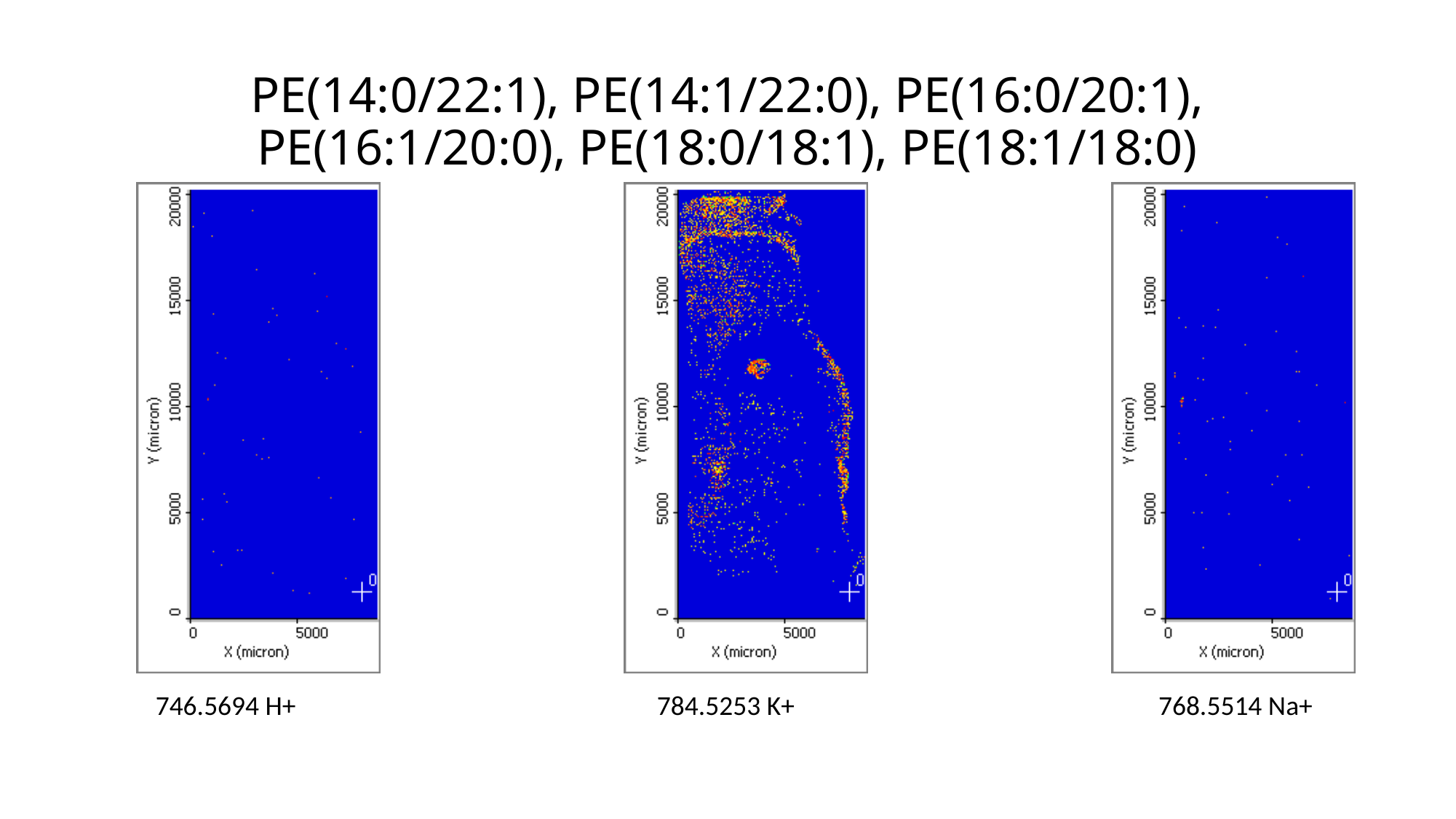

# PE(14:0/22:1), PE(14:1/22:0), PE(16:0/20:1), PE(16:1/20:0), PE(18:0/18:1), PE(18:1/18:0)
746.5694 H+
784.5253 K+
768.5514 Na+

### Slide 19
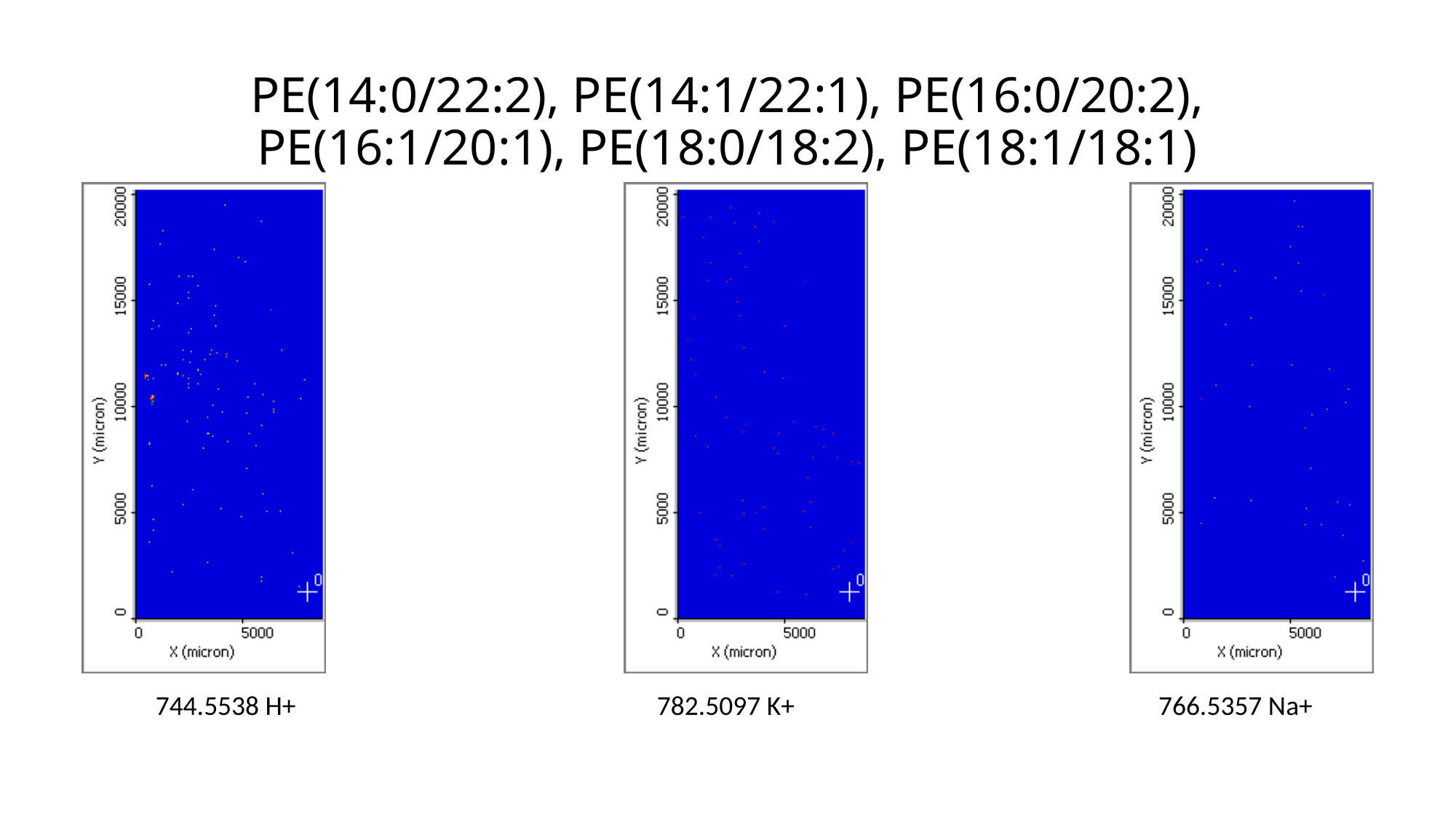

# PE(14:0/22:2), PE(14:1/22:1), PE(16:0/20:2), PE(16:1/20:1), PE(18:0/18:2), PE(18:1/18:1)
744.5538 H+
782.5097 K+
766.5357 Na+

### Slide 20
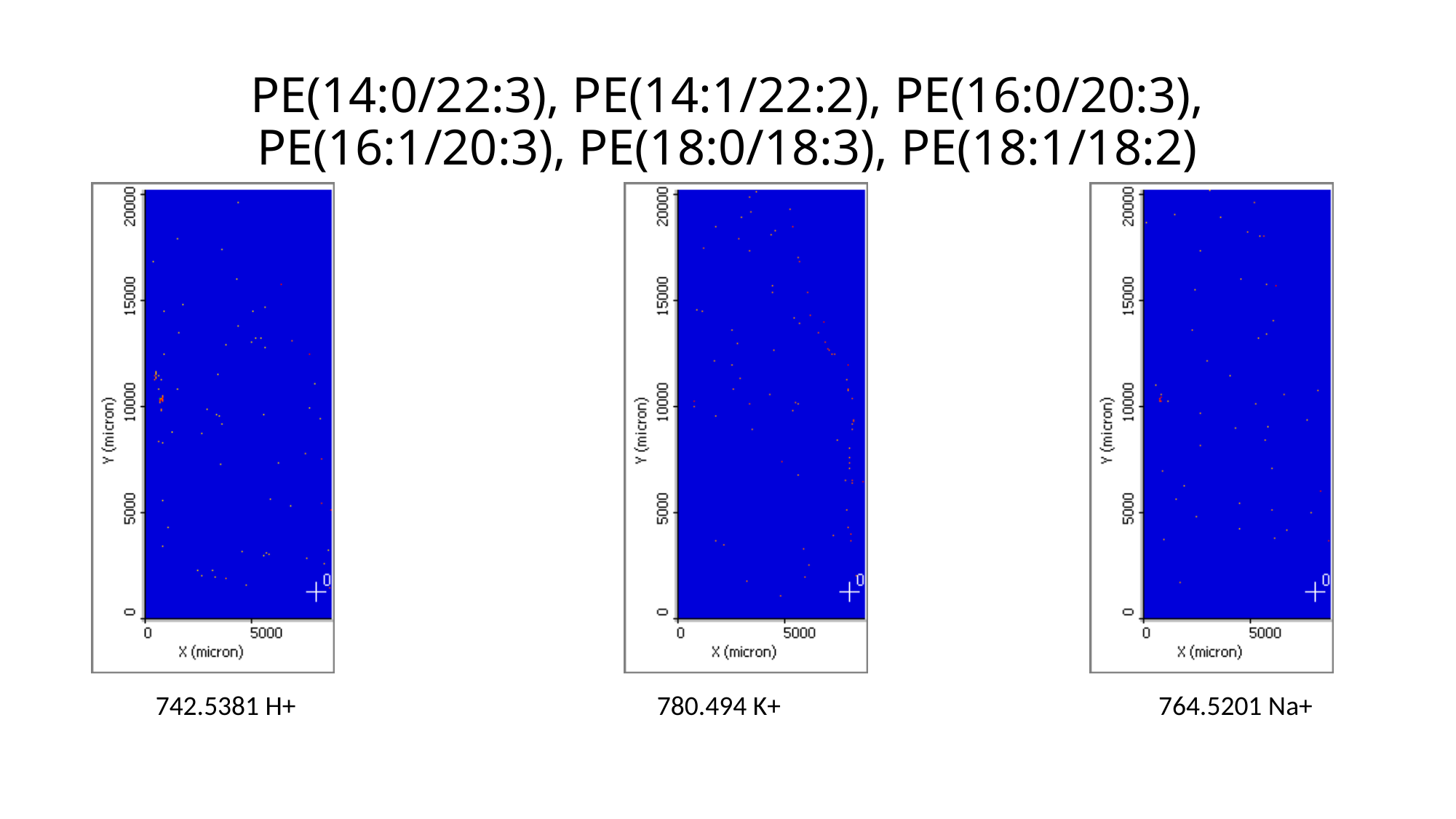

# PE(14:0/22:3), PE(14:1/22:2), PE(16:0/20:3), PE(16:1/20:3), PE(18:0/18:3), PE(18:1/18:2)
742.5381 H+
780.494 K+
764.5201 Na+

### Slide 21
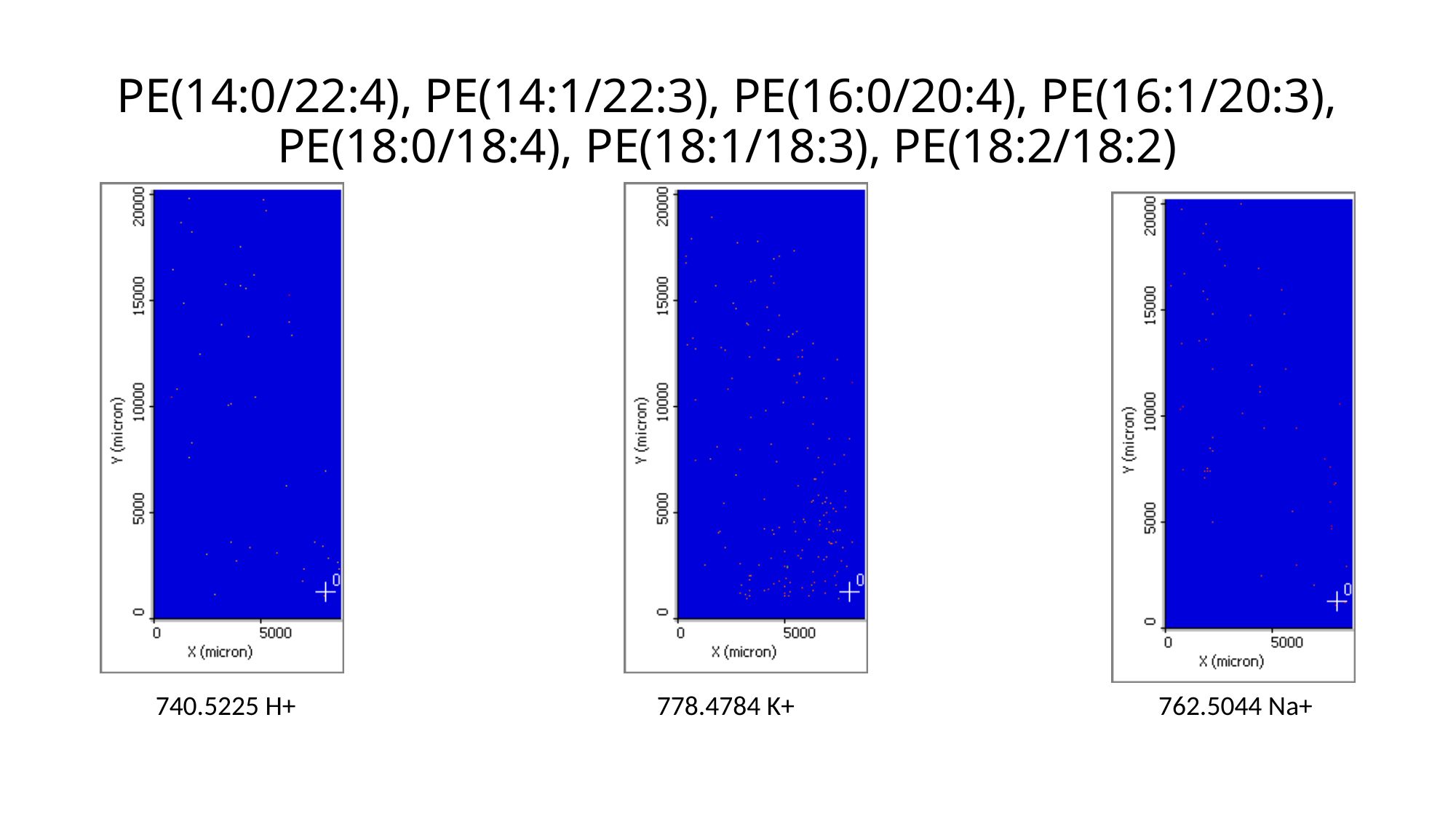

# PE(14:0/22:4), PE(14:1/22:3), PE(16:0/20:4), PE(16:1/20:3), PE(18:0/18:4), PE(18:1/18:3), PE(18:2/18:2)
740.5225 H+
778.4784 K+
762.5044 Na+

### Slide 22
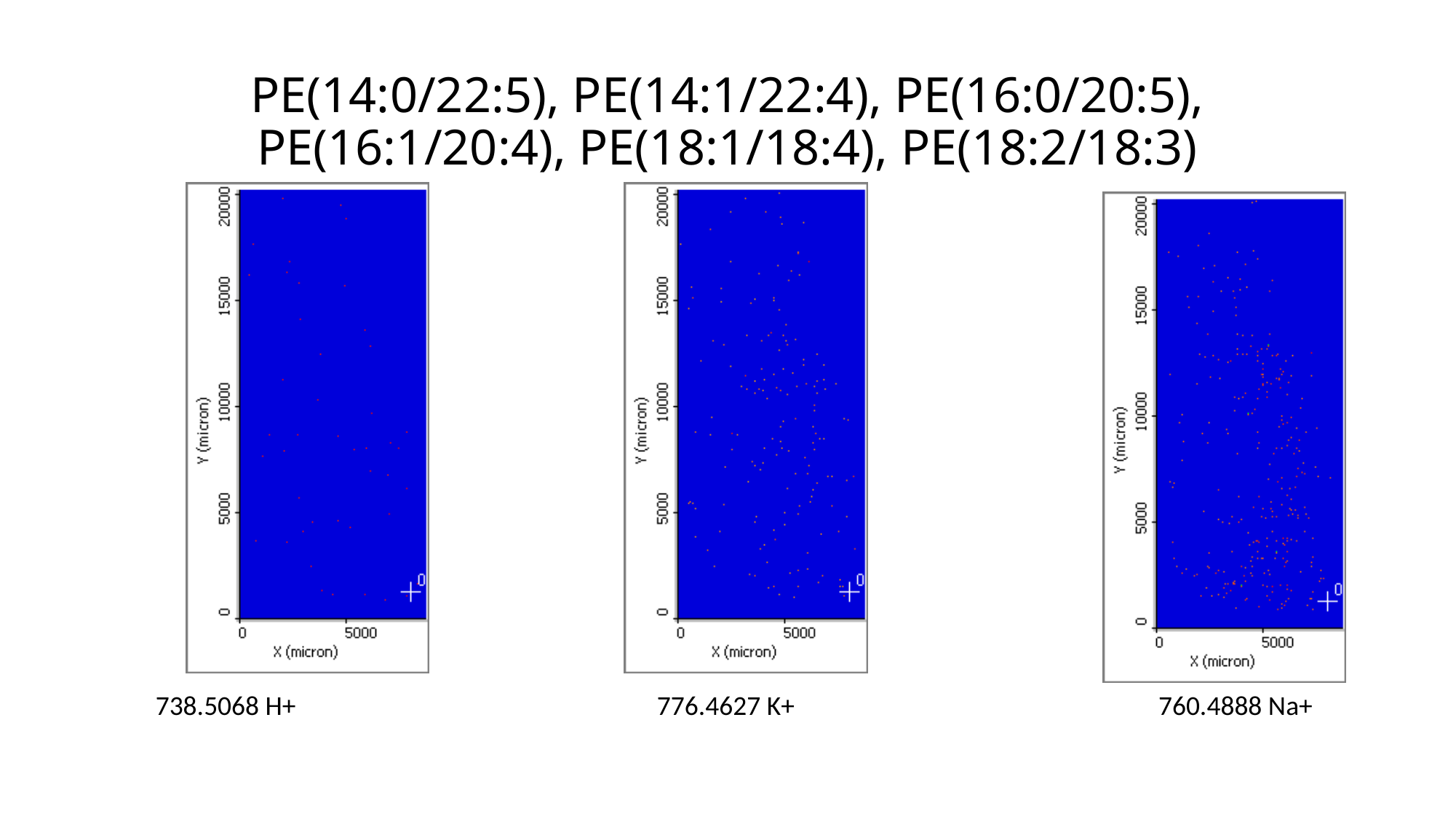

# PE(14:0/22:5), PE(14:1/22:4), PE(16:0/20:5), PE(16:1/20:4), PE(18:1/18:4), PE(18:2/18:3)
738.5068 H+
776.4627 K+
760.4888 Na+

### Slide 23
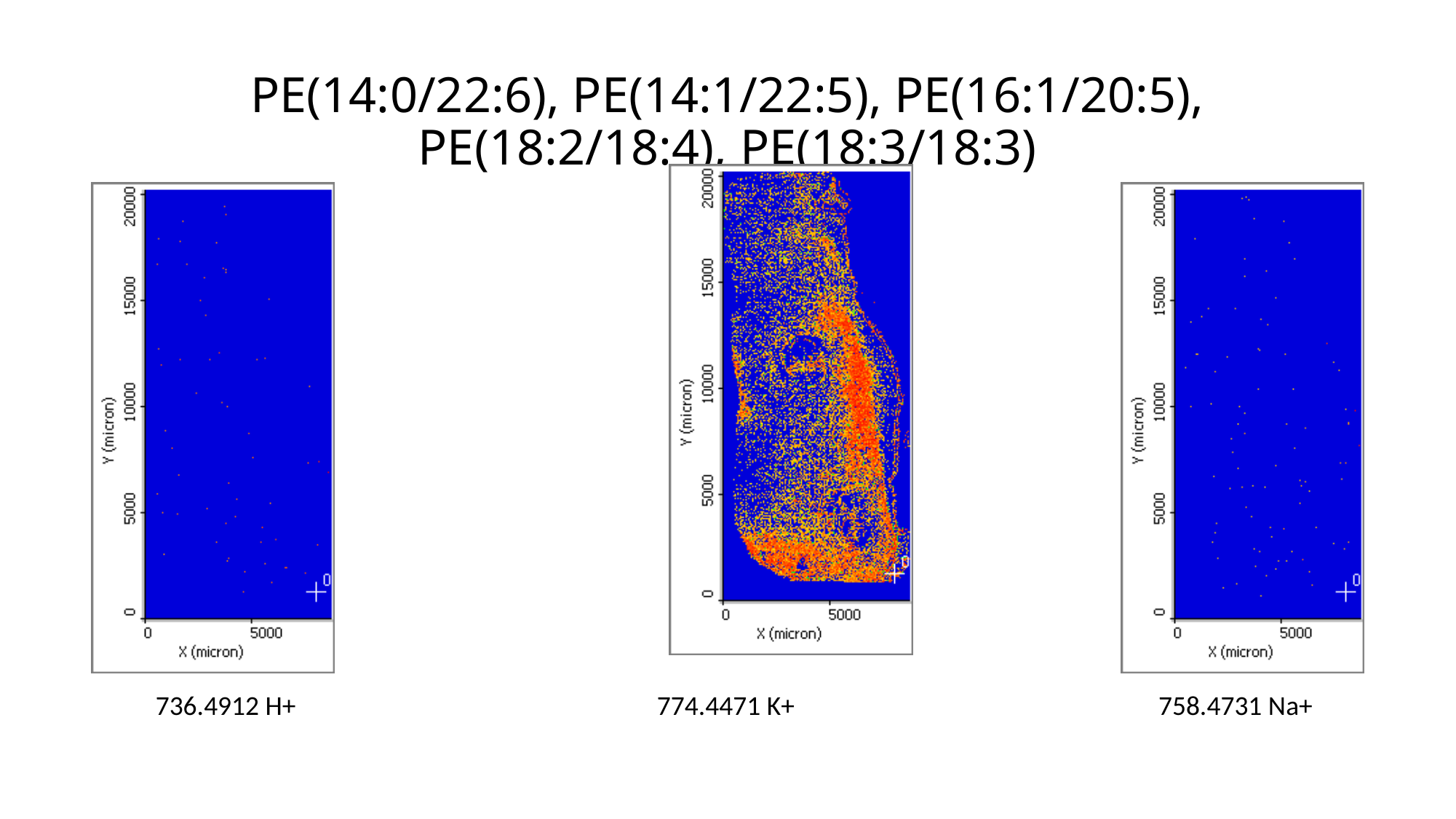

# PE(14:0/22:6), PE(14:1/22:5), PE(16:1/20:5), PE(18:2/18:4), PE(18:3/18:3)
736.4912 H+
774.4471 K+
758.4731 Na+

### Slide 24
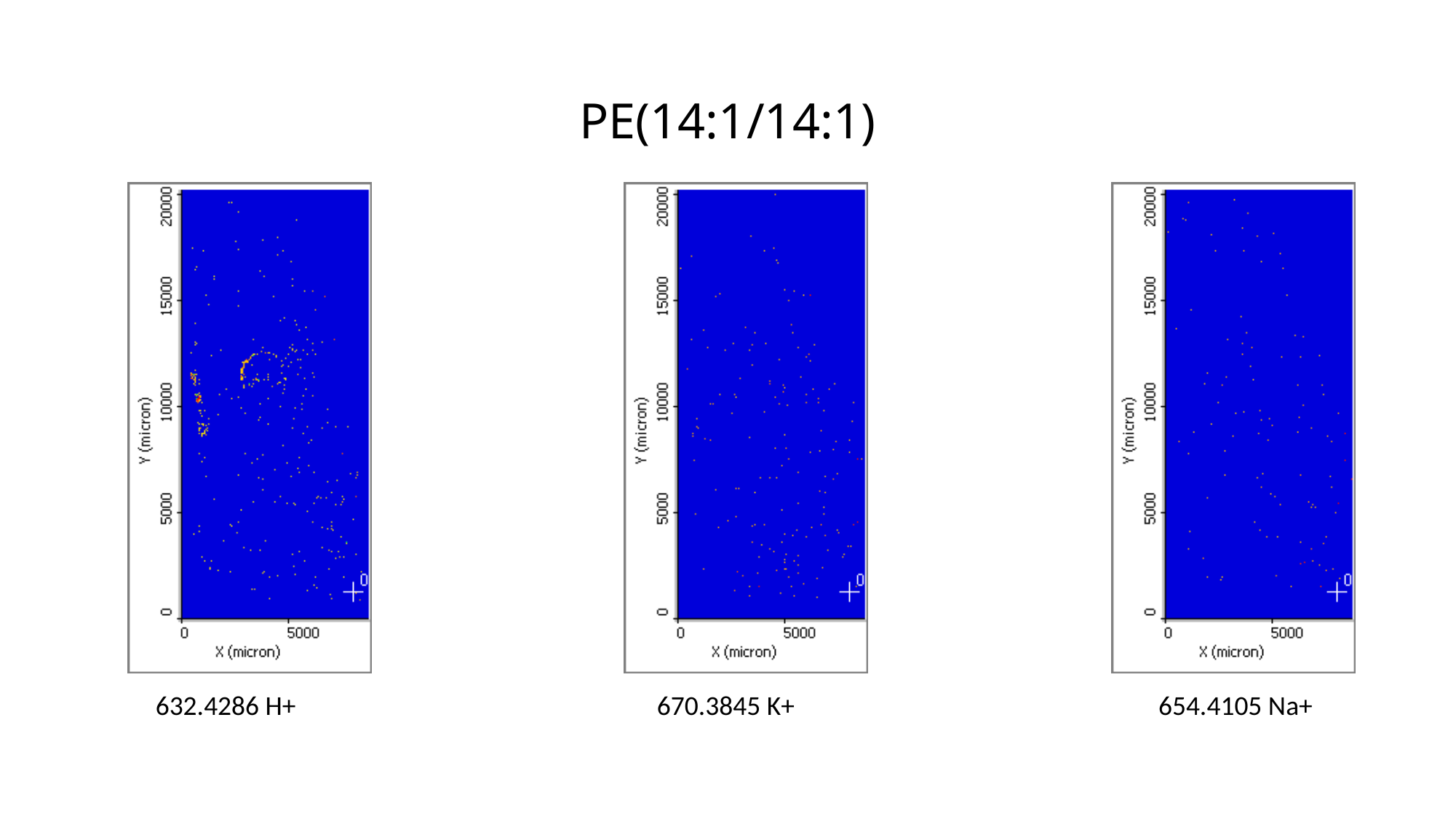

# PE(14:1/14:1)
632.4286 H+
670.3845 K+
654.4105 Na+

### Slide 25
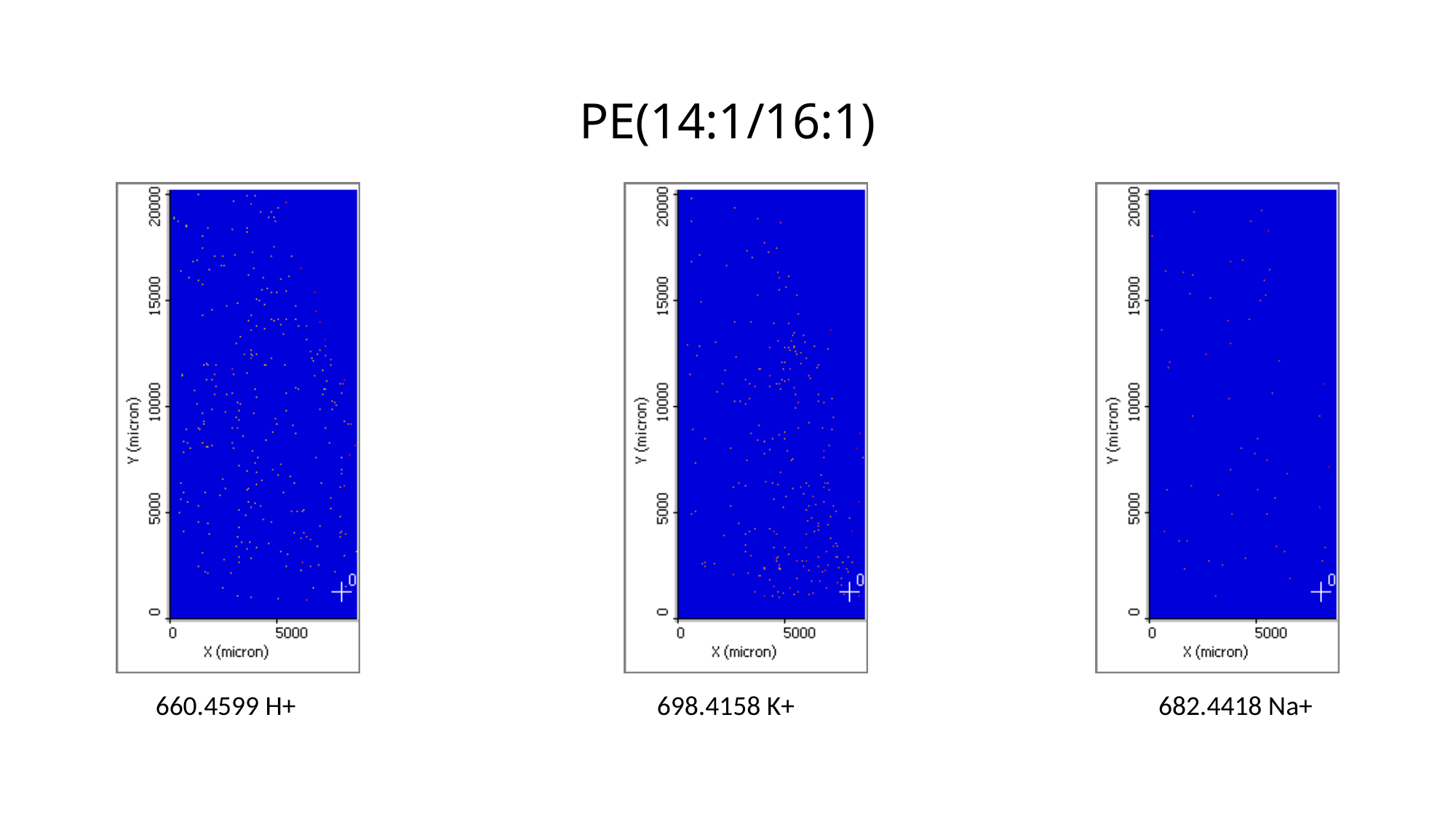

# PE(14:1/16:1)
660.4599 H+
698.4158 K+
682.4418 Na+

### Slide 26
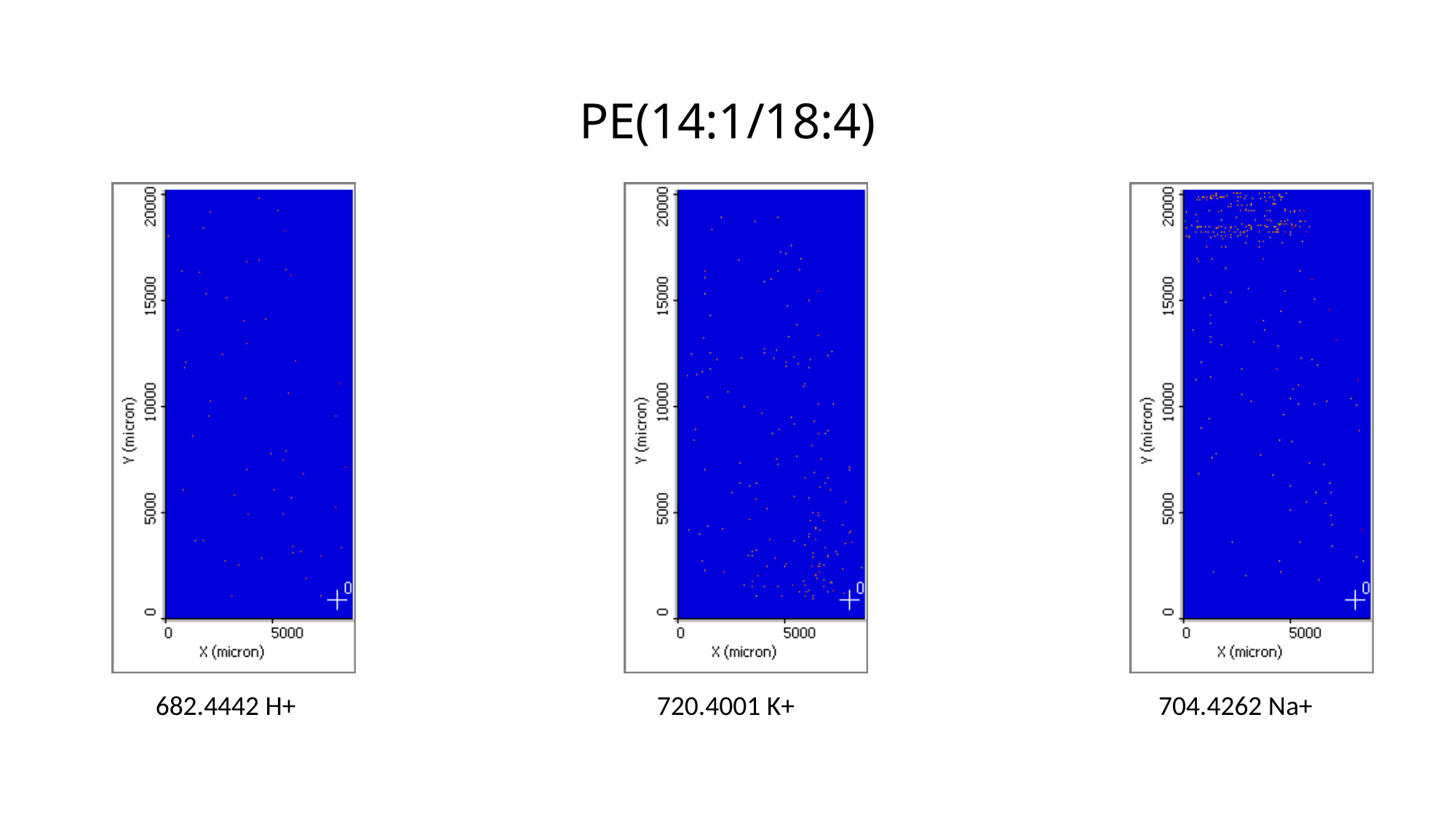

# PE(14:1/18:4)
682.4442 H+
720.4001 K+
704.4262 Na+

### Slide 27
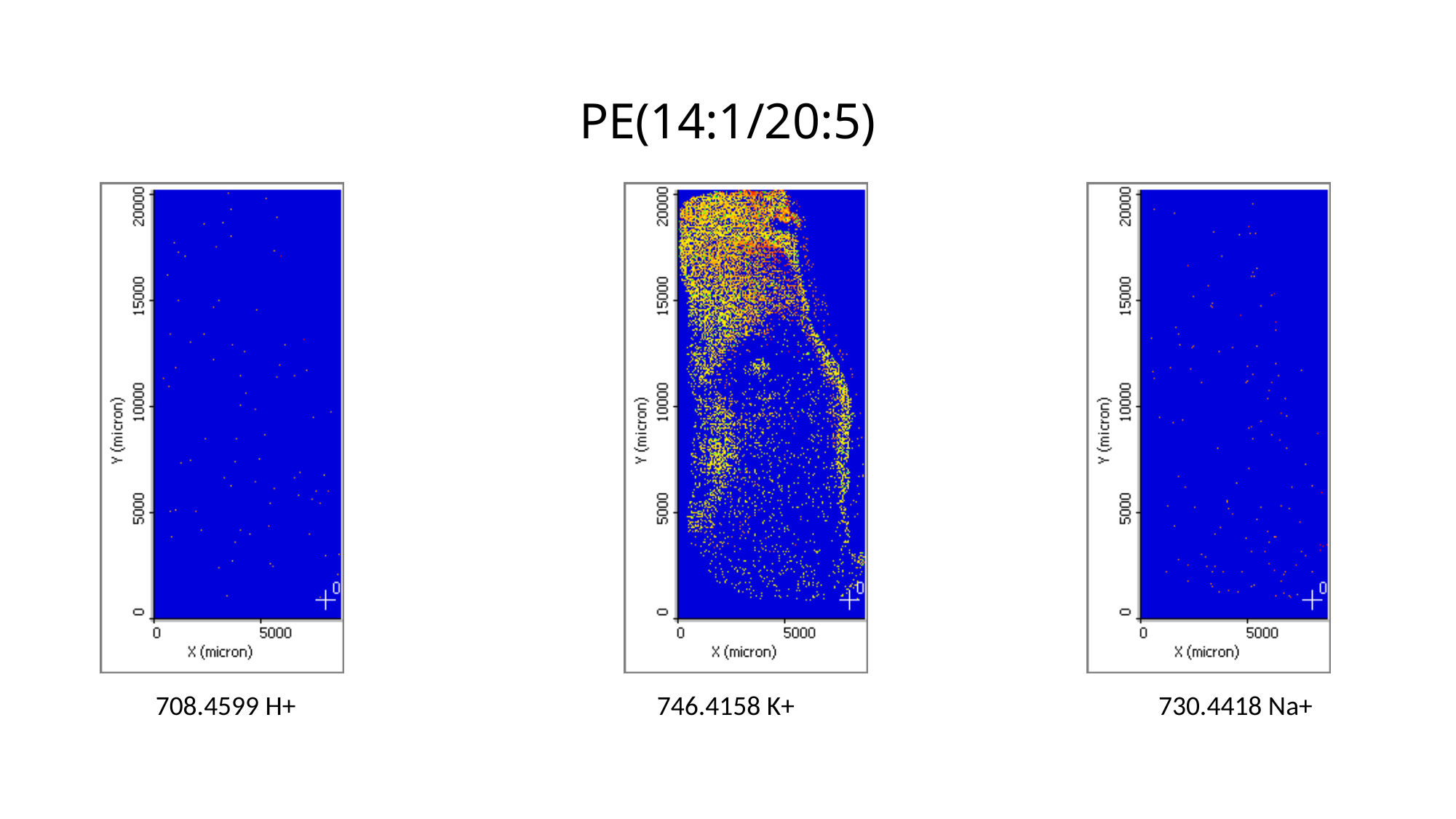

# PE(14:1/20:5)
708.4599 H+
746.4158 K+
730.4418 Na+

### Slide 28
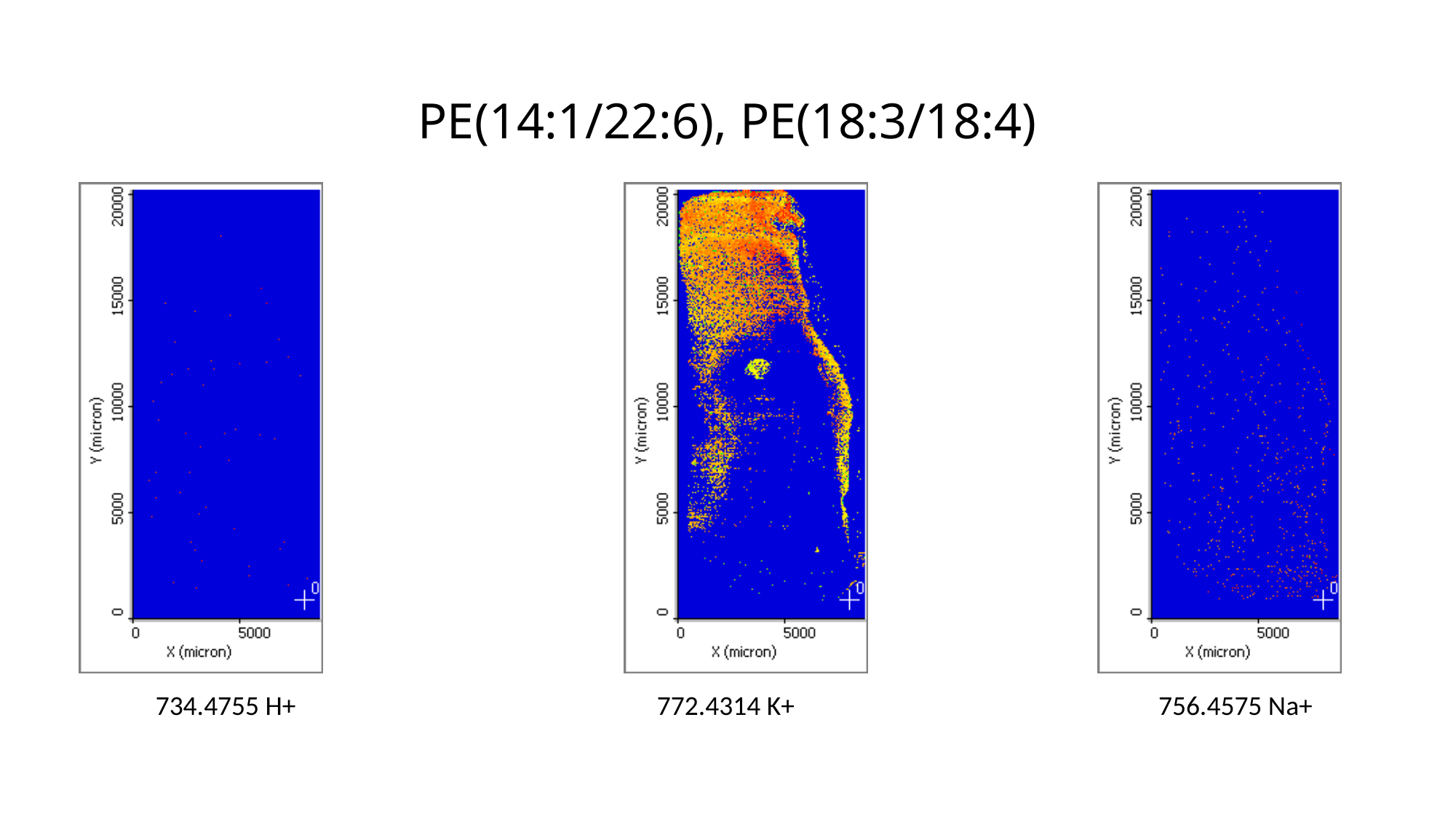

# PE(14:1/22:6), PE(18:3/18:4)
734.4755 H+
772.4314 K+
756.4575 Na+

### Slide 29
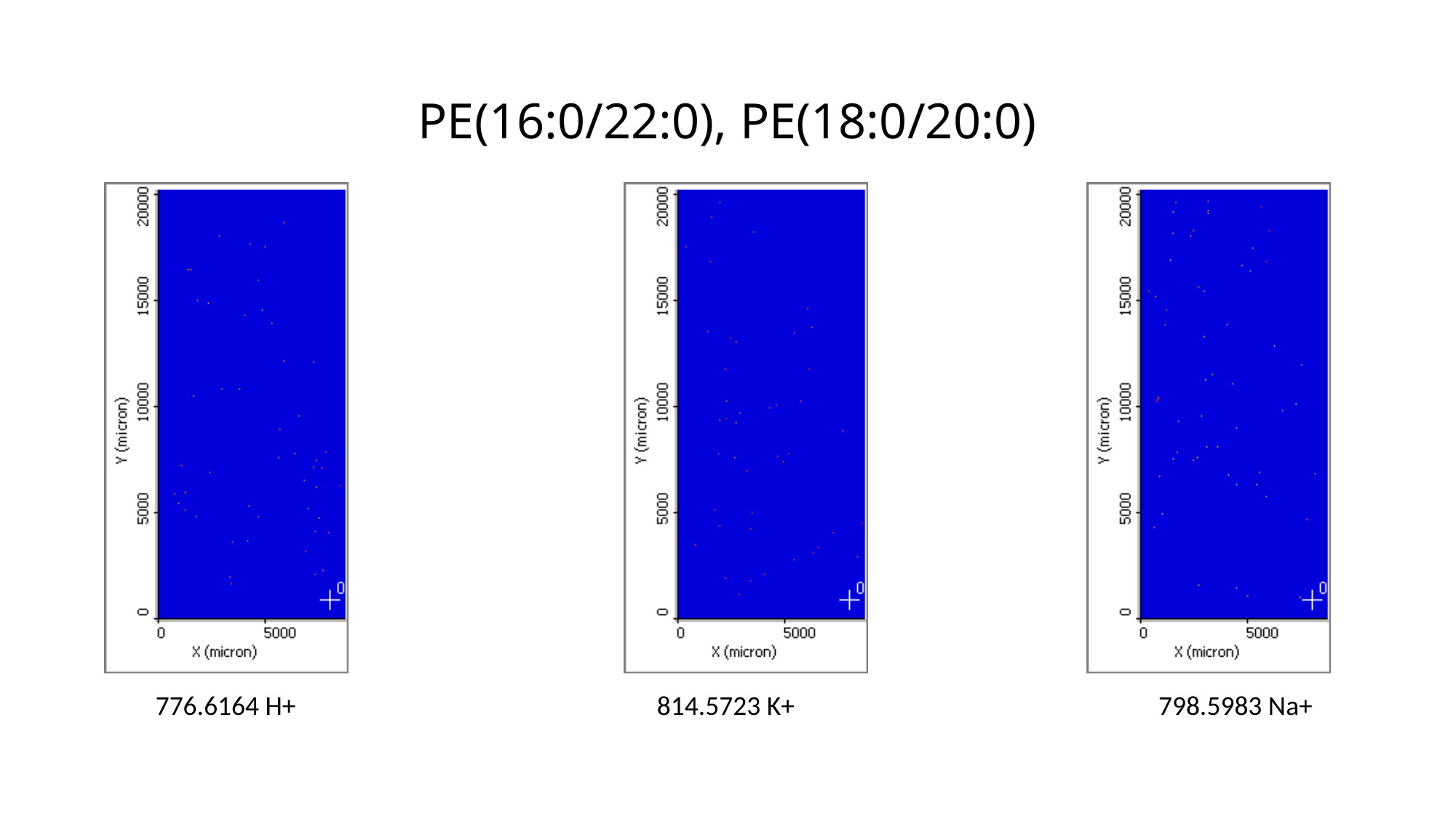

# PE(16:0/22:0), PE(18:0/20:0)
776.6164 H+
814.5723 K+
798.5983 Na+

### Slide 30
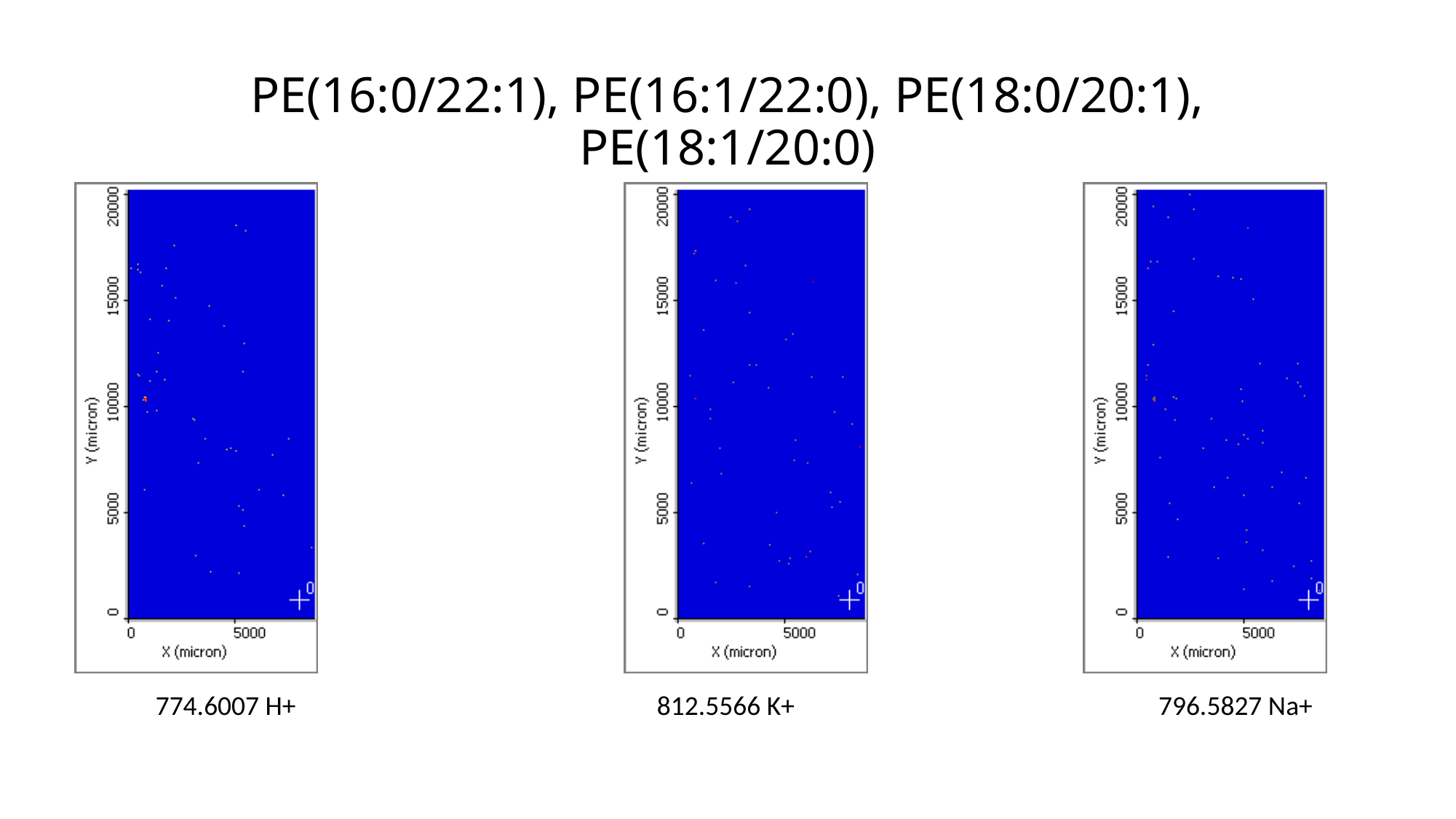

# PE(16:0/22:1), PE(16:1/22:0), PE(18:0/20:1), PE(18:1/20:0)
774.6007 H+
812.5566 K+
796.5827 Na+

### Slide 31
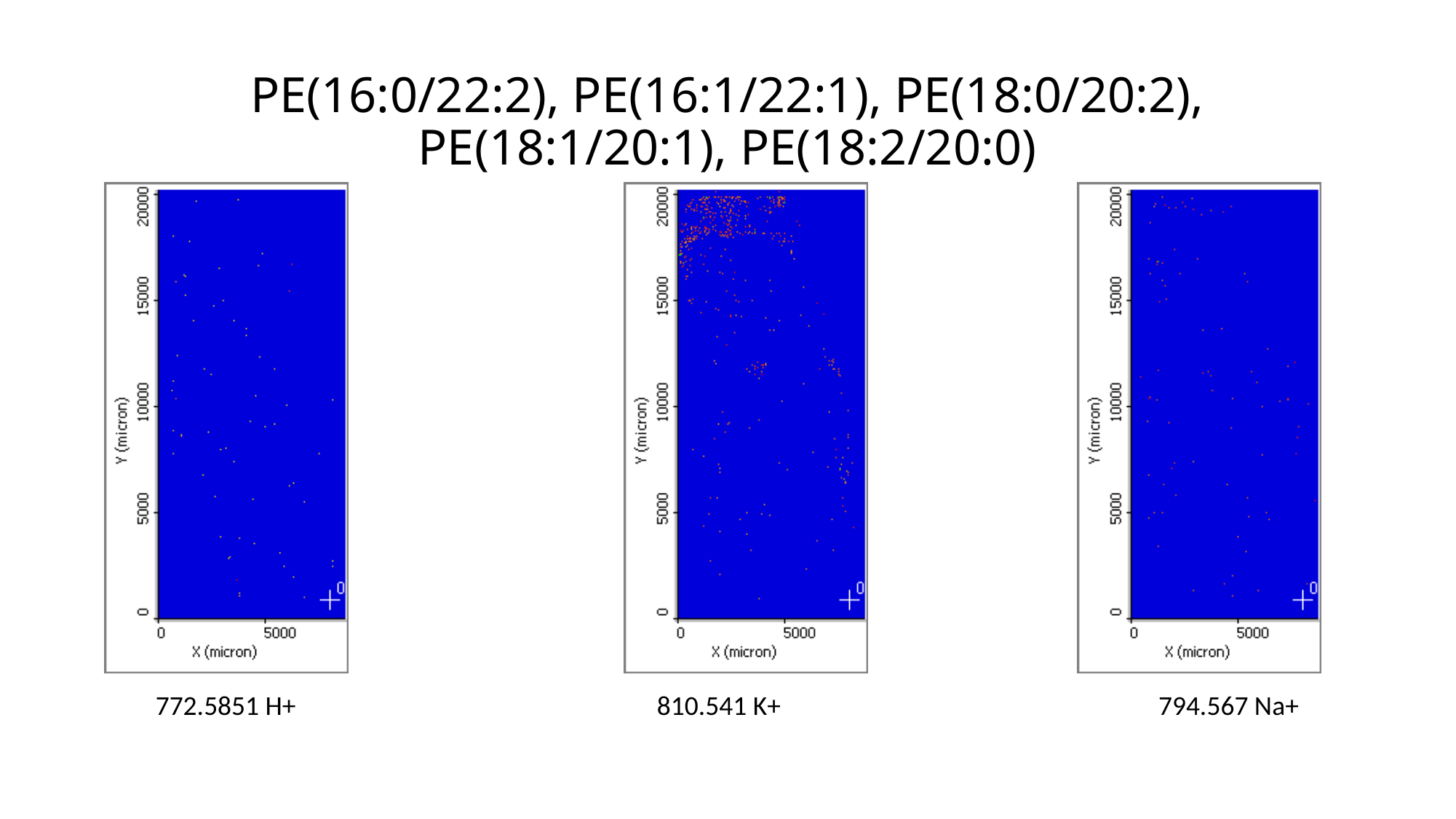

# PE(16:0/22:2), PE(16:1/22:1), PE(18:0/20:2), PE(18:1/20:1), PE(18:2/20:0)
772.5851 H+
810.541 K+
794.567 Na+

### Slide 32
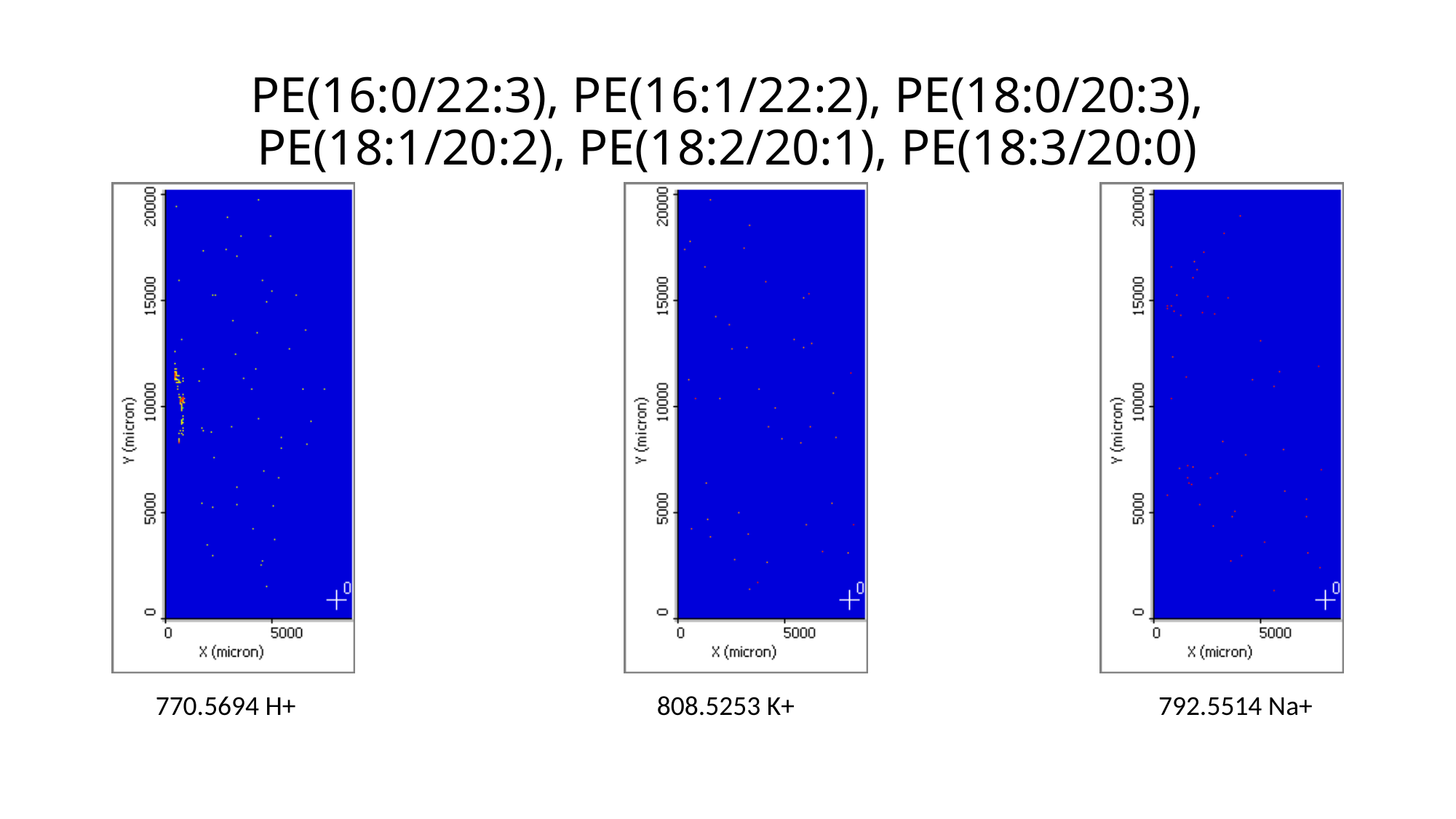

# PE(16:0/22:3), PE(16:1/22:2), PE(18:0/20:3), PE(18:1/20:2), PE(18:2/20:1), PE(18:3/20:0)
770.5694 H+
808.5253 K+
792.5514 Na+

### Slide 33
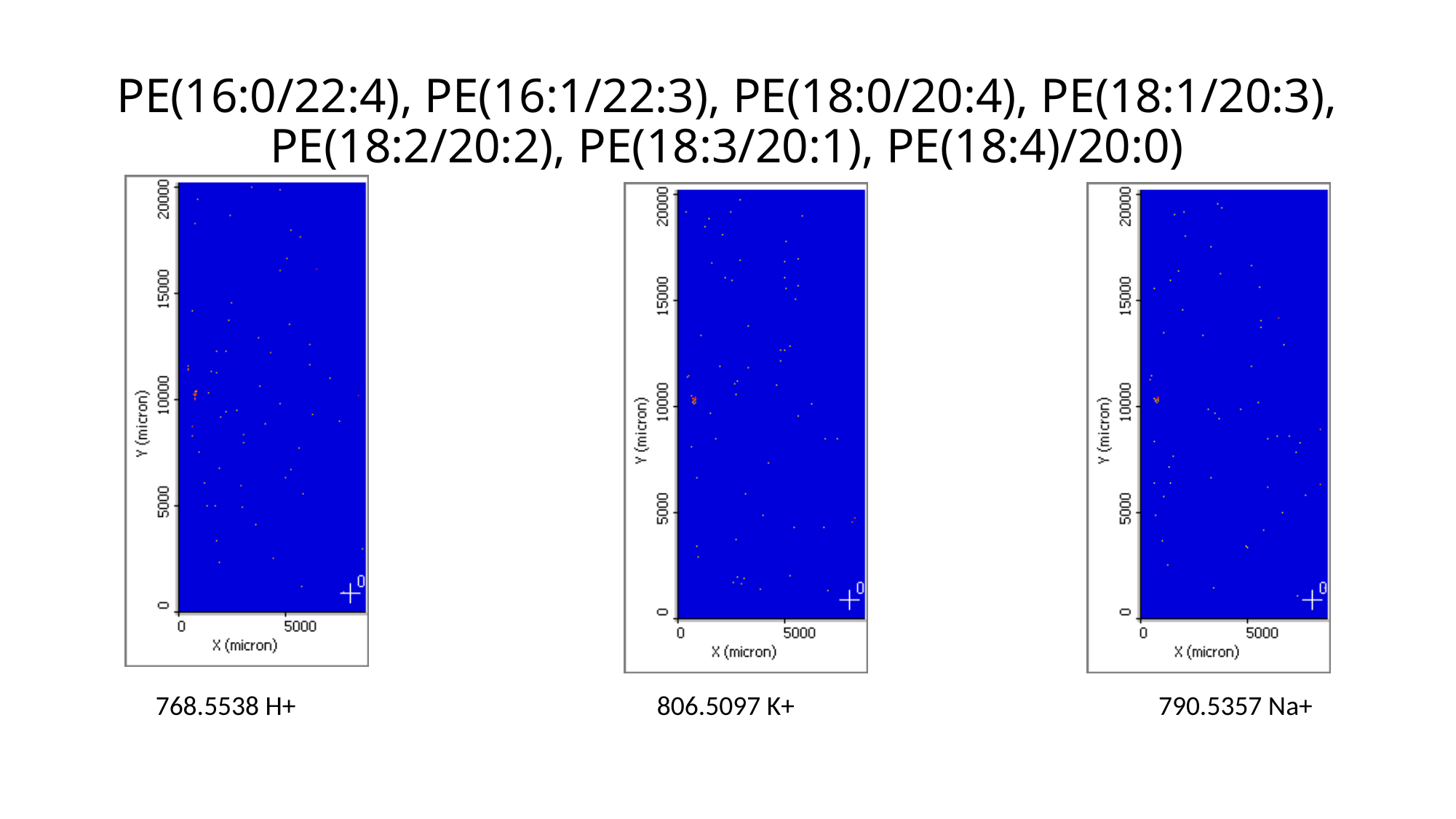

# PE(16:0/22:4), PE(16:1/22:3), PE(18:0/20:4), PE(18:1/20:3), PE(18:2/20:2), PE(18:3/20:1), PE(18:4)/20:0)
768.5538 H+
806.5097 K+
790.5357 Na+

### Slide 34
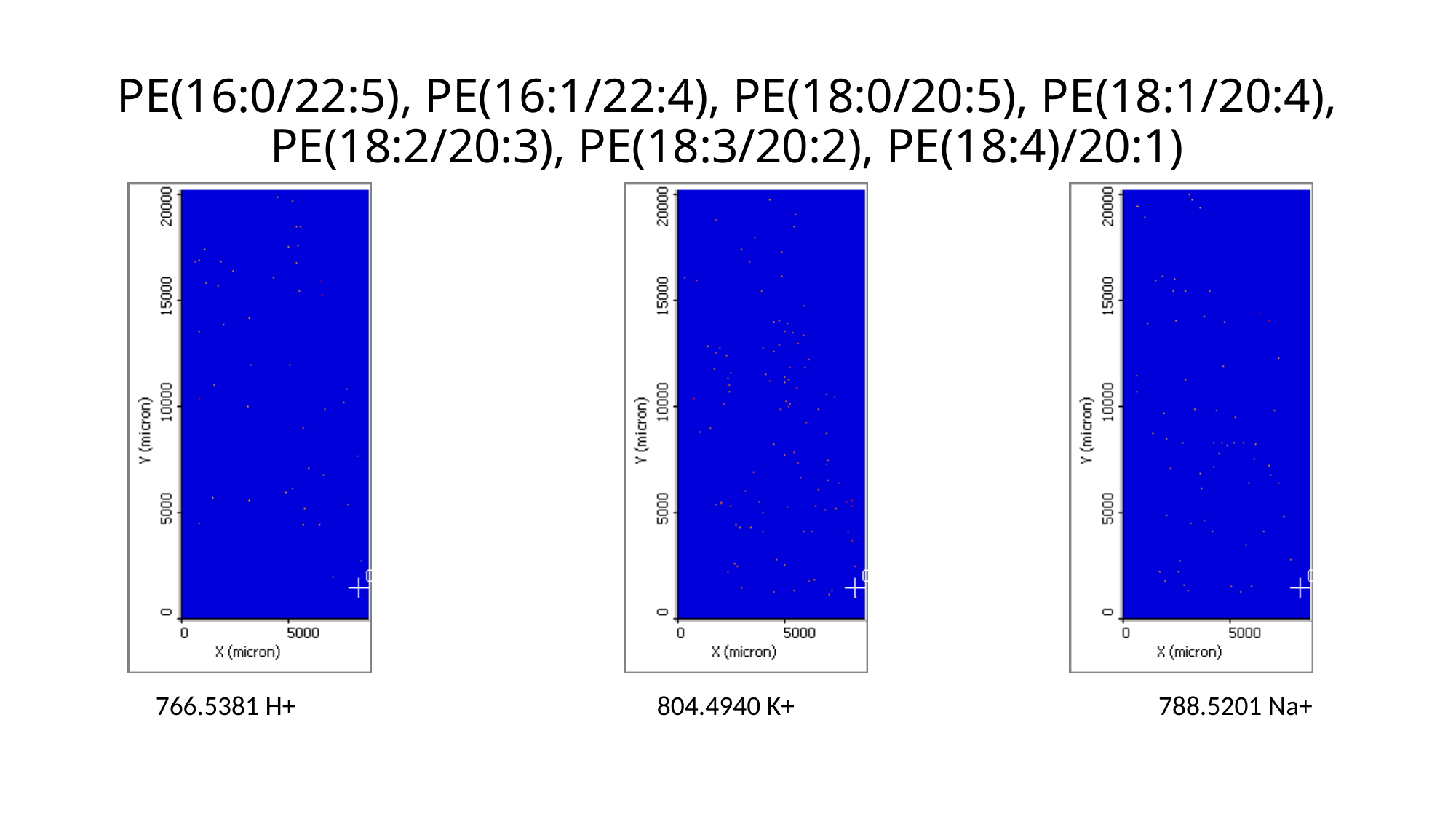

# PE(16:0/22:5), PE(16:1/22:4), PE(18:0/20:5), PE(18:1/20:4), PE(18:2/20:3), PE(18:3/20:2), PE(18:4)/20:1)
766.5381 H+
804.4940 K+
788.5201 Na+

### Slide 35
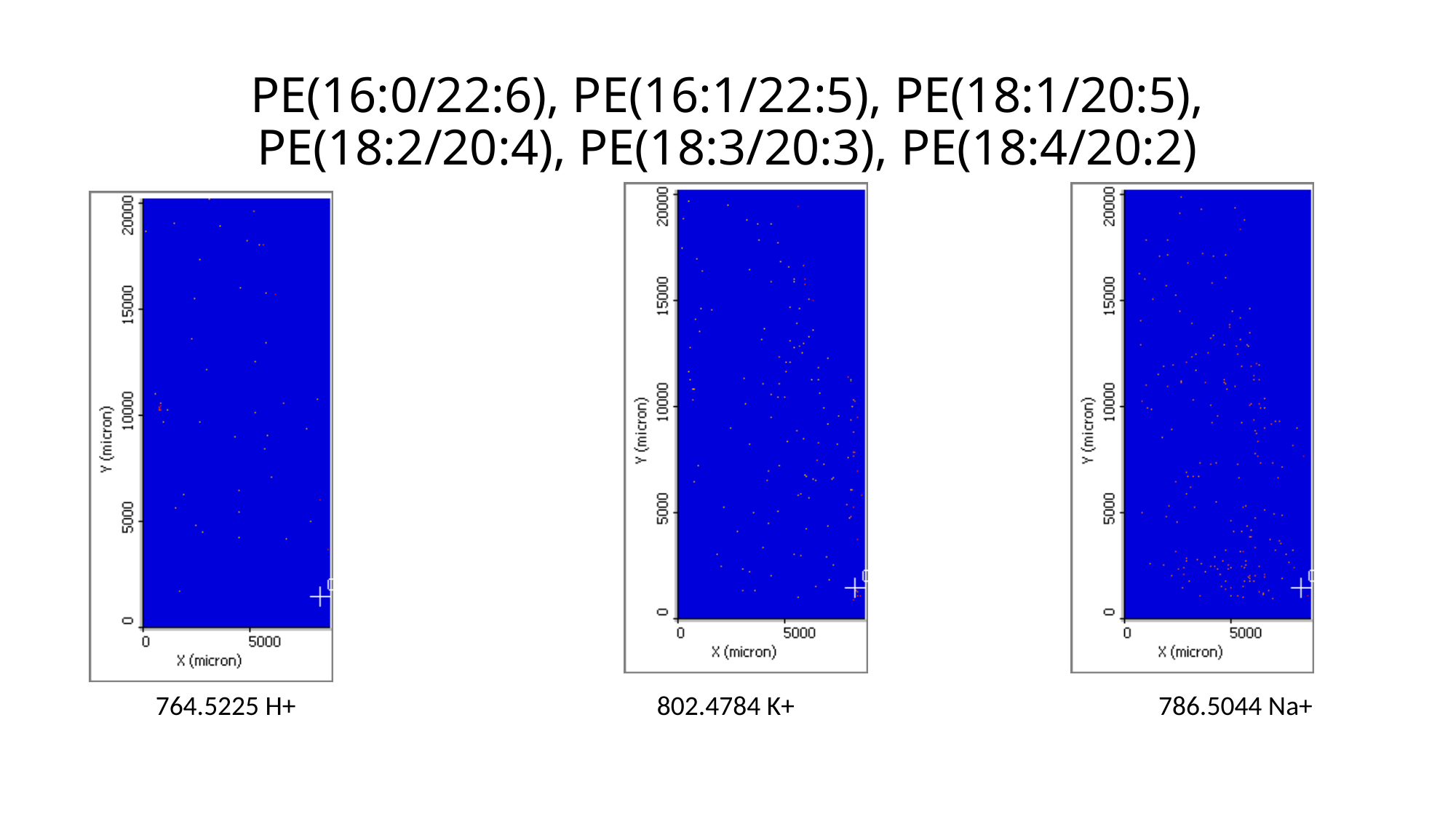

# PE(16:0/22:6), PE(16:1/22:5), PE(18:1/20:5), PE(18:2/20:4), PE(18:3/20:3), PE(18:4/20:2)
764.5225 H+
802.4784 K+
786.5044 Na+

### Slide 36
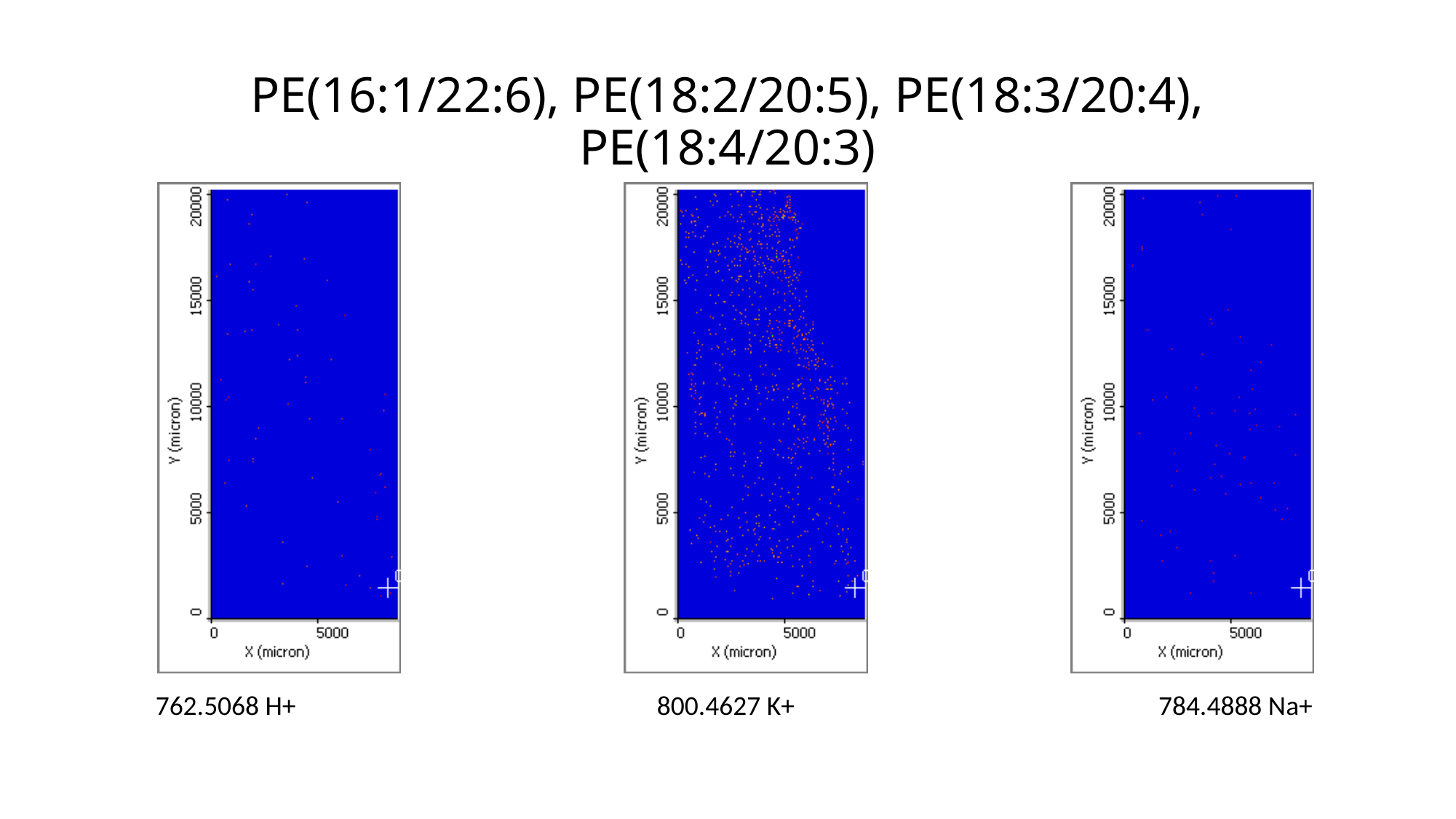

# PE(16:1/22:6), PE(18:2/20:5), PE(18:3/20:4), PE(18:4/20:3)
762.5068 H+
800.4627 K+
784.4888 Na+

### Slide 37
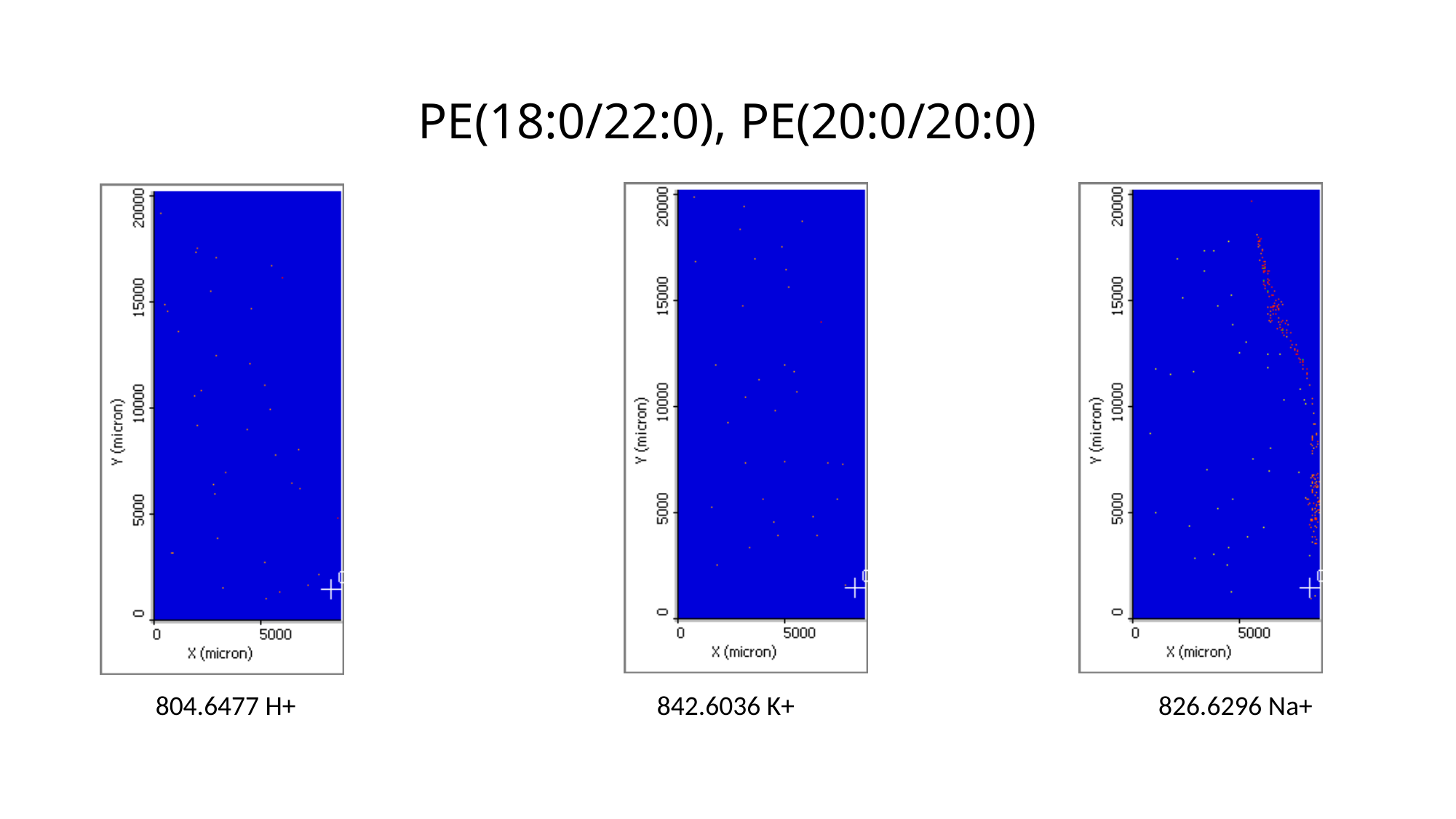

# PE(18:0/22:0), PE(20:0/20:0)
804.6477 H+
842.6036 K+
826.6296 Na+

### Slide 38
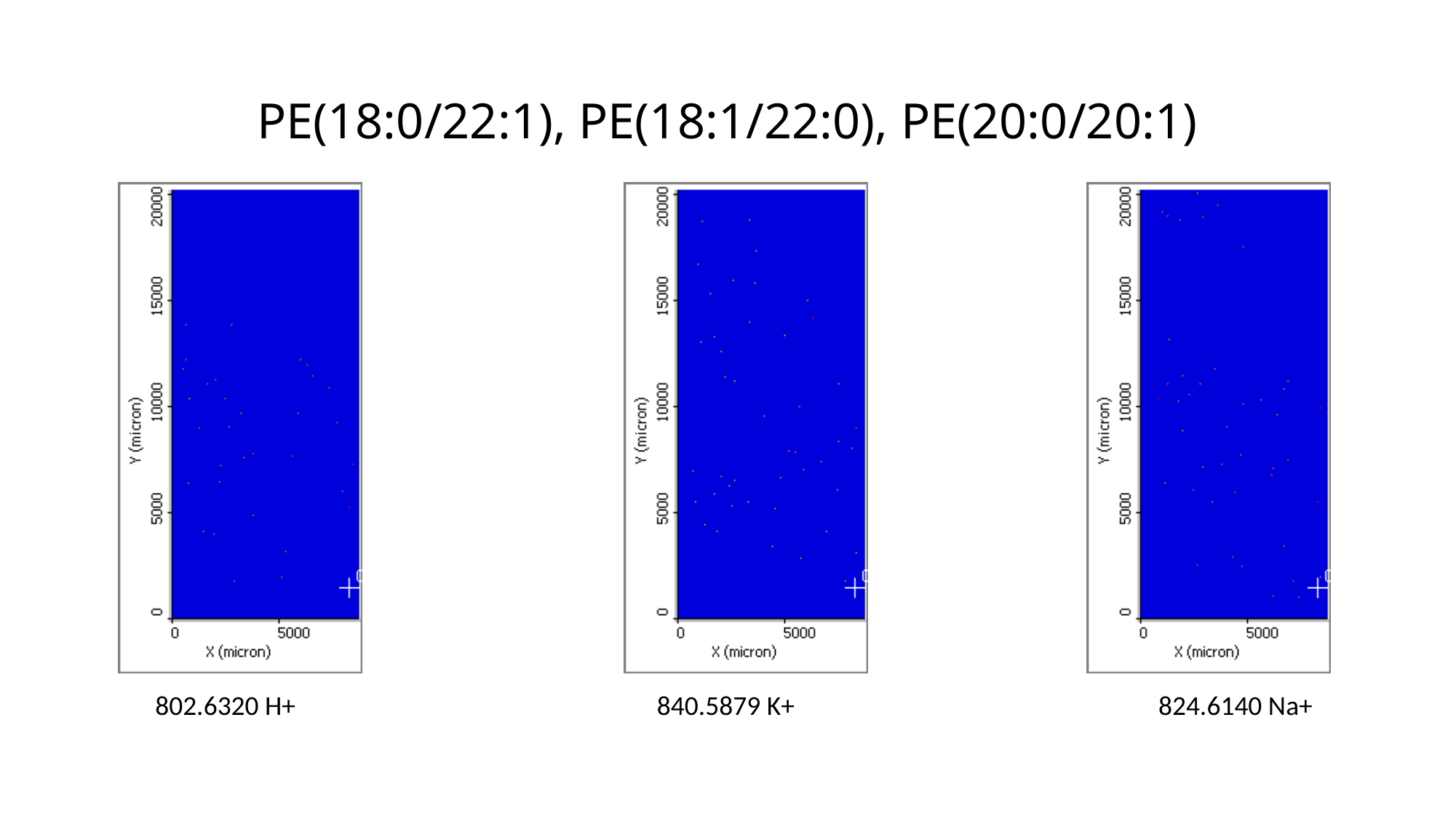

# PE(18:0/22:1), PE(18:1/22:0), PE(20:0/20:1)
802.6320 H+
840.5879 K+
824.6140 Na+

### Slide 39
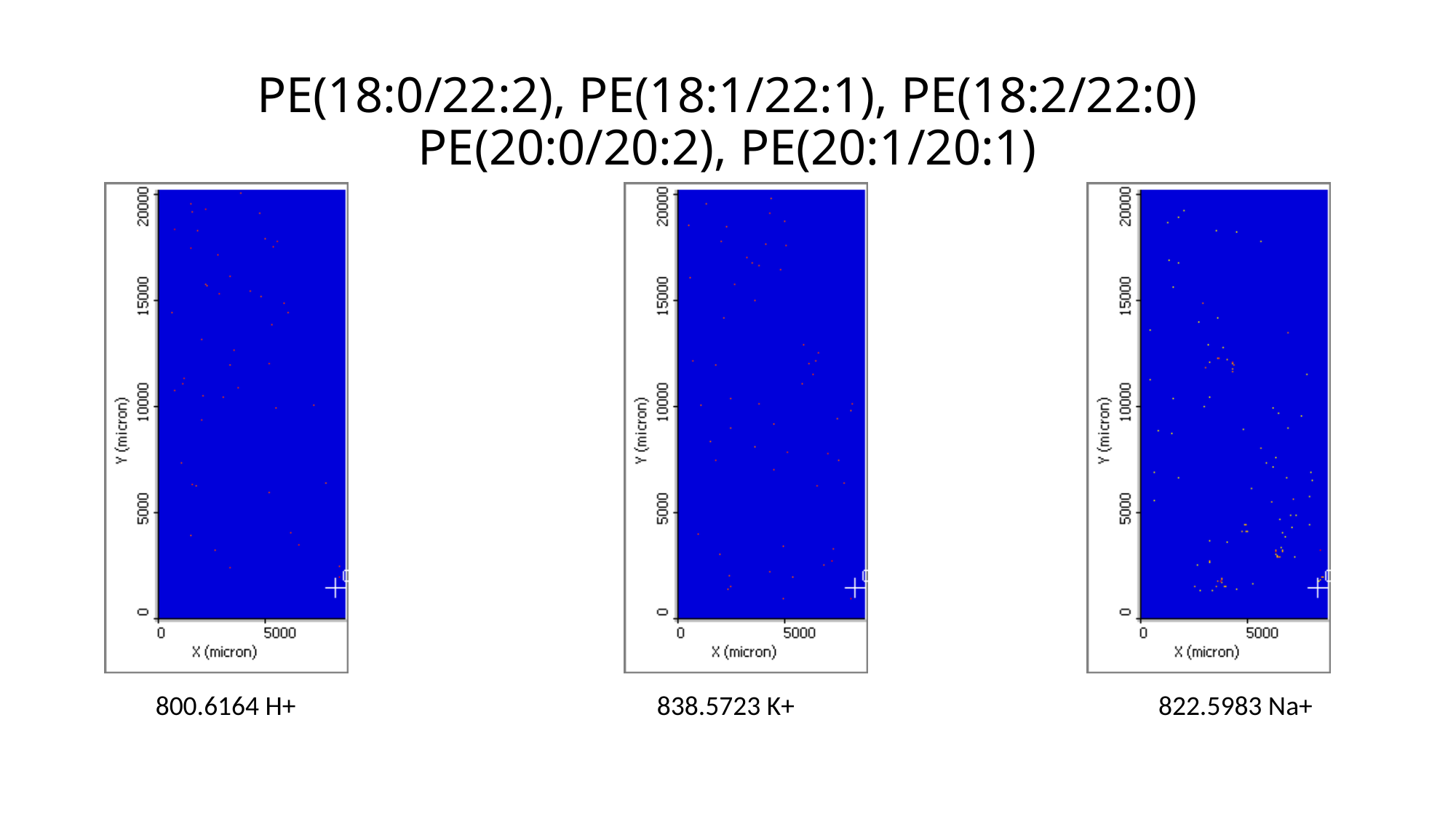

# PE(18:0/22:2), PE(18:1/22:1), PE(18:2/22:0) PE(20:0/20:2), PE(20:1/20:1)
800.6164 H+
838.5723 K+
822.5983 Na+

### Slide 40
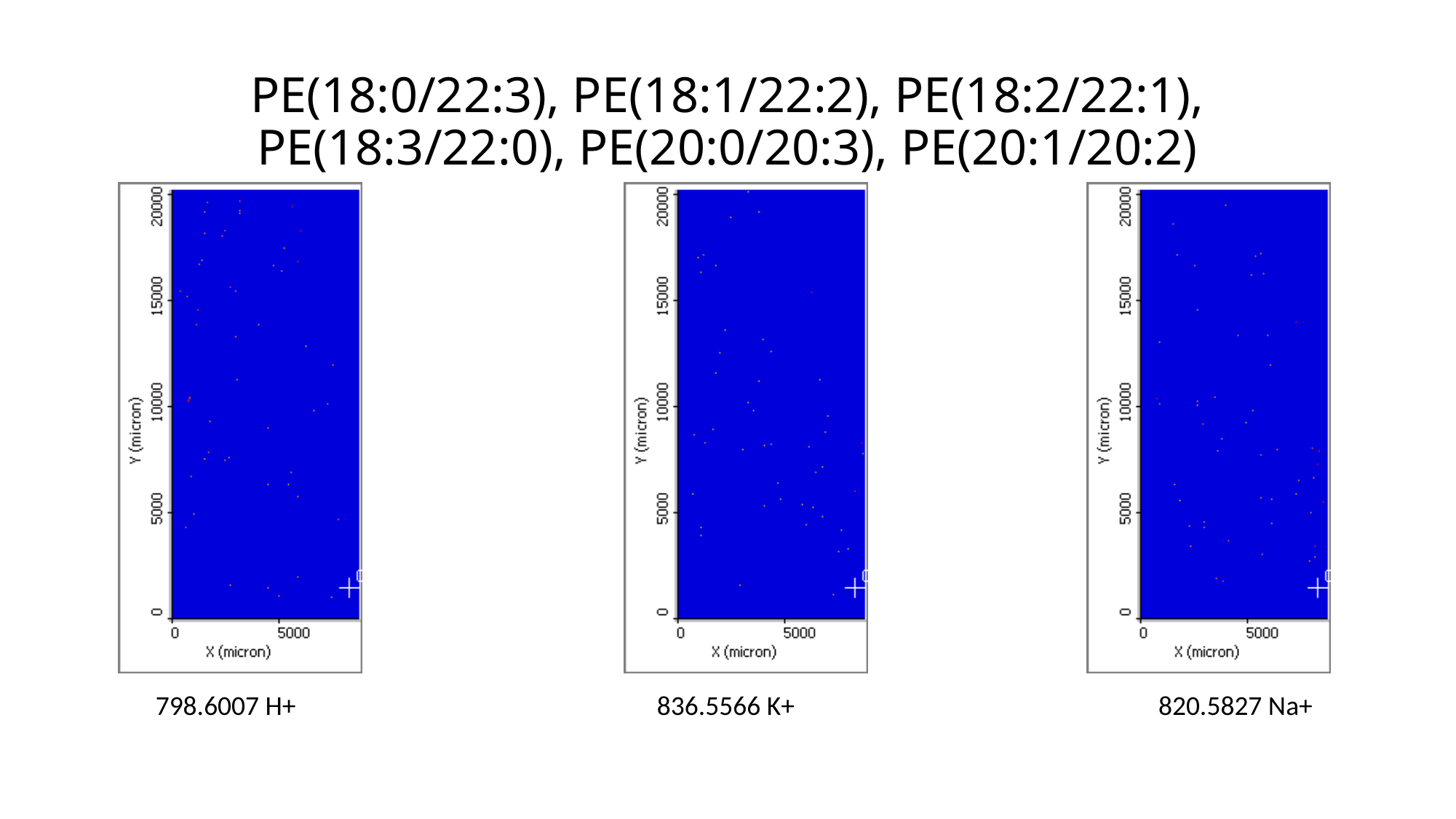

# PE(18:0/22:3), PE(18:1/22:2), PE(18:2/22:1), PE(18:3/22:0), PE(20:0/20:3), PE(20:1/20:2)
798.6007 H+
836.5566 K+
820.5827 Na+

### Slide 41
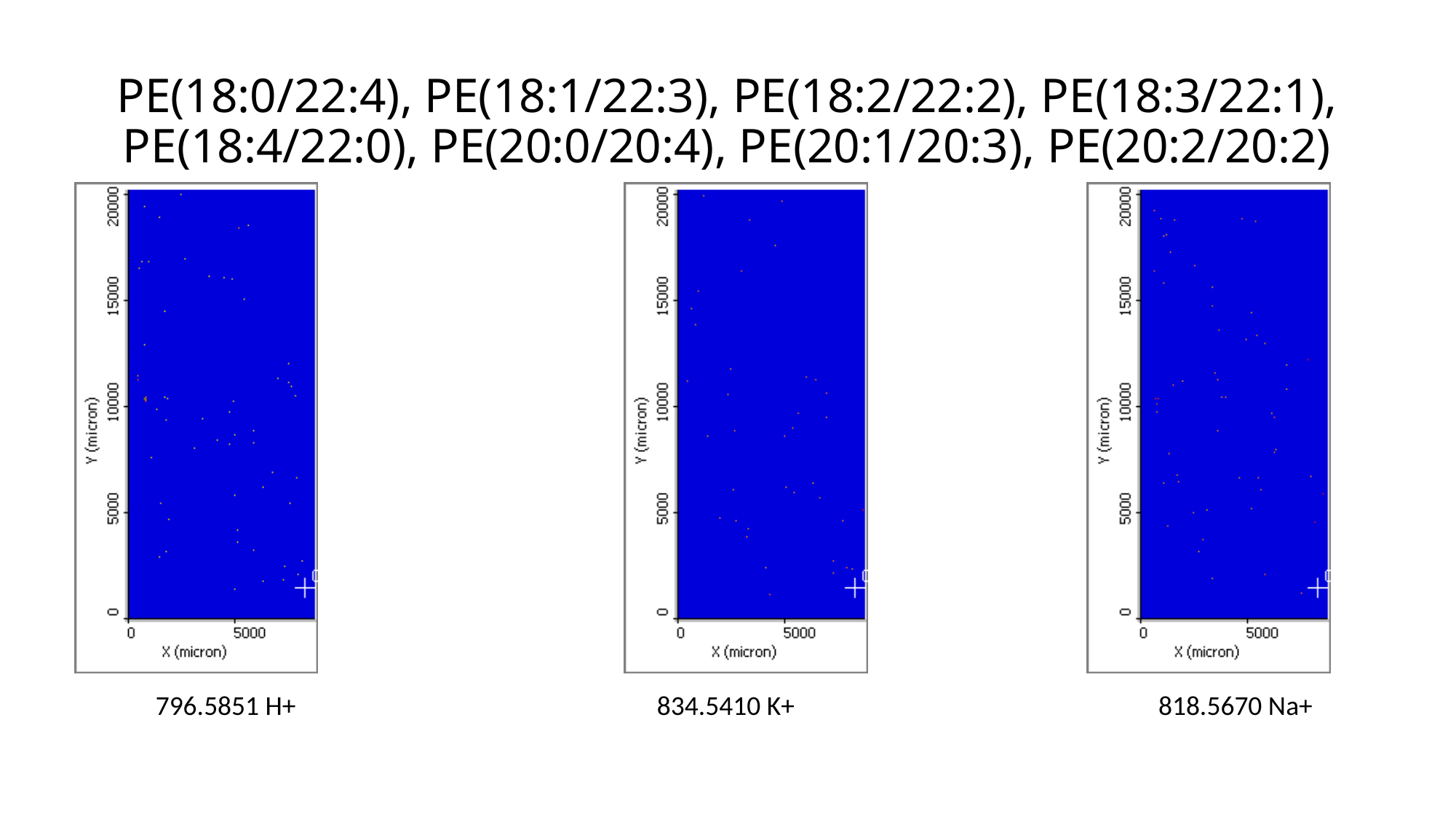

# PE(18:0/22:4), PE(18:1/22:3), PE(18:2/22:2), PE(18:3/22:1), PE(18:4/22:0), PE(20:0/20:4), PE(20:1/20:3), PE(20:2/20:2)
796.5851 H+
834.5410 K+
818.5670 Na+

### Slide 42
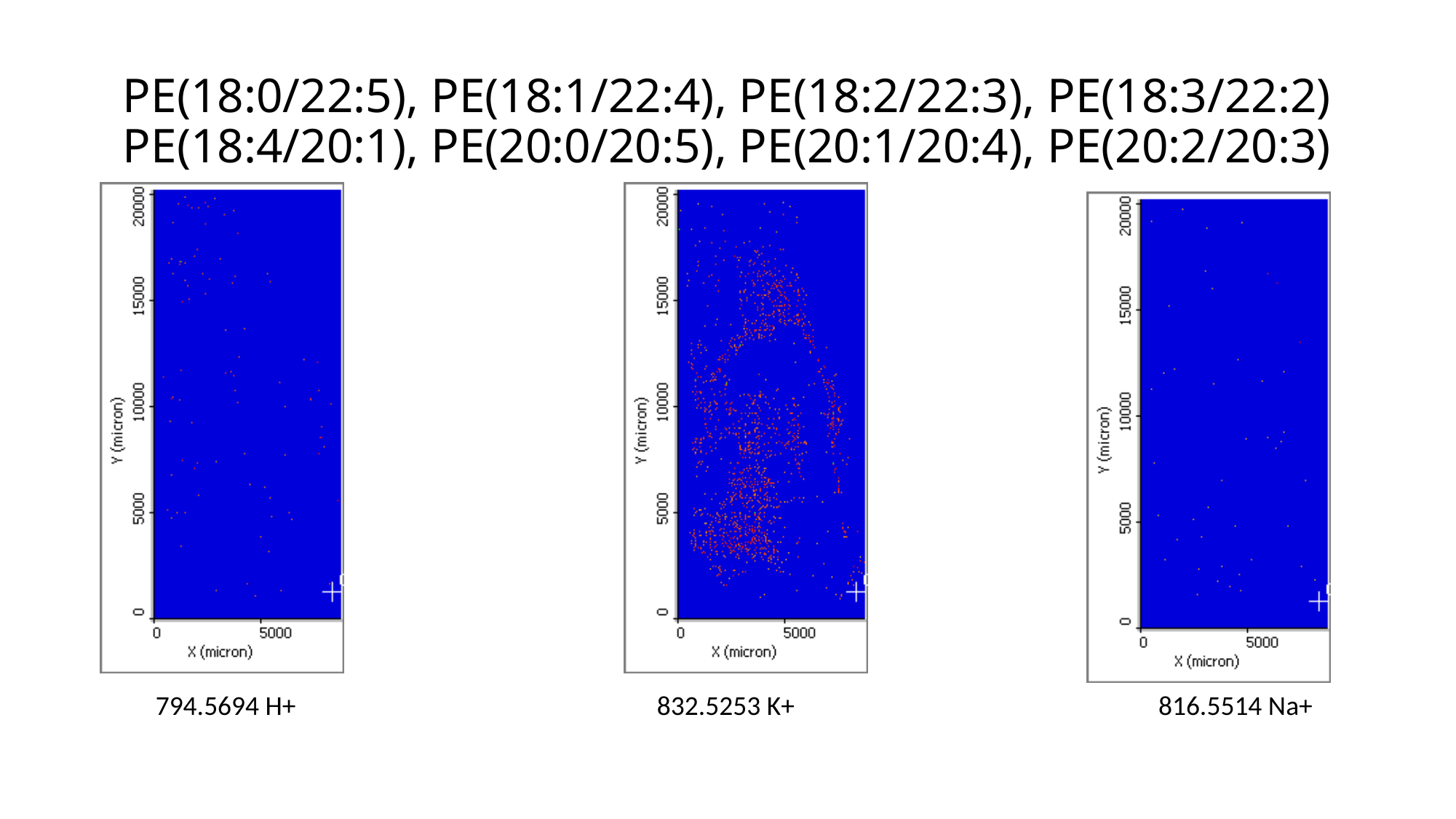

# PE(18:0/22:5), PE(18:1/22:4), PE(18:2/22:3), PE(18:3/22:2) PE(18:4/20:1), PE(20:0/20:5), PE(20:1/20:4), PE(20:2/20:3)
794.5694 H+
832.5253 K+
816.5514 Na+

### Slide 43
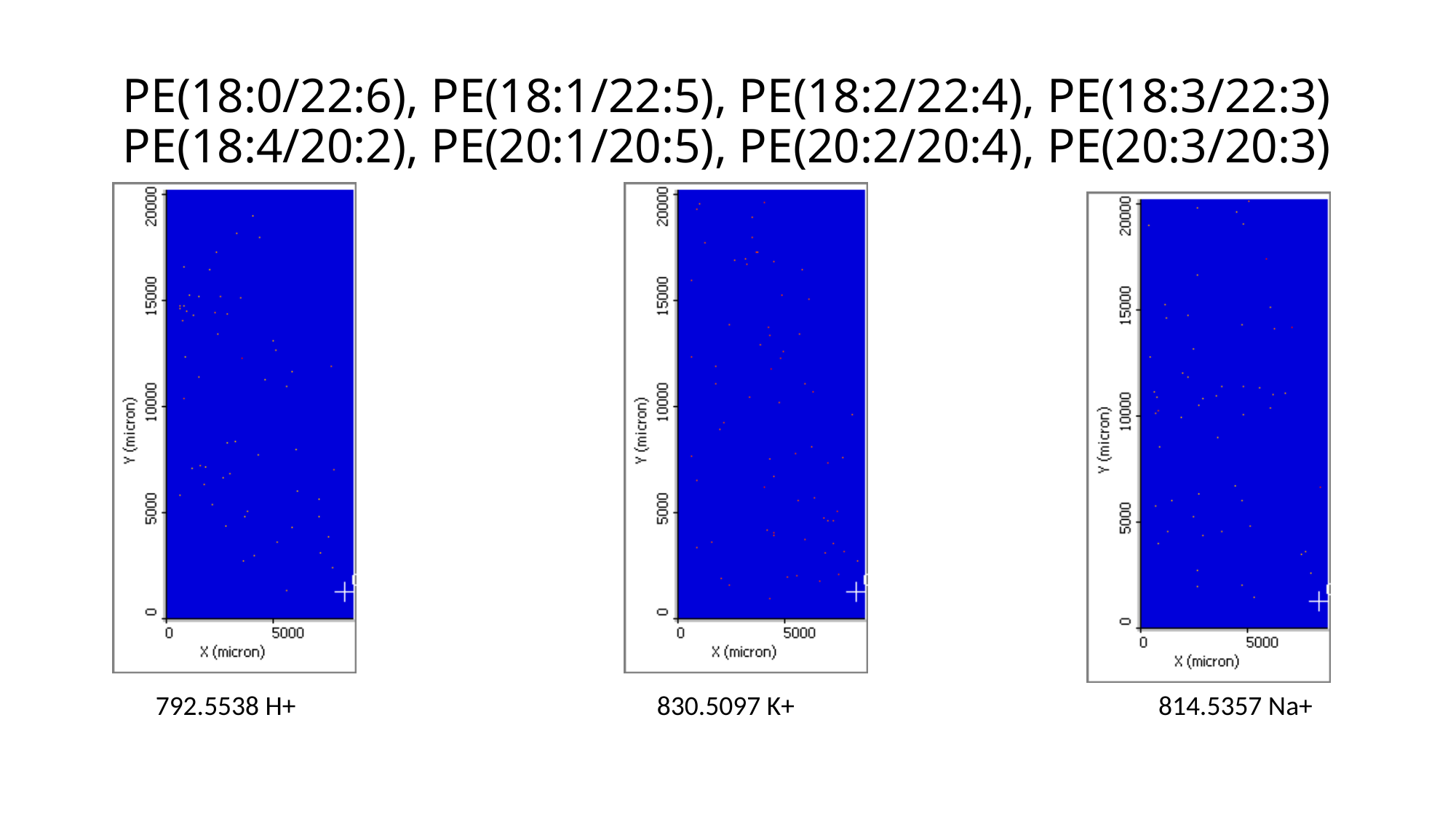

# PE(18:0/22:6), PE(18:1/22:5), PE(18:2/22:4), PE(18:3/22:3) PE(18:4/20:2), PE(20:1/20:5), PE(20:2/20:4), PE(20:3/20:3)
792.5538 H+
830.5097 K+
814.5357 Na+

### Slide 44
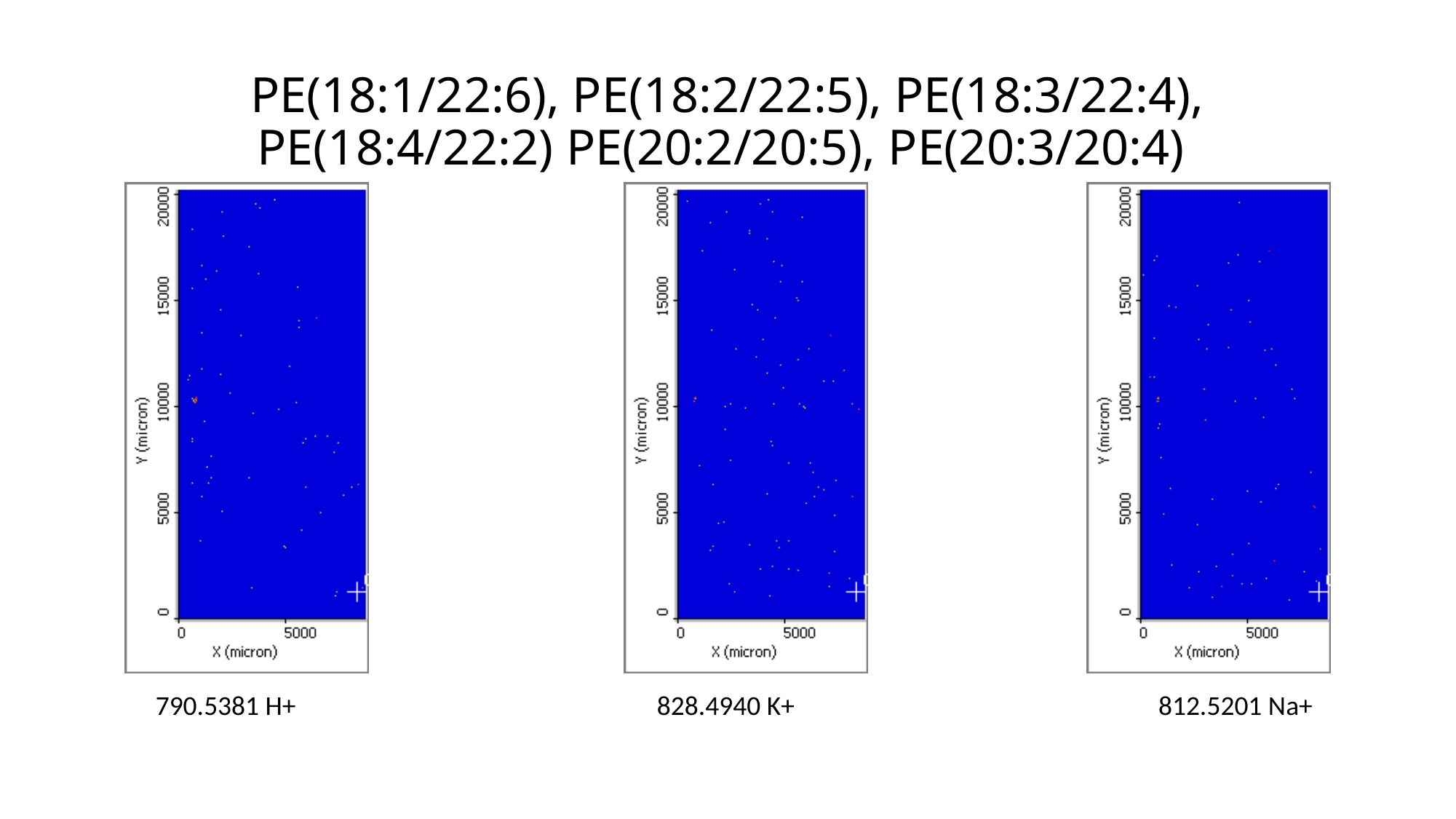

# PE(18:1/22:6), PE(18:2/22:5), PE(18:3/22:4), PE(18:4/22:2) PE(20:2/20:5), PE(20:3/20:4)
790.5381 H+
828.4940 K+
812.5201 Na+

### Slide 45
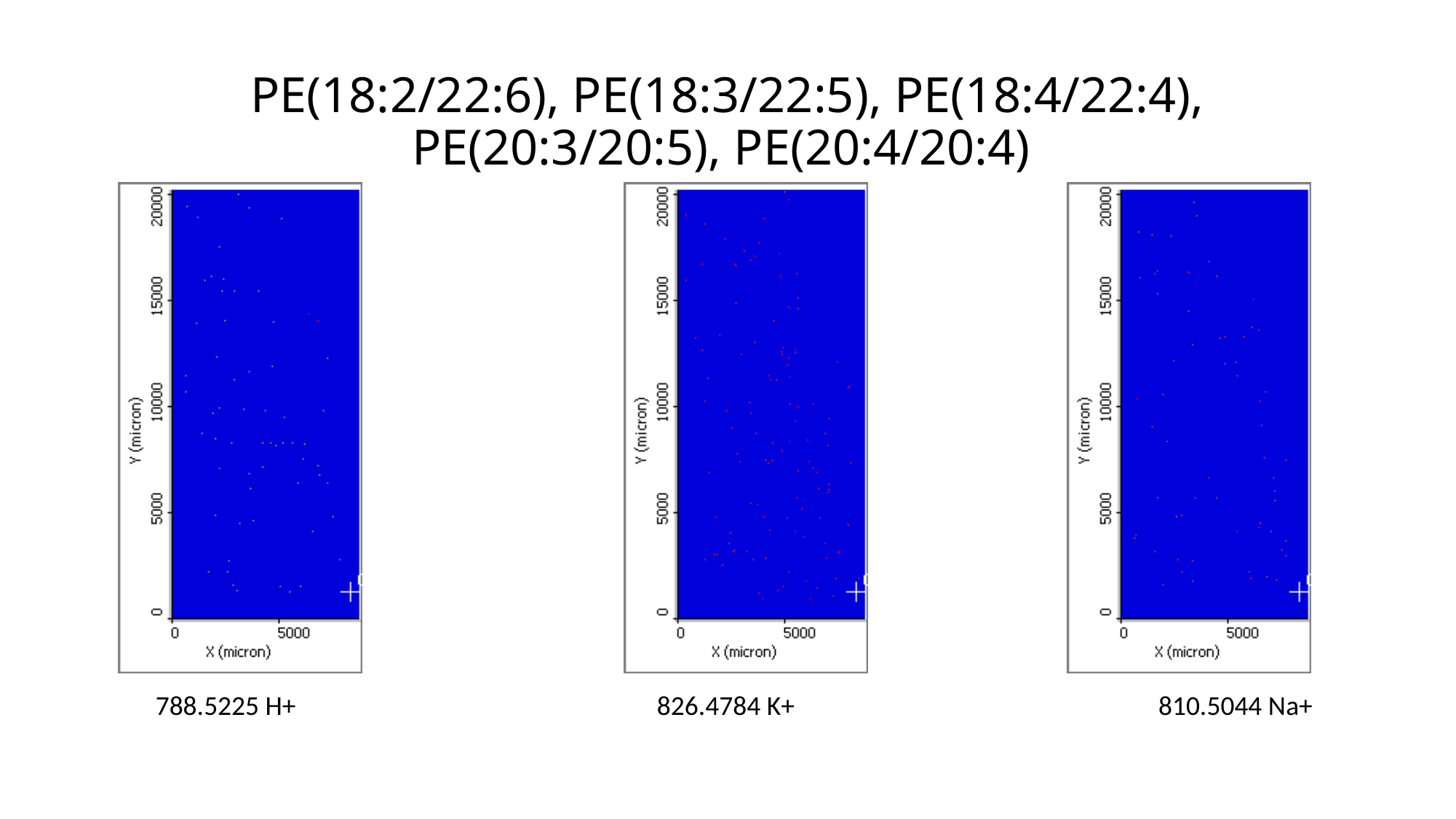

# PE(18:2/22:6), PE(18:3/22:5), PE(18:4/22:4), PE(20:3/20:5), PE(20:4/20:4)
788.5225 H+
826.4784 K+
810.5044 Na+

### Slide 46
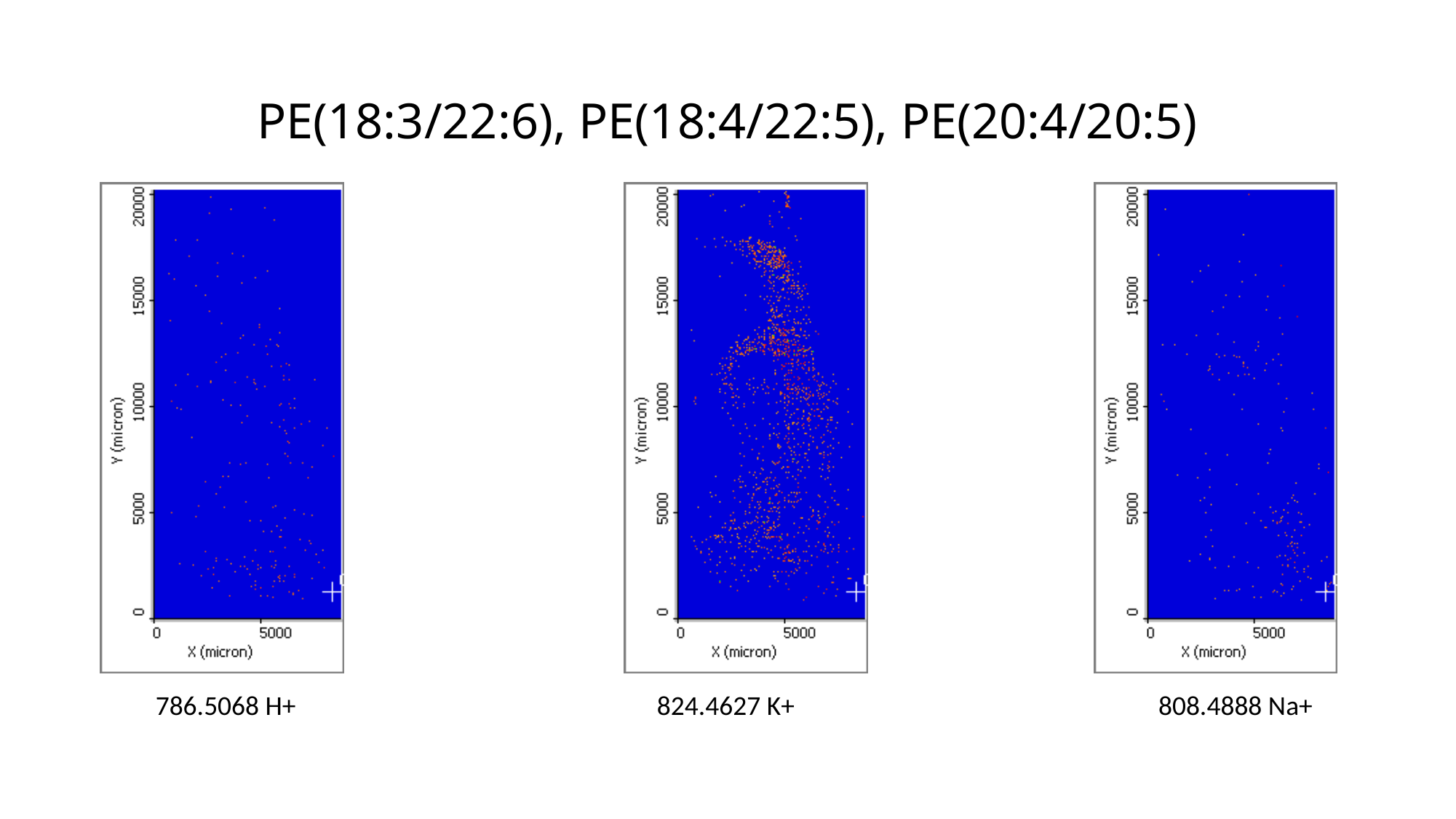

# PE(18:3/22:6), PE(18:4/22:5), PE(20:4/20:5)
786.5068 H+
824.4627 K+
808.4888 Na+

### Slide 47
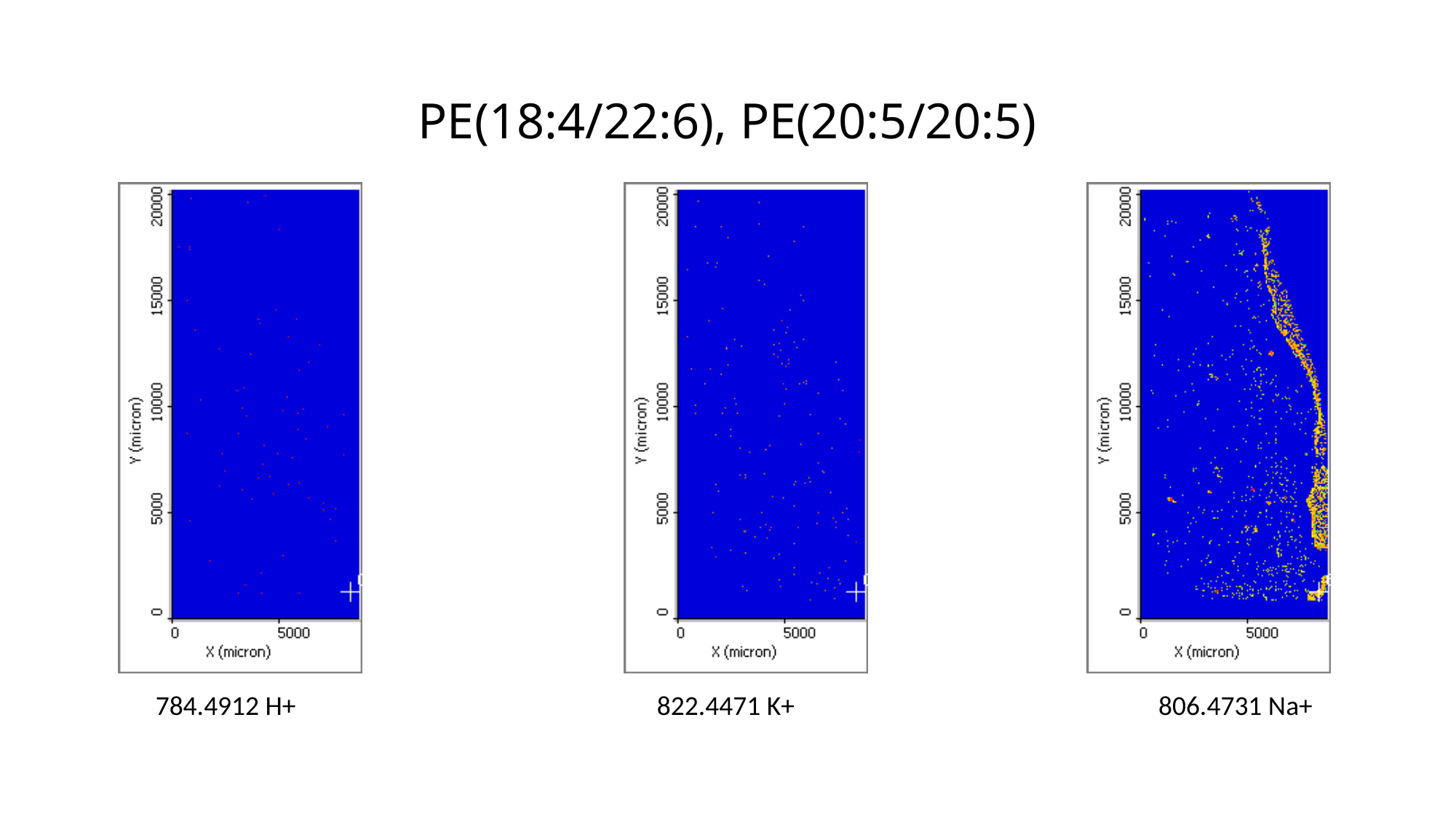

# PE(18:4/22:6), PE(20:5/20:5)
784.4912 H+
822.4471 K+
806.4731 Na+

### Slide 48
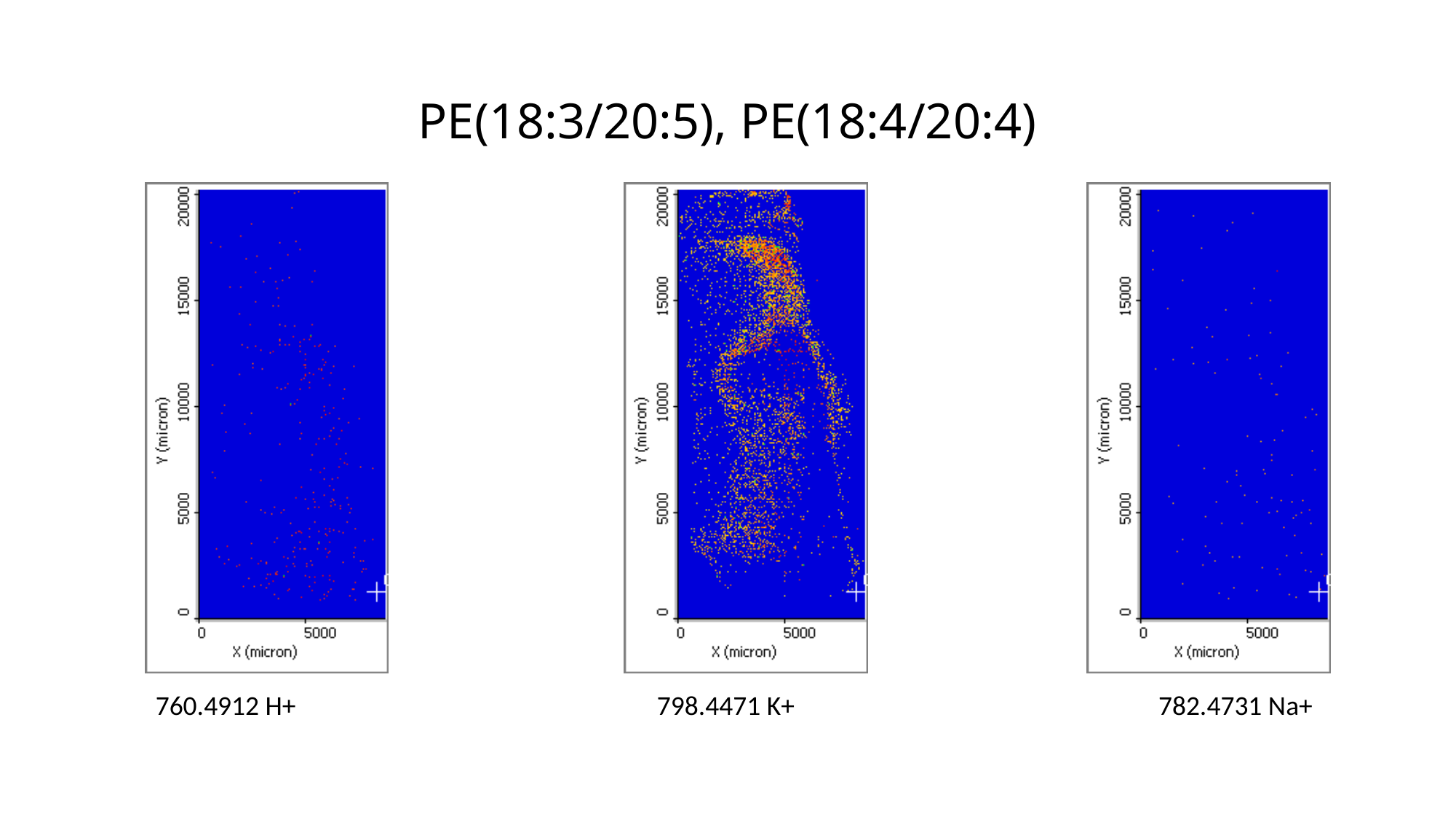

# PE(18:3/20:5), PE(18:4/20:4)
760.4912 H+
798.4471 K+
782.4731 Na+

### Slide 49
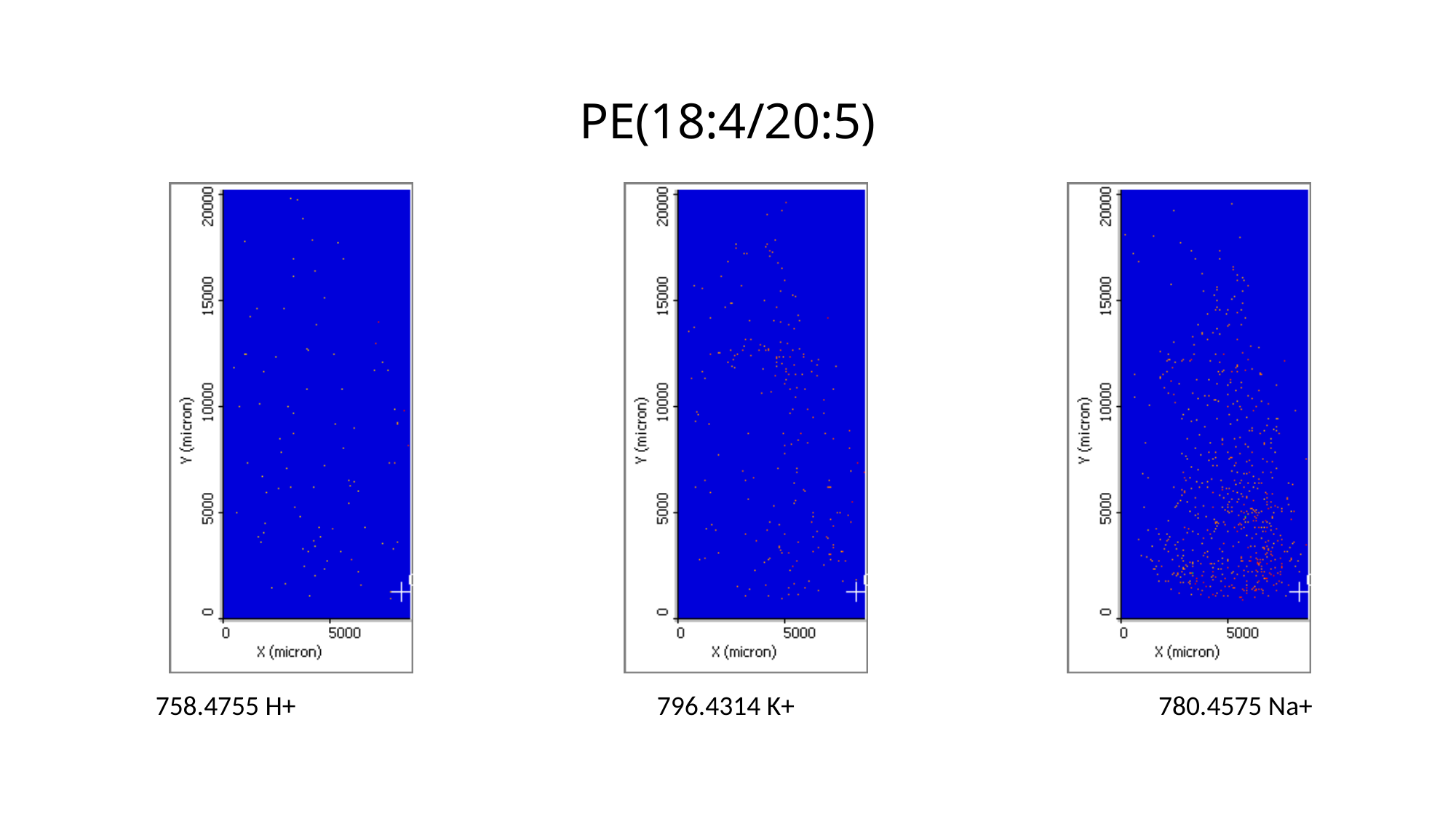

# PE(18:4/20:5)
758.4755 H+
796.4314 K+
780.4575 Na+

### Slide 50
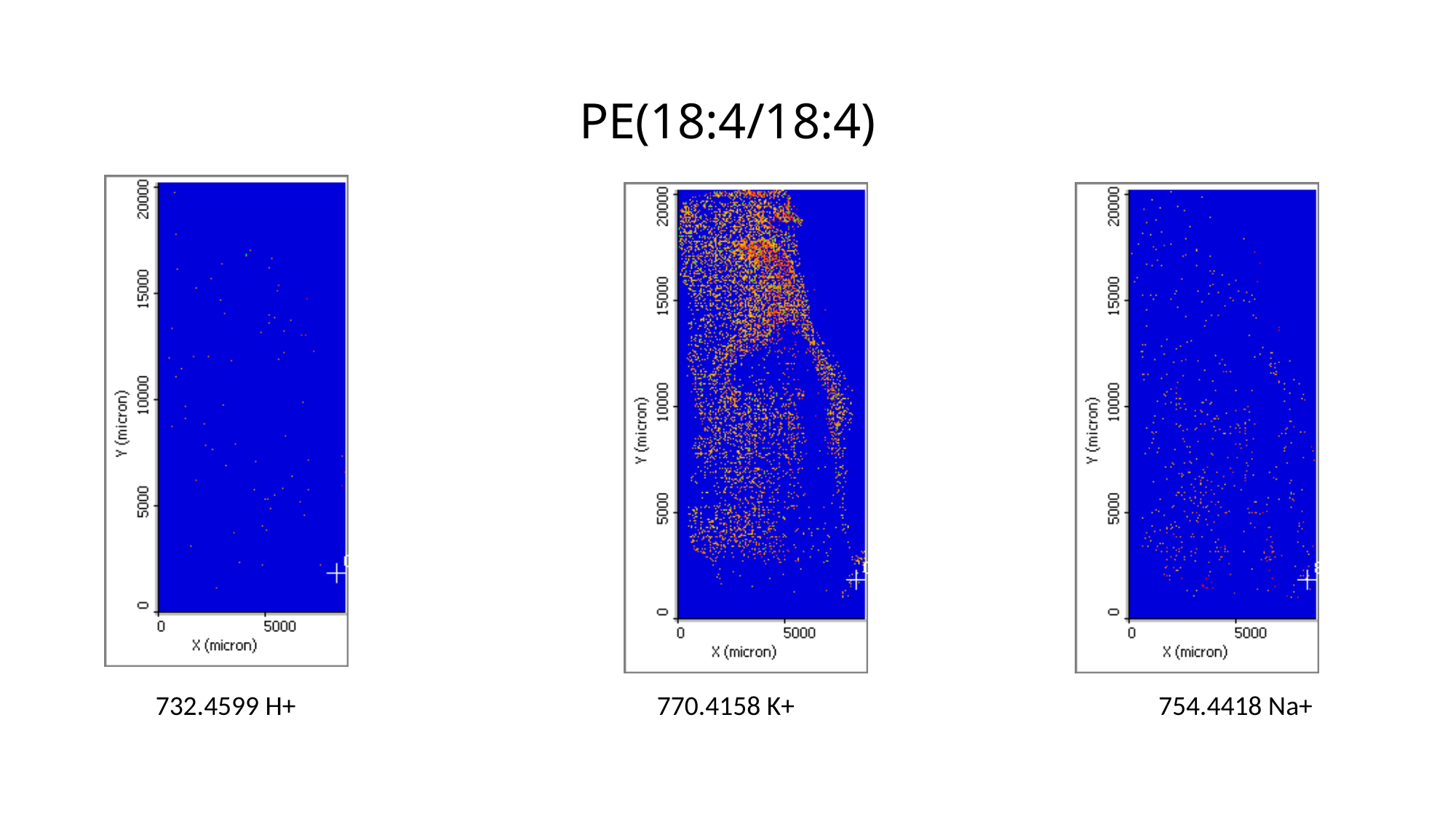

# PE(18:4/18:4)
732.4599 H+
770.4158 K+
754.4418 Na+

### Slide 51
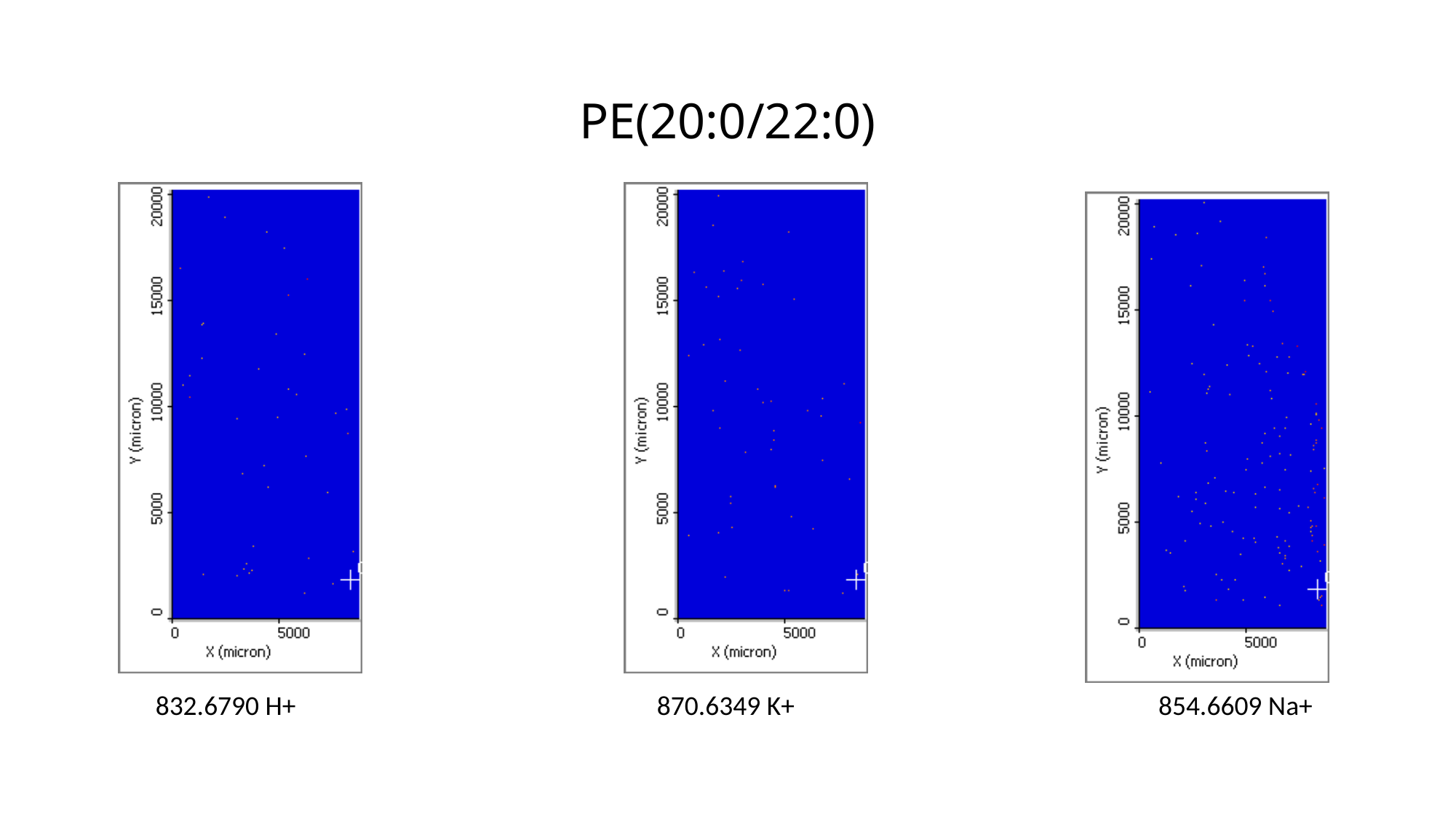

# PE(20:0/22:0)
832.6790 H+
870.6349 K+
854.6609 Na+

### Slide 52
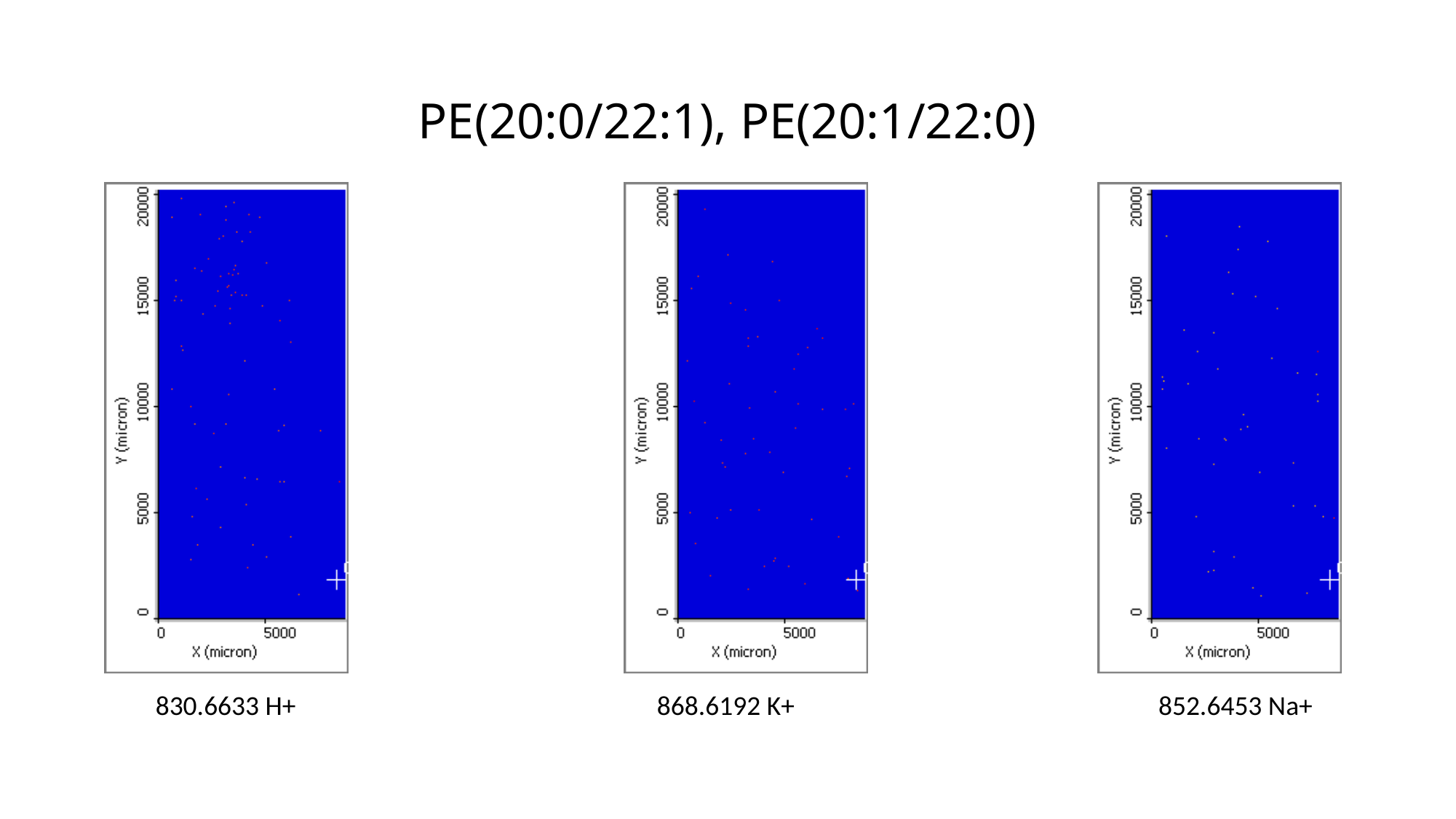

# PE(20:0/22:1), PE(20:1/22:0)
830.6633 H+
868.6192 K+
852.6453 Na+

### Slide 53
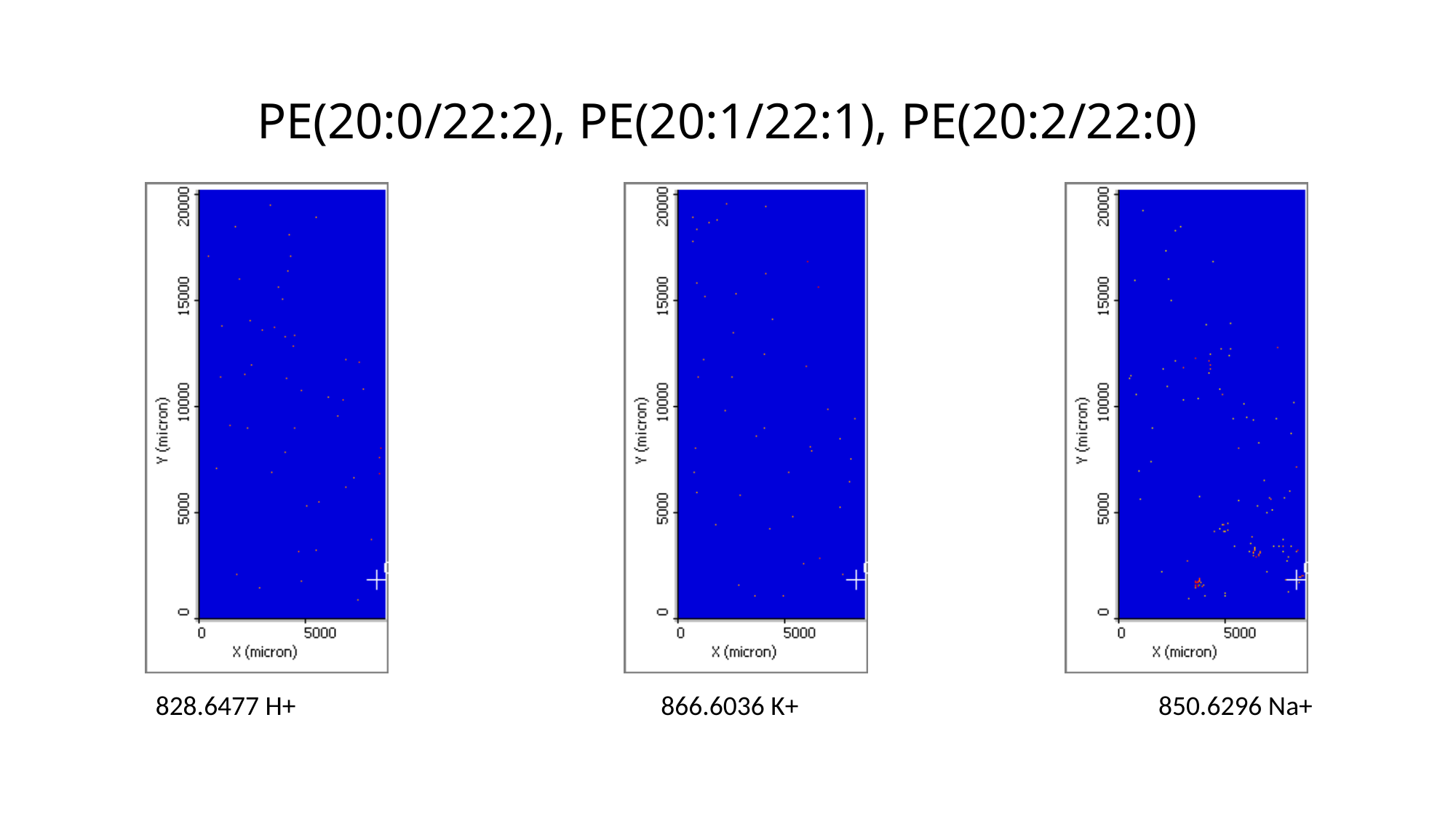

# PE(20:0/22:2), PE(20:1/22:1), PE(20:2/22:0)
828.6477 H+
 866.6036 K+
850.6296 Na+

### Slide 54
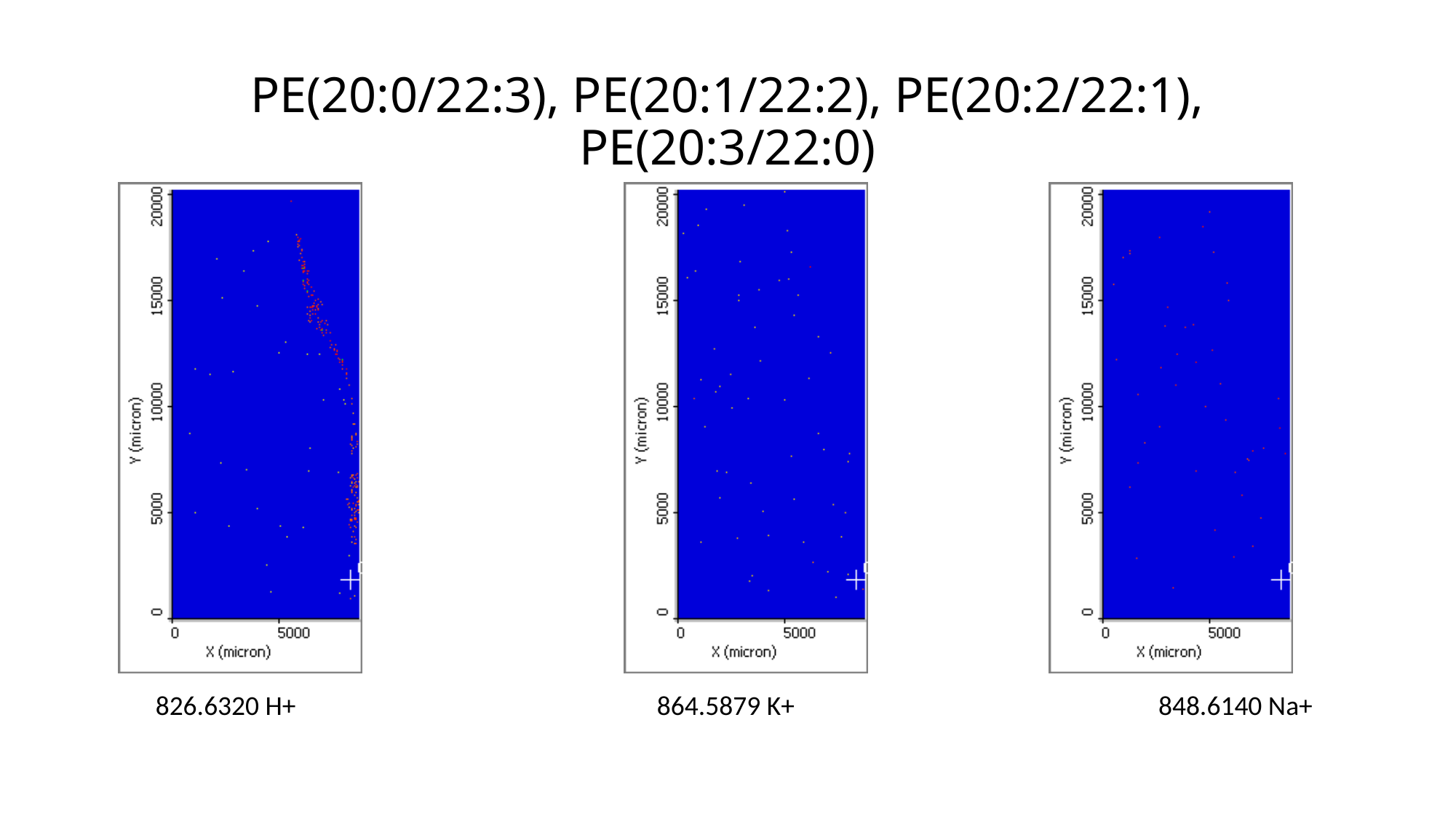

# PE(20:0/22:3), PE(20:1/22:2), PE(20:2/22:1), PE(20:3/22:0)
826.6320 H+
864.5879 K+
848.6140 Na+

### Slide 55
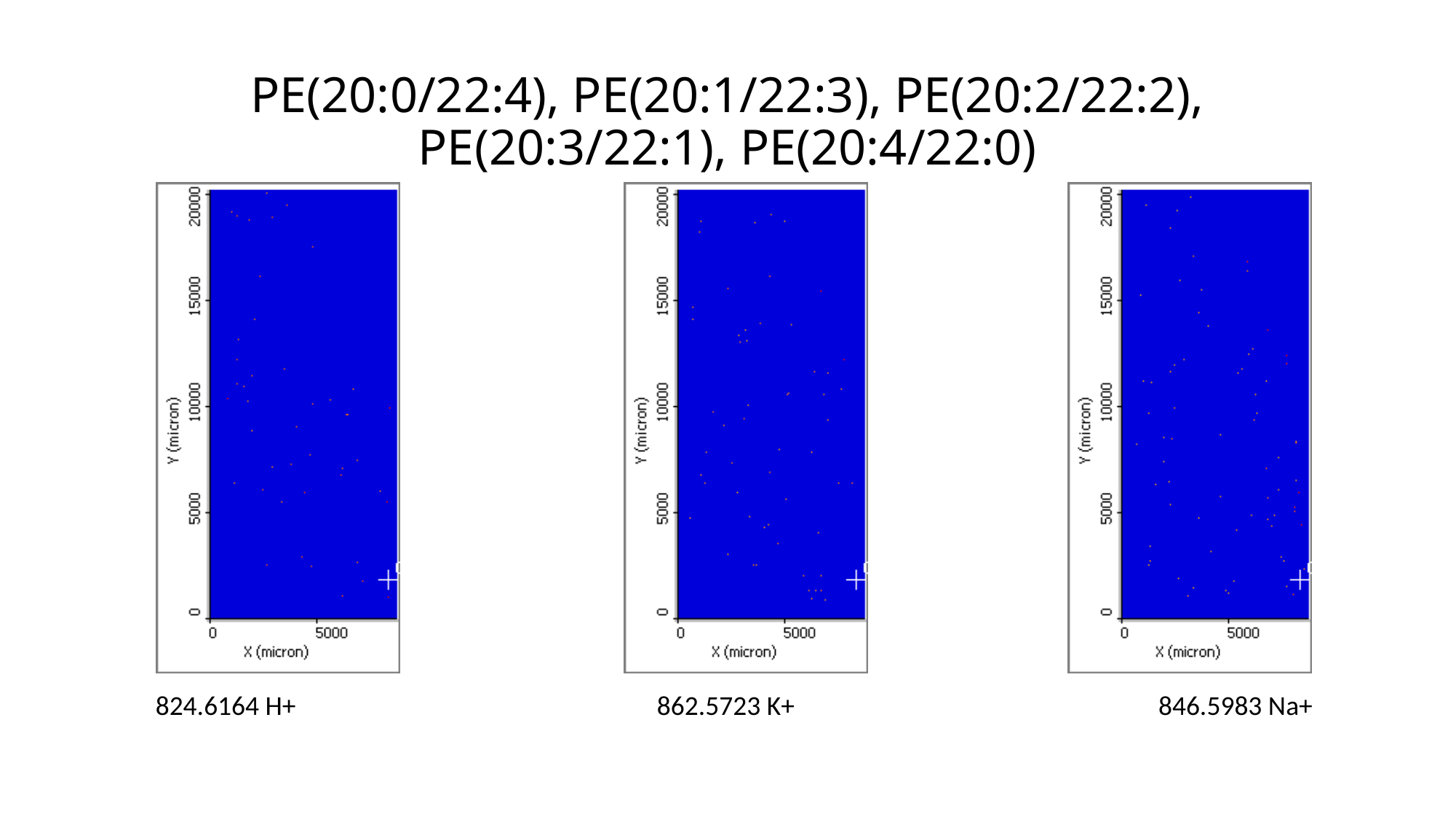

# PE(20:0/22:4), PE(20:1/22:3), PE(20:2/22:2), PE(20:3/22:1), PE(20:4/22:0)
824.6164 H+
862.5723 K+
846.5983 Na+

### Slide 56
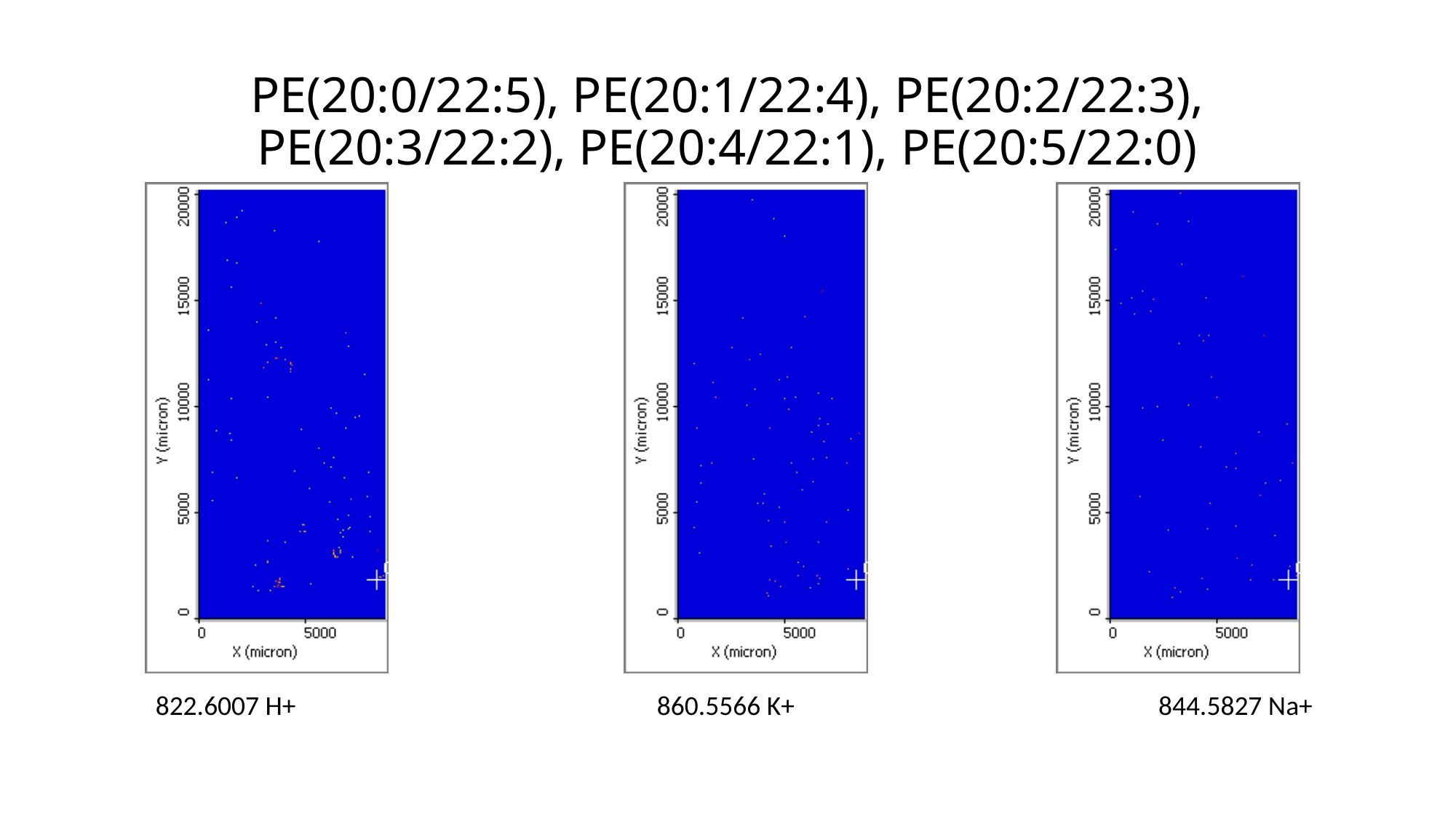

# PE(20:0/22:5), PE(20:1/22:4), PE(20:2/22:3), PE(20:3/22:2), PE(20:4/22:1), PE(20:5/22:0)
822.6007 H+
860.5566 K+
844.5827 Na+

### Slide 57
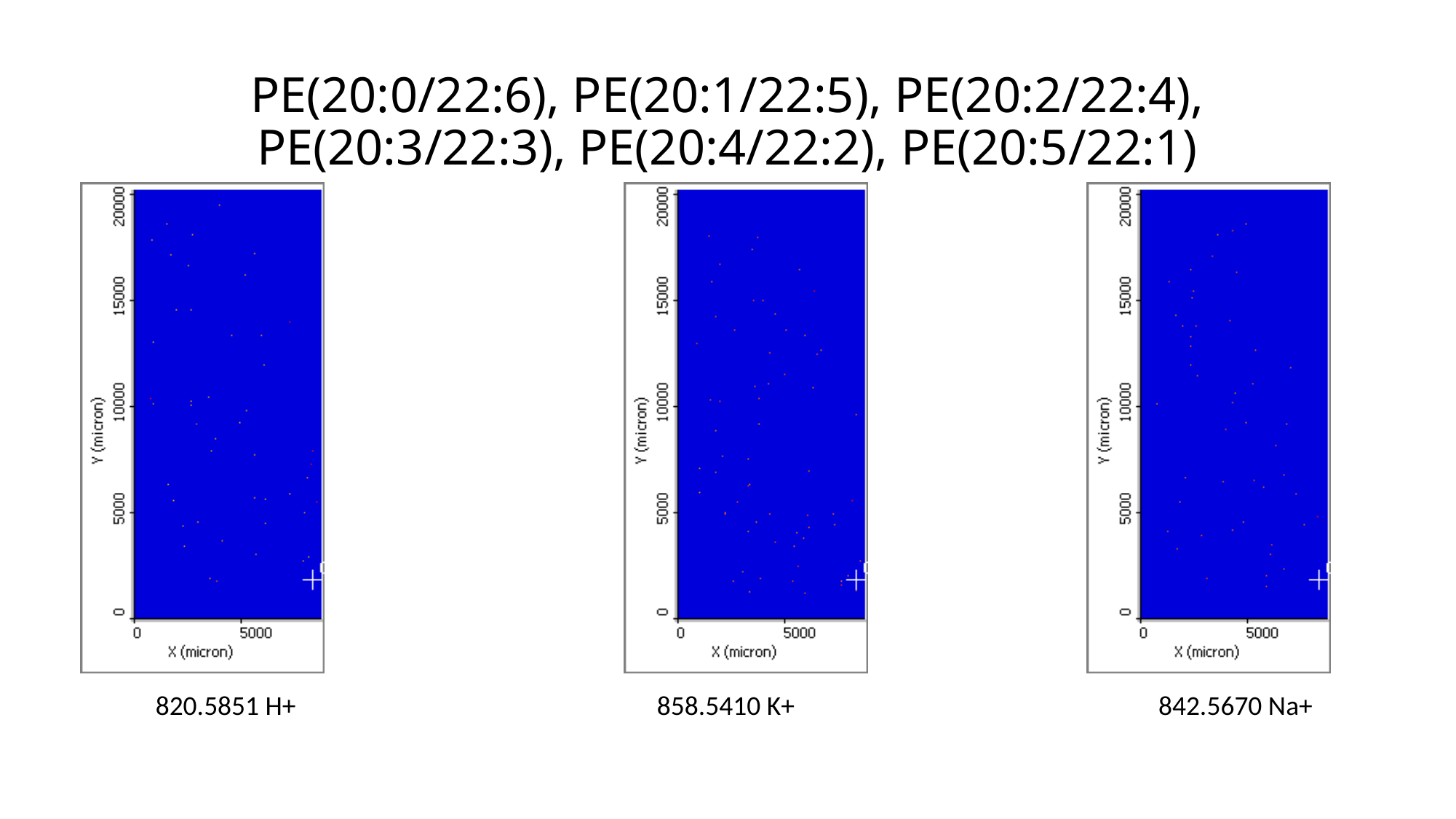

# PE(20:0/22:6), PE(20:1/22:5), PE(20:2/22:4), PE(20:3/22:3), PE(20:4/22:2), PE(20:5/22:1)
820.5851 H+
858.5410 K+
842.5670 Na+

### Slide 58
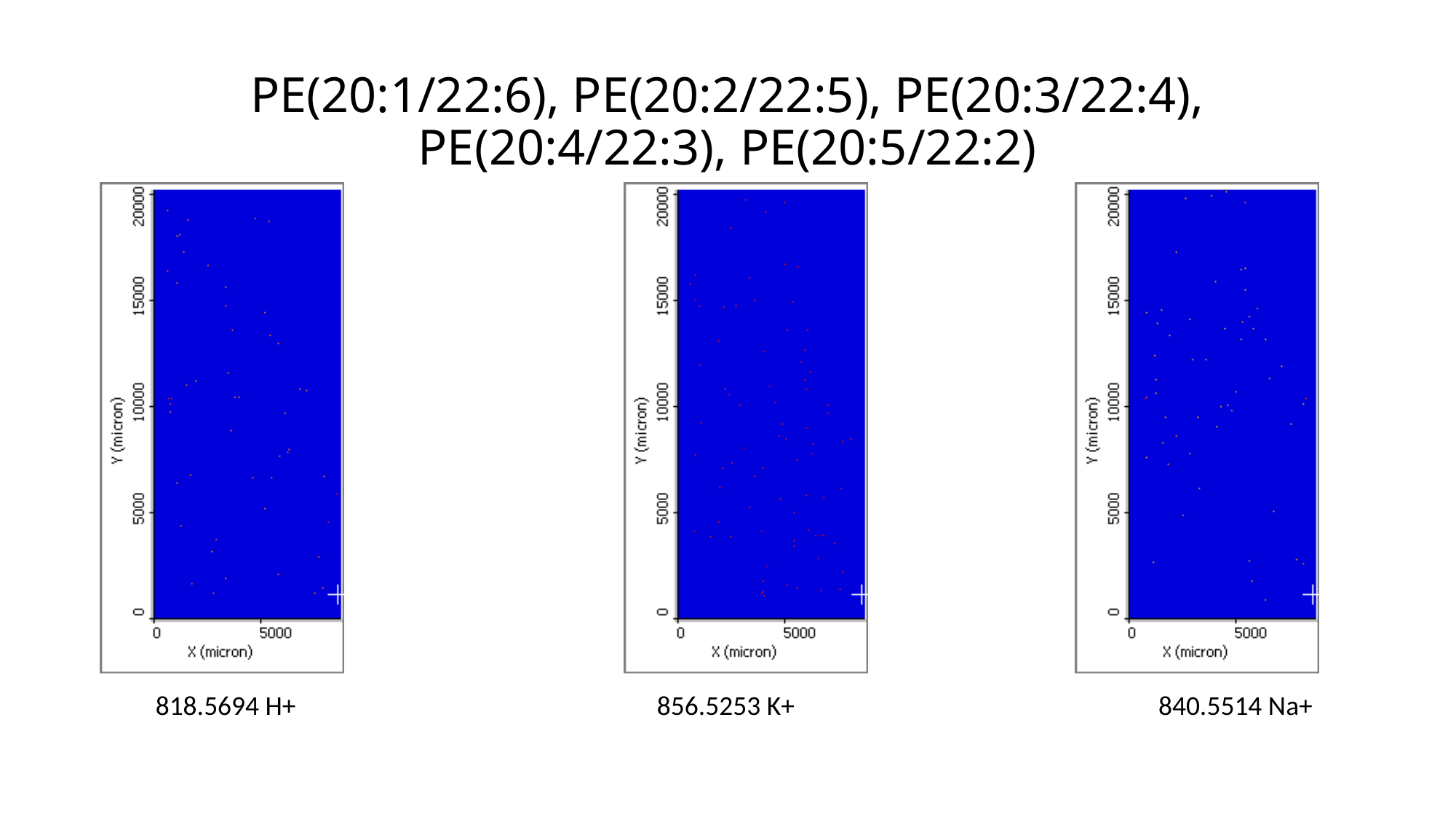

# PE(20:1/22:6), PE(20:2/22:5), PE(20:3/22:4), PE(20:4/22:3), PE(20:5/22:2)
818.5694 H+
856.5253 K+
840.5514 Na+

### Slide 59
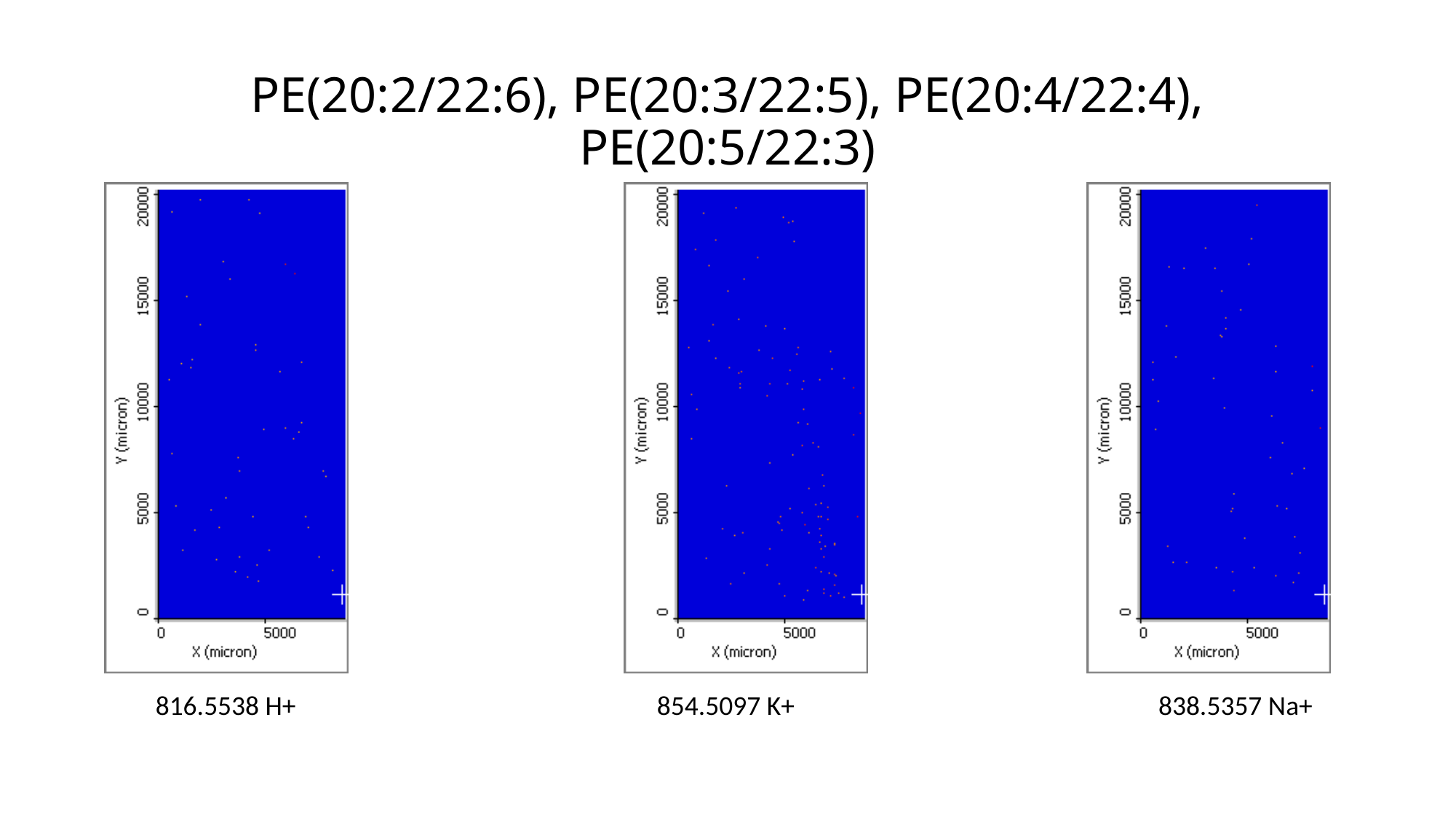

# PE(20:2/22:6), PE(20:3/22:5), PE(20:4/22:4), PE(20:5/22:3)
816.5538 H+
854.5097 K+
838.5357 Na+

### Slide 60
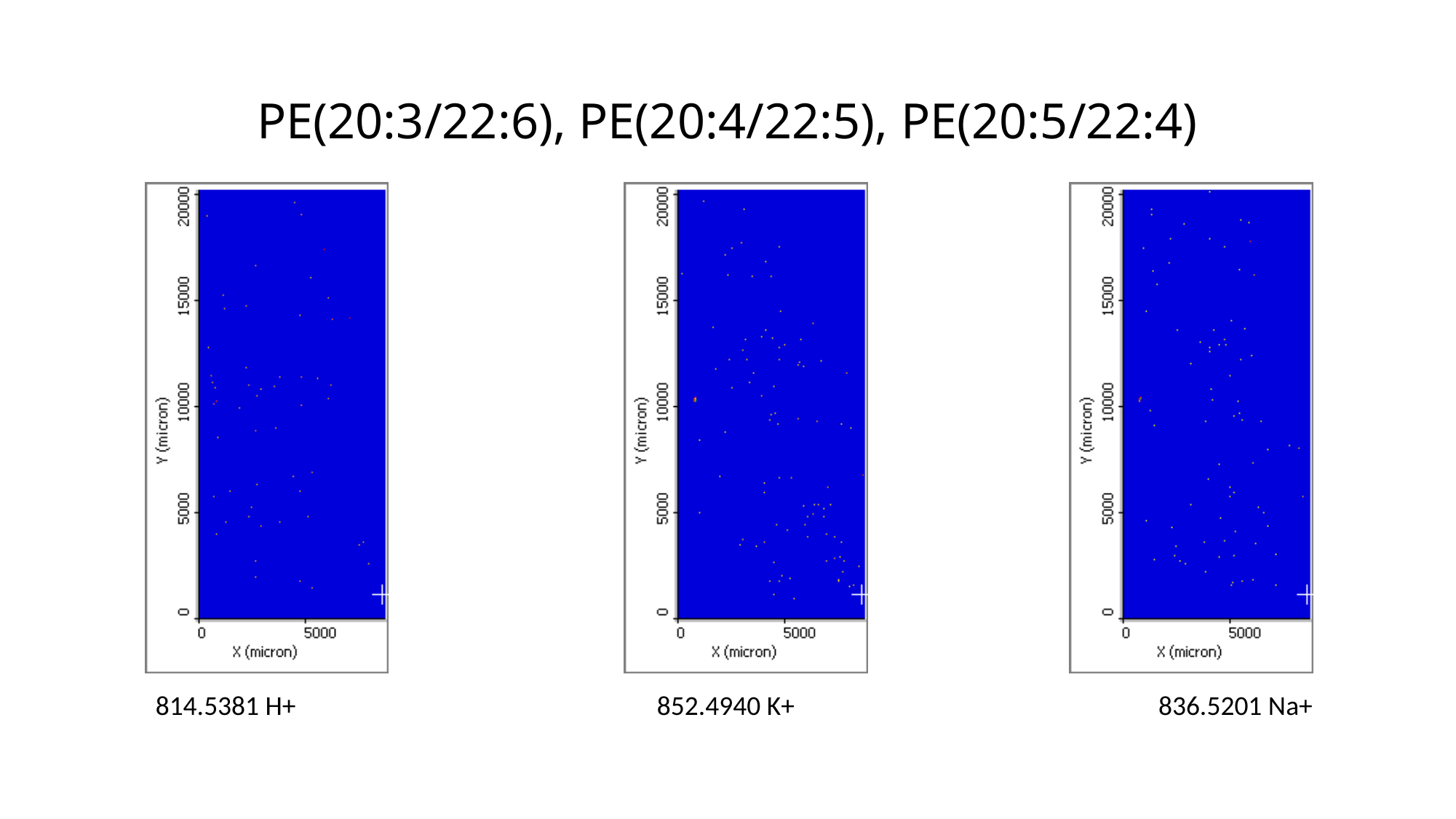

# PE(20:3/22:6), PE(20:4/22:5), PE(20:5/22:4)
814.5381 H+
852.4940 K+
836.5201 Na+

### Slide 61
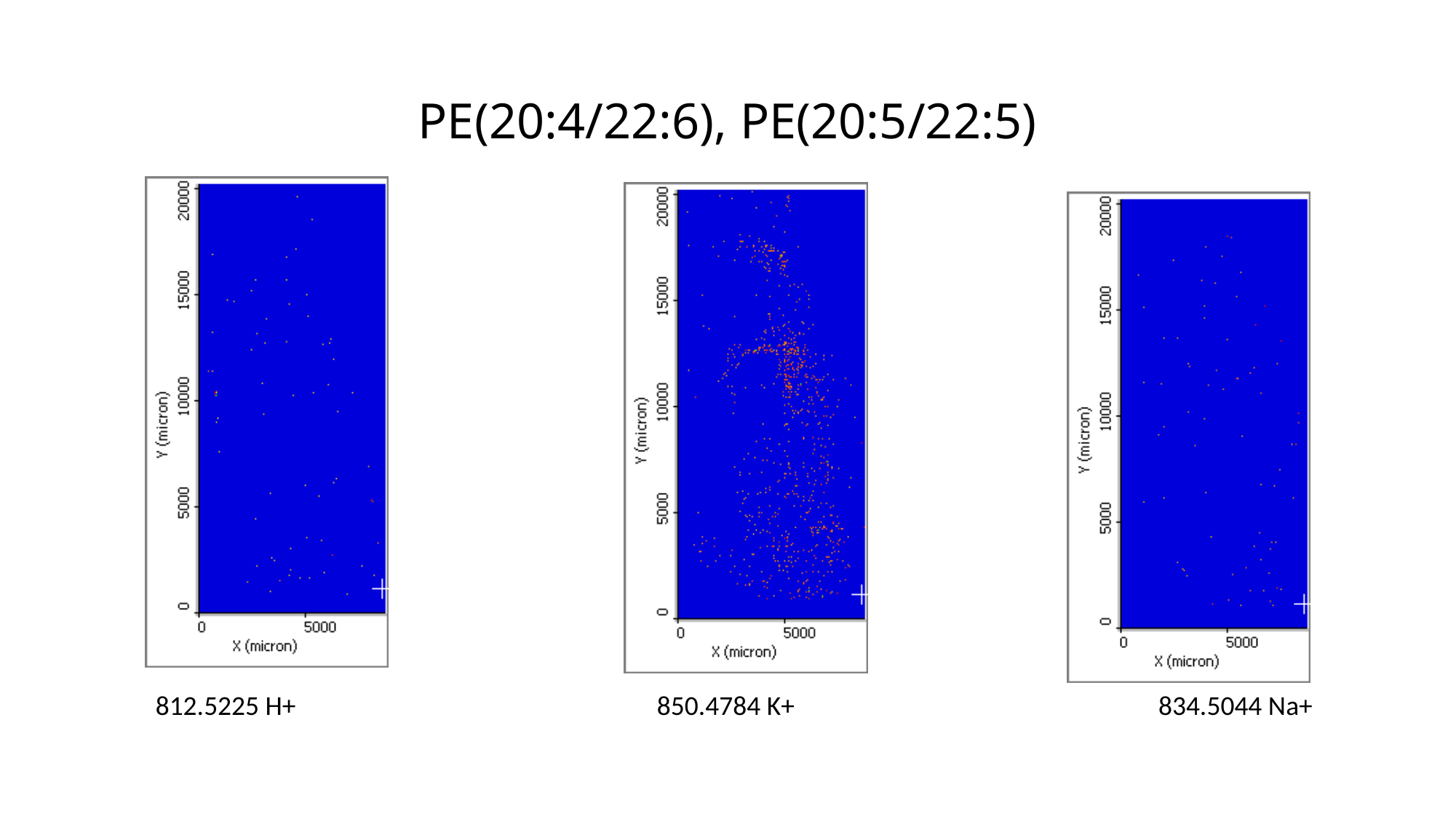

# PE(20:4/22:6), PE(20:5/22:5)
812.5225 H+
850.4784 K+
834.5044 Na+

### Slide 62
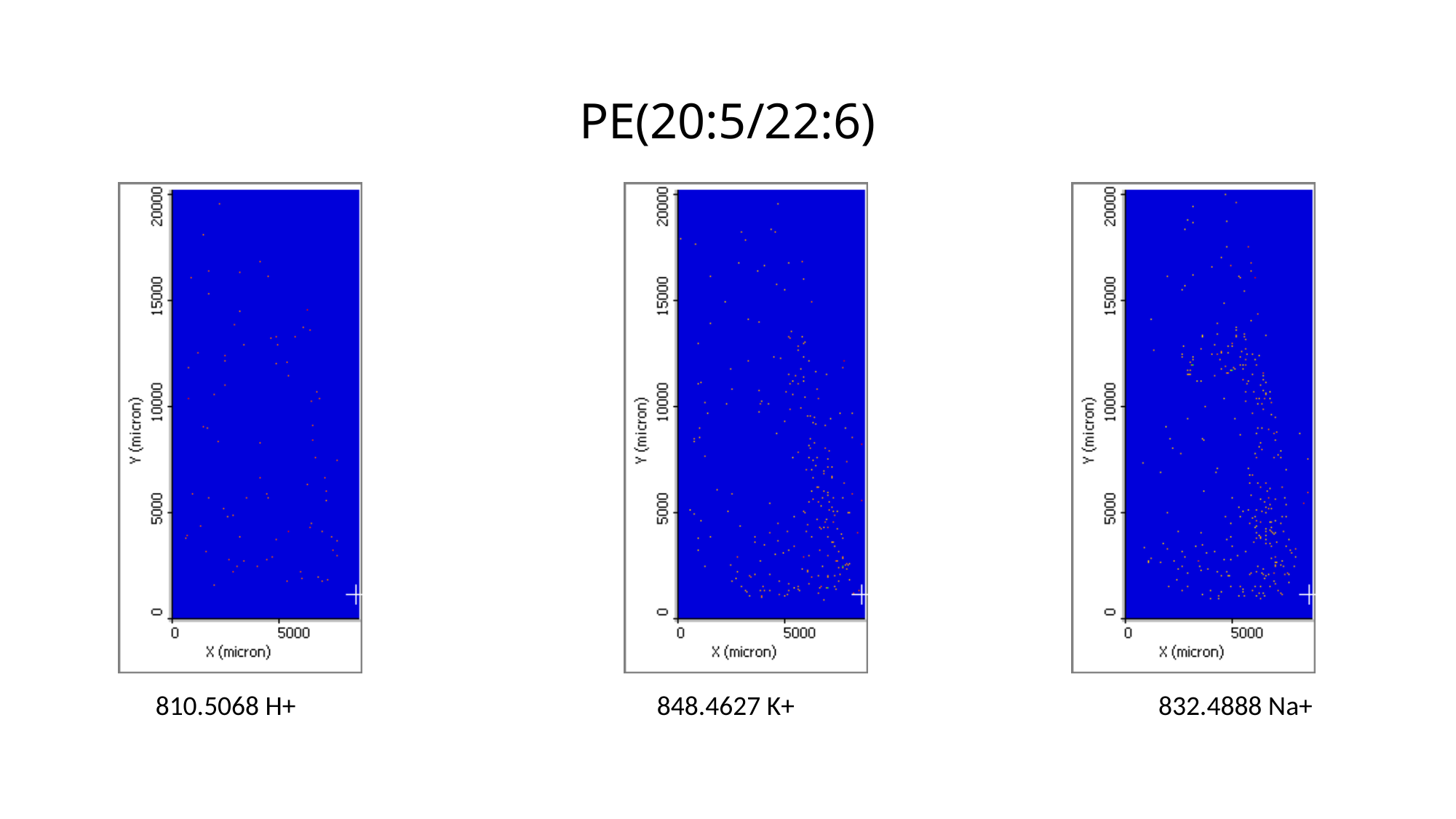

# PE(20:5/22:6)
810.5068 H+
848.4627 K+
832.4888 Na+

### Slide 63
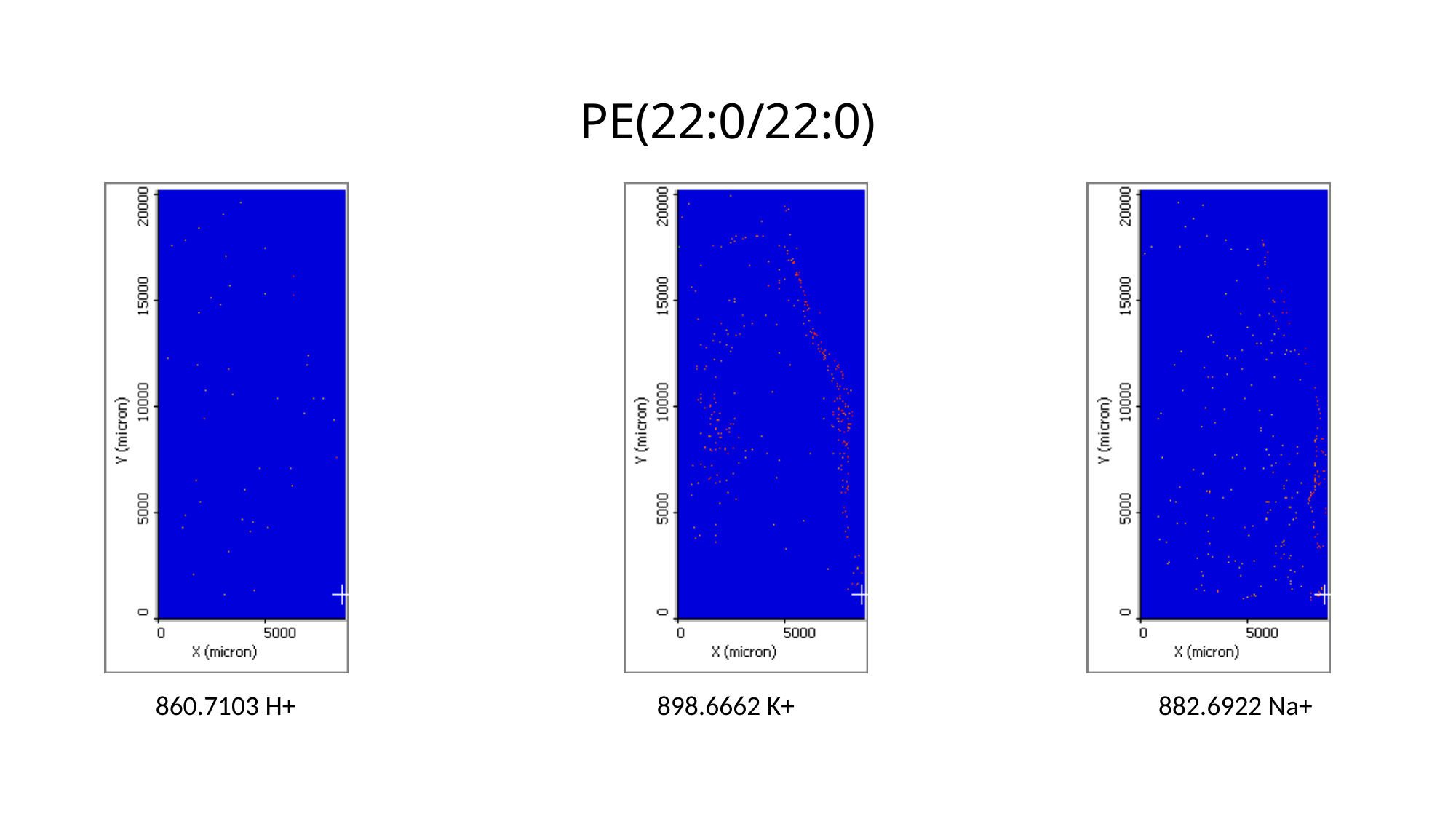

# PE(22:0/22:0)
860.7103 H+
898.6662 K+
882.6922 Na+

### Slide 64
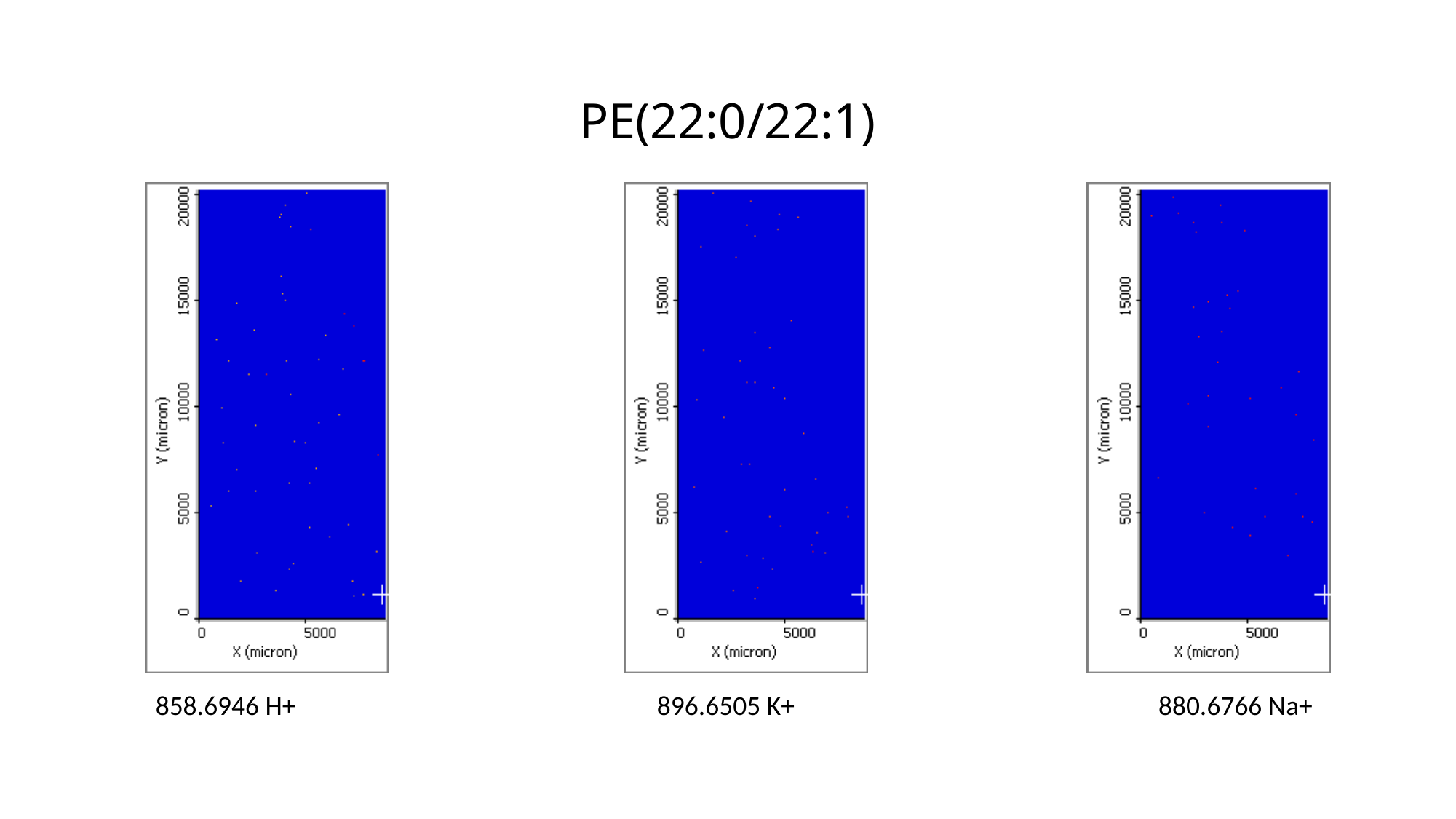

# PE(22:0/22:1)
858.6946 H+
896.6505 K+
880.6766 Na+

### Slide 65
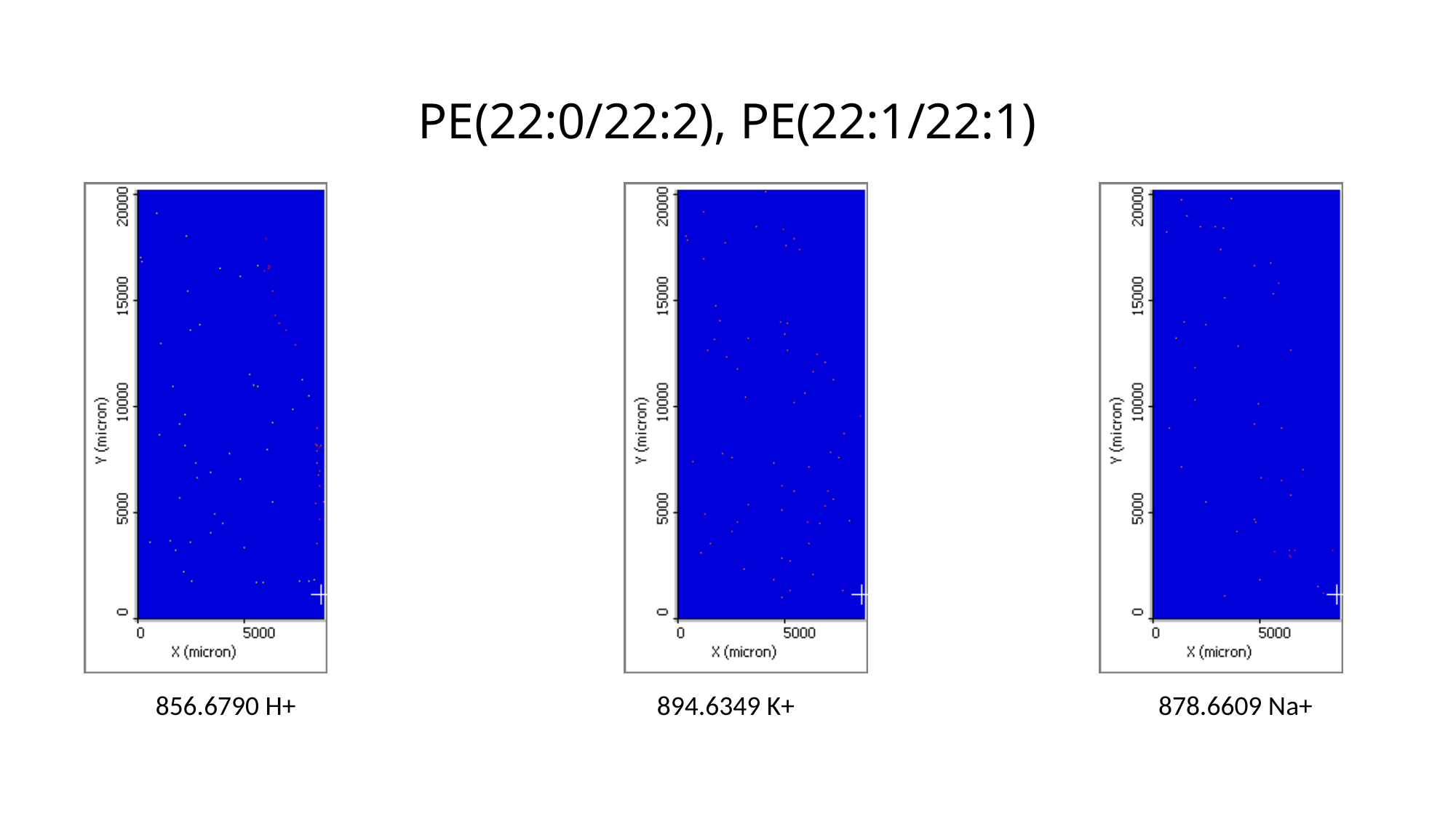

# PE(22:0/22:2), PE(22:1/22:1)
856.6790 H+
894.6349 K+
878.6609 Na+

### Slide 66
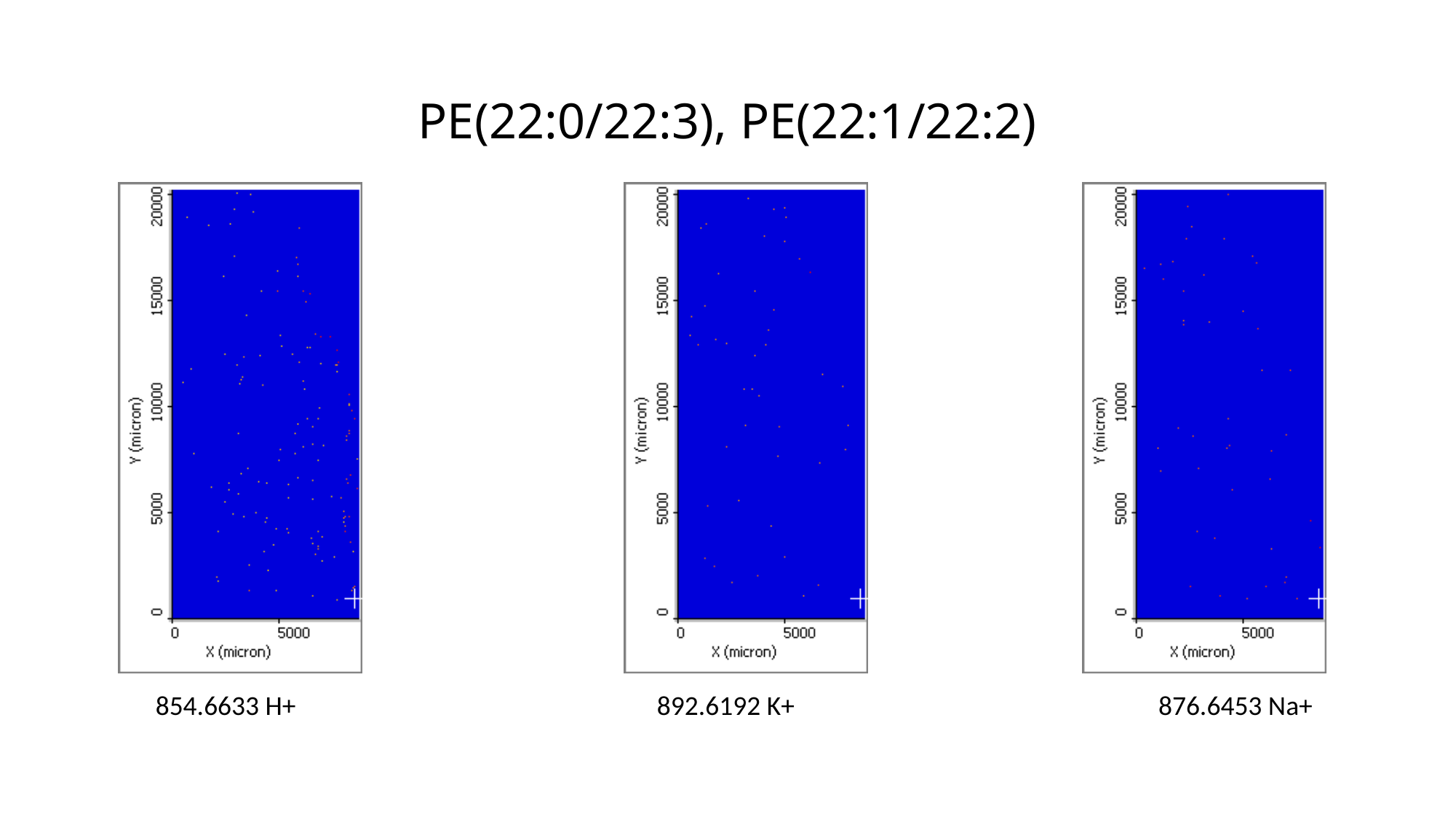

# PE(22:0/22:3), PE(22:1/22:2)
854.6633 H+
892.6192 K+
876.6453 Na+

### Slide 67
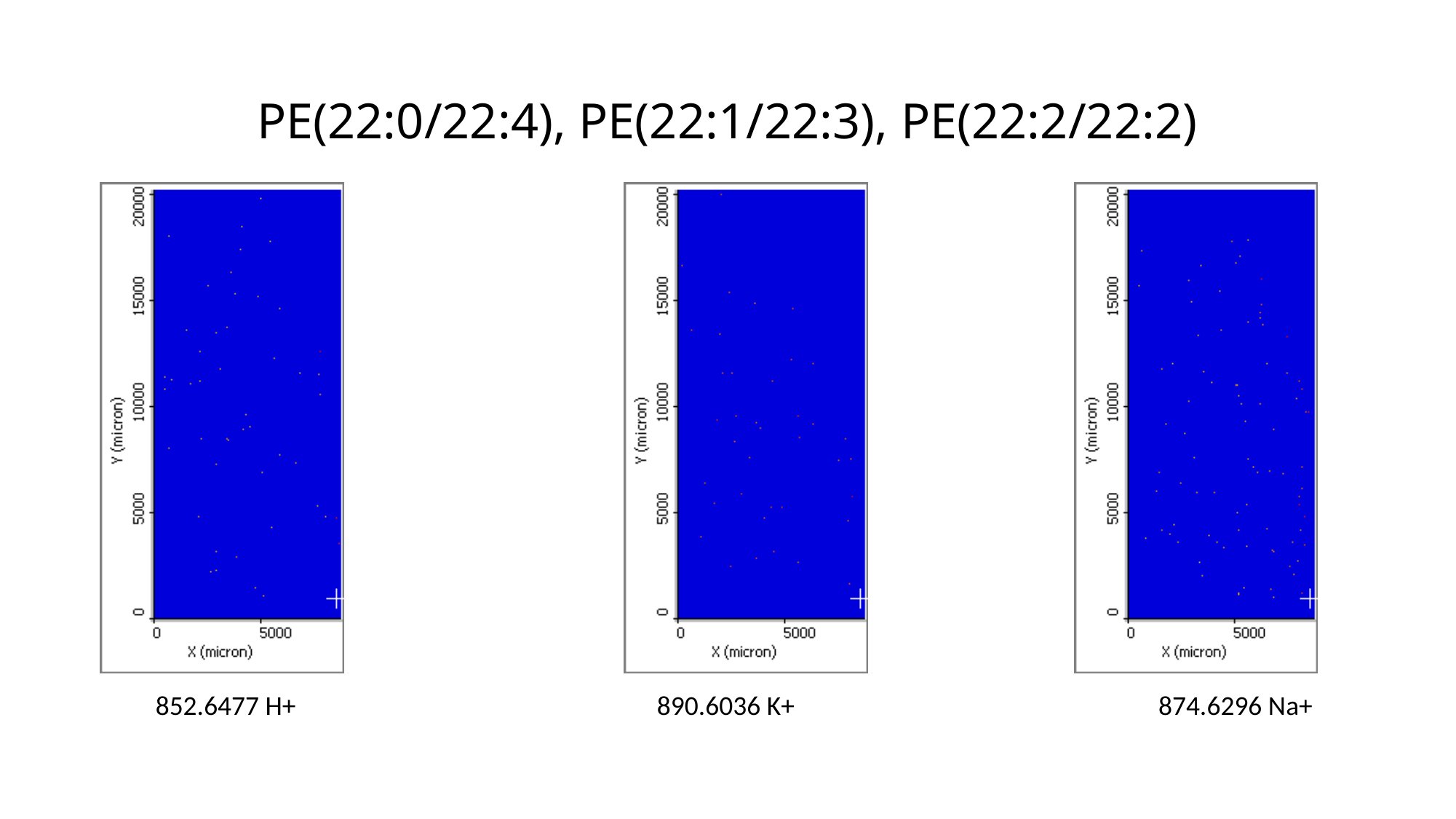

# PE(22:0/22:4), PE(22:1/22:3), PE(22:2/22:2)
852.6477 H+
890.6036 K+
874.6296 Na+

### Slide 68
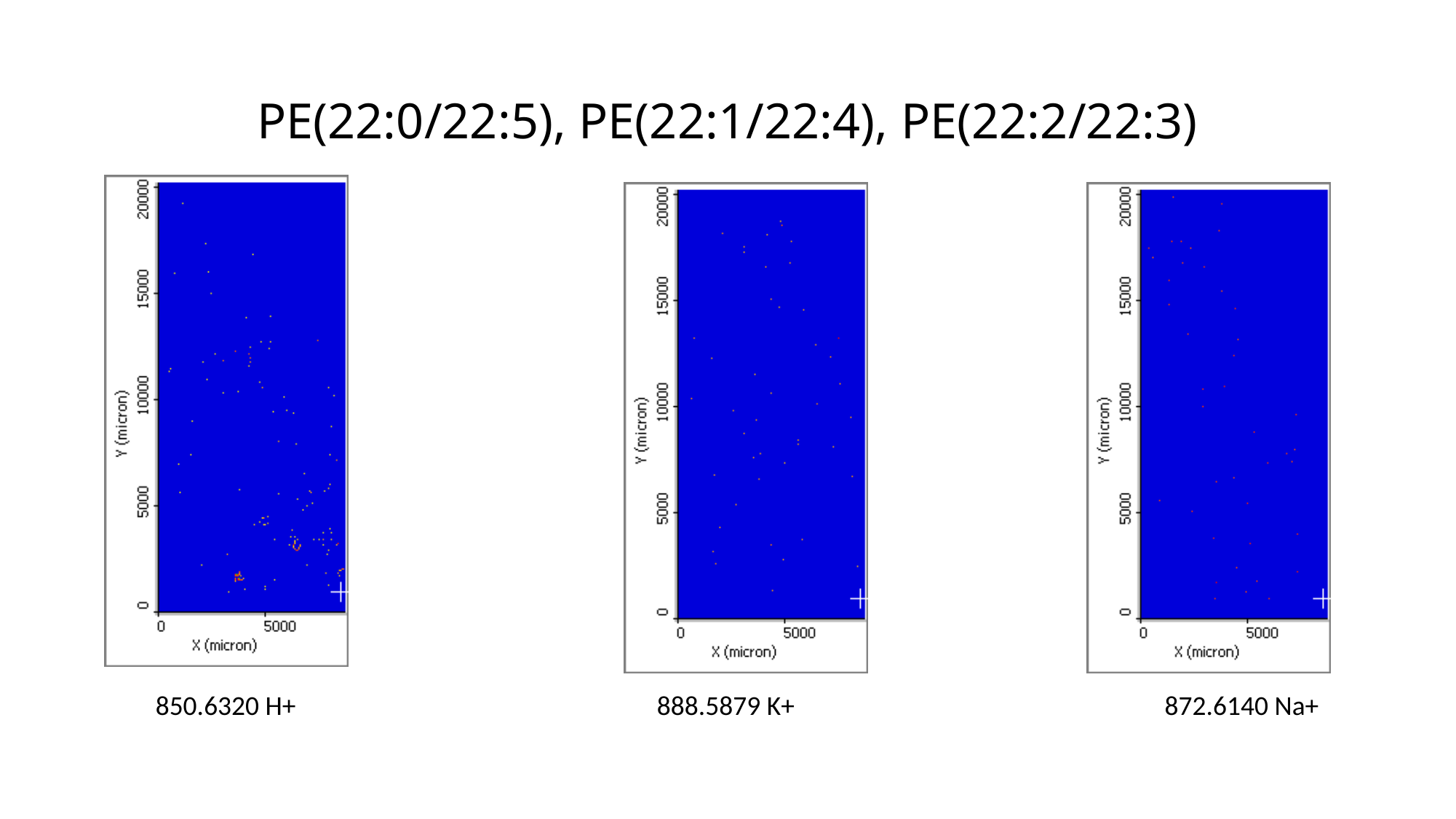

# PE(22:0/22:5), PE(22:1/22:4), PE(22:2/22:3)
850.6320 H+
888.5879 K+
 872.6140 Na+

### Slide 69
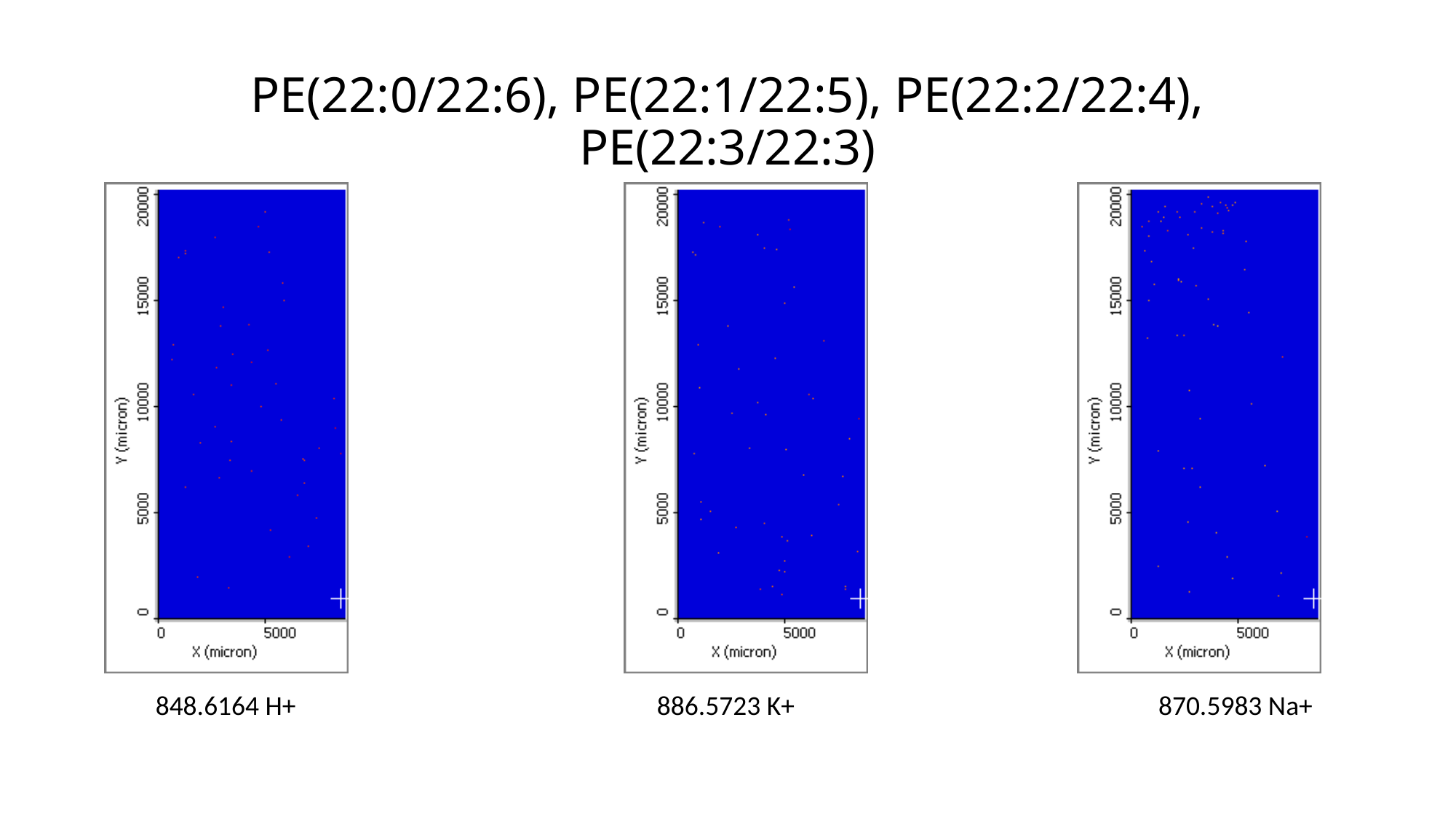

# PE(22:0/22:6), PE(22:1/22:5), PE(22:2/22:4), PE(22:3/22:3)
848.6164 H+
886.5723 K+
870.5983 Na+

### Slide 70
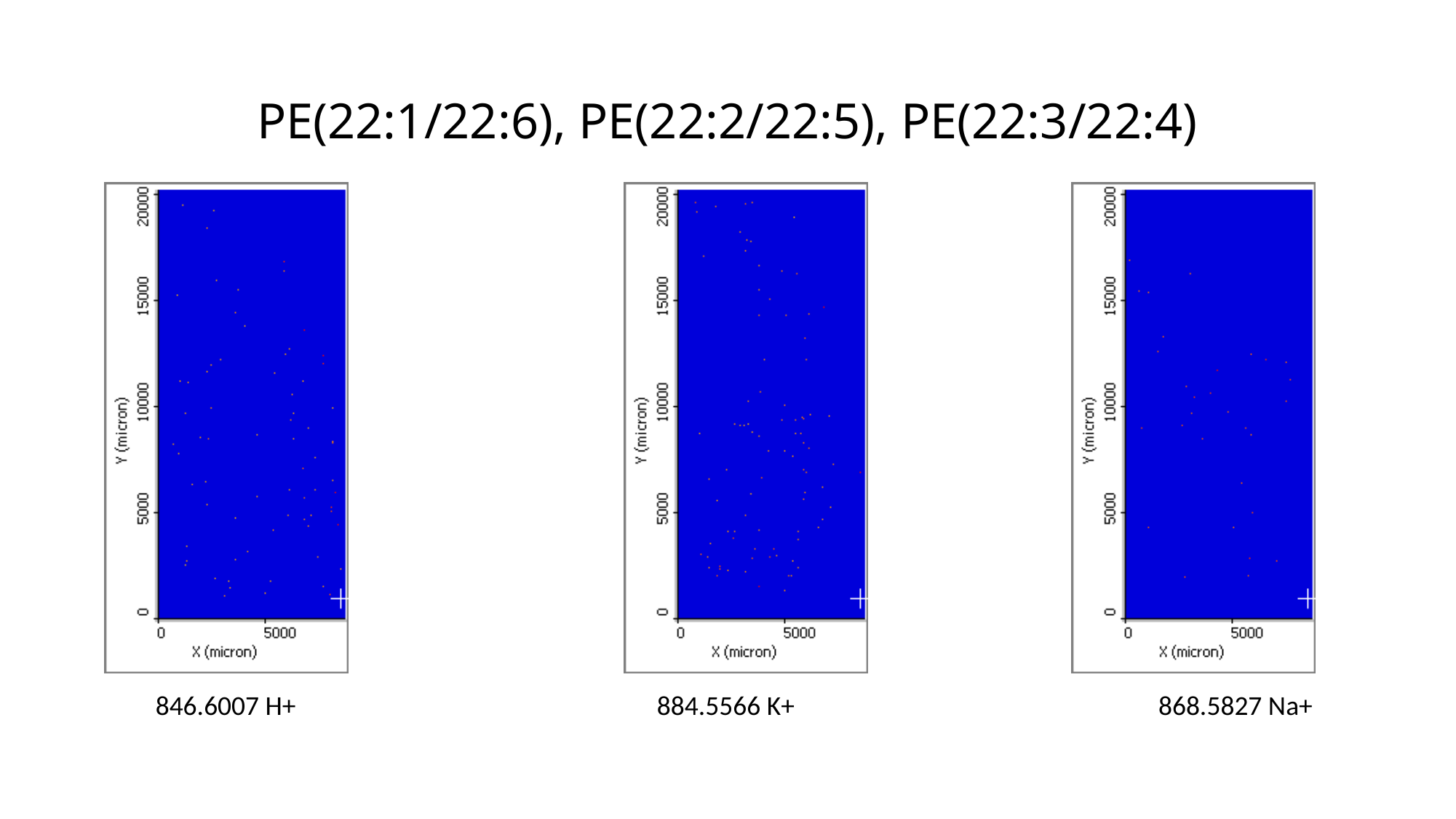

# PE(22:1/22:6), PE(22:2/22:5), PE(22:3/22:4)
846.6007 H+
884.5566 K+
868.5827 Na+

### Slide 71
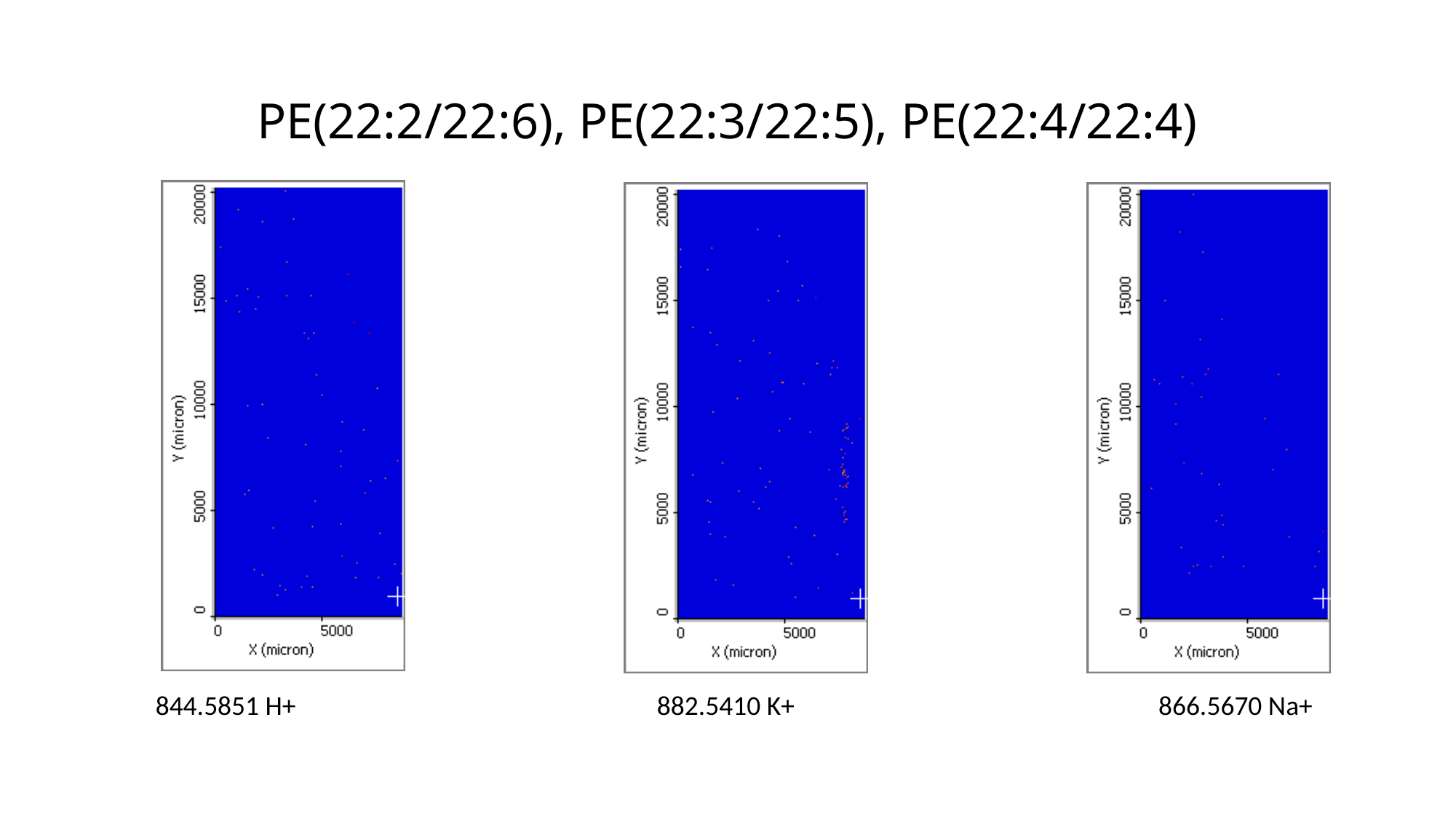

# PE(22:2/22:6), PE(22:3/22:5), PE(22:4/22:4)
844.5851 H+
882.5410 K+
866.5670 Na+

### Slide 72
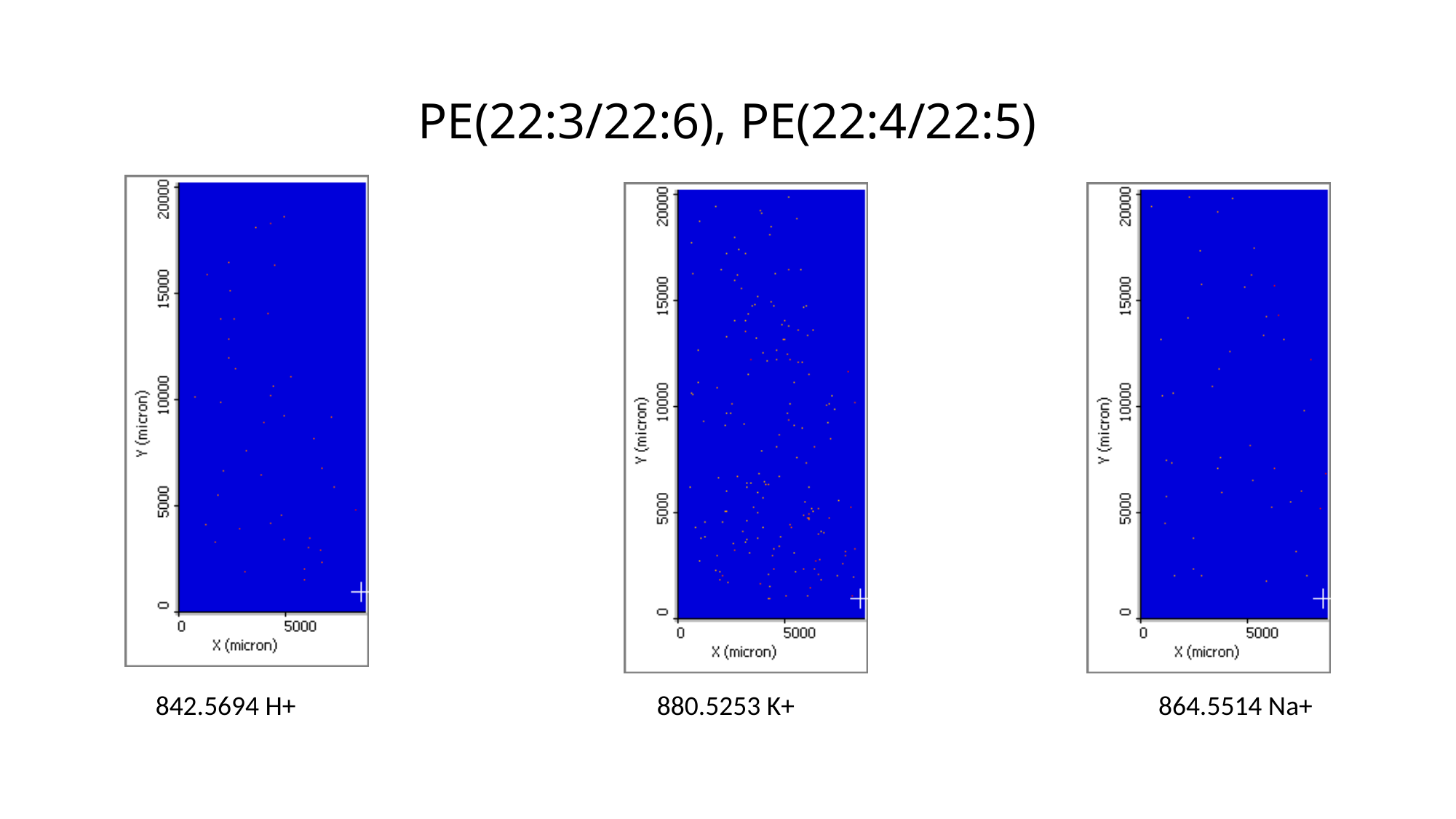

# PE(22:3/22:6), PE(22:4/22:5)
842.5694 H+
880.5253 K+
864.5514 Na+

### Slide 73
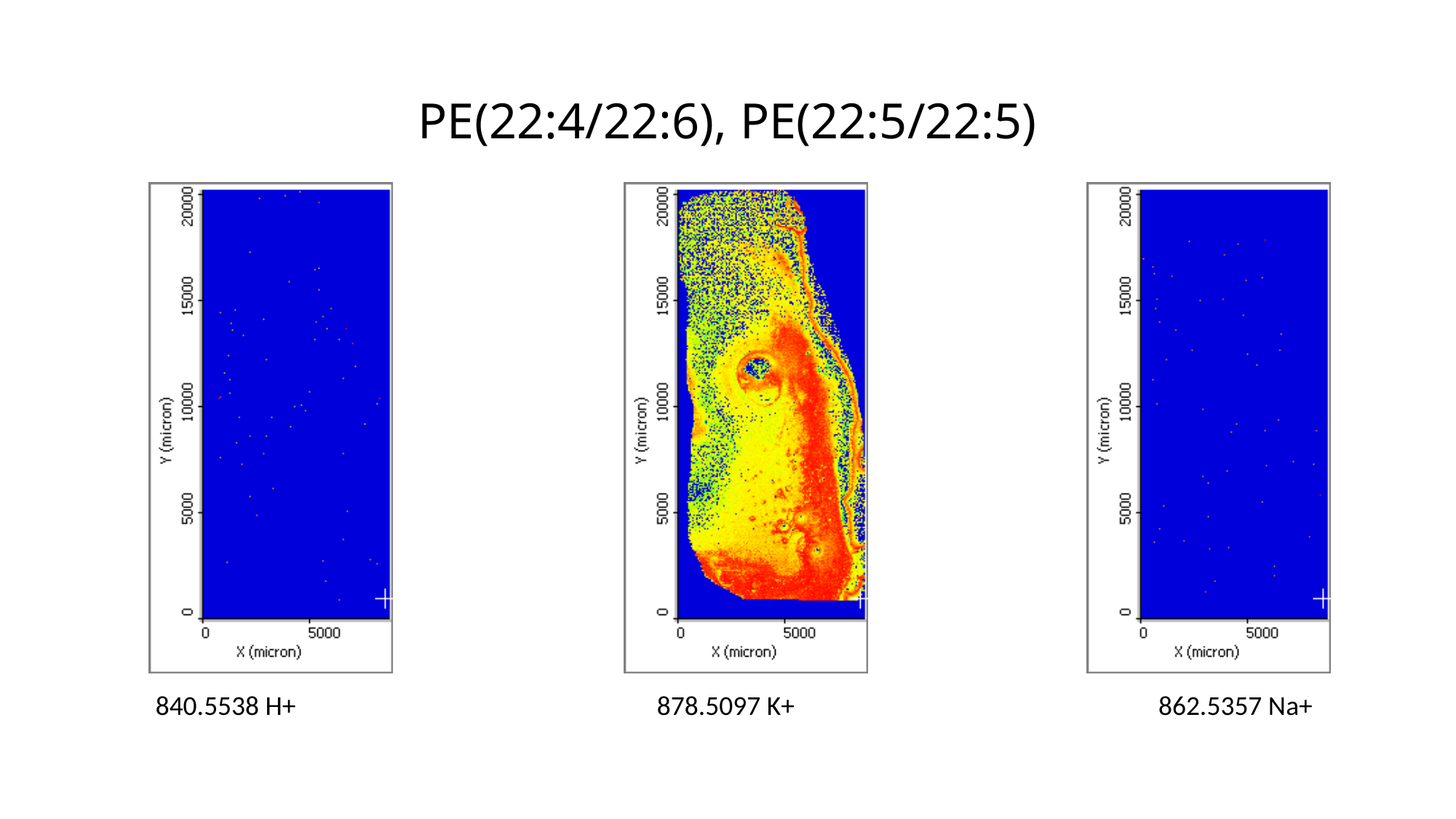

# PE(22:4/22:6), PE(22:5/22:5)
840.5538 H+
878.5097 K+
862.5357 Na+

### Slide 74
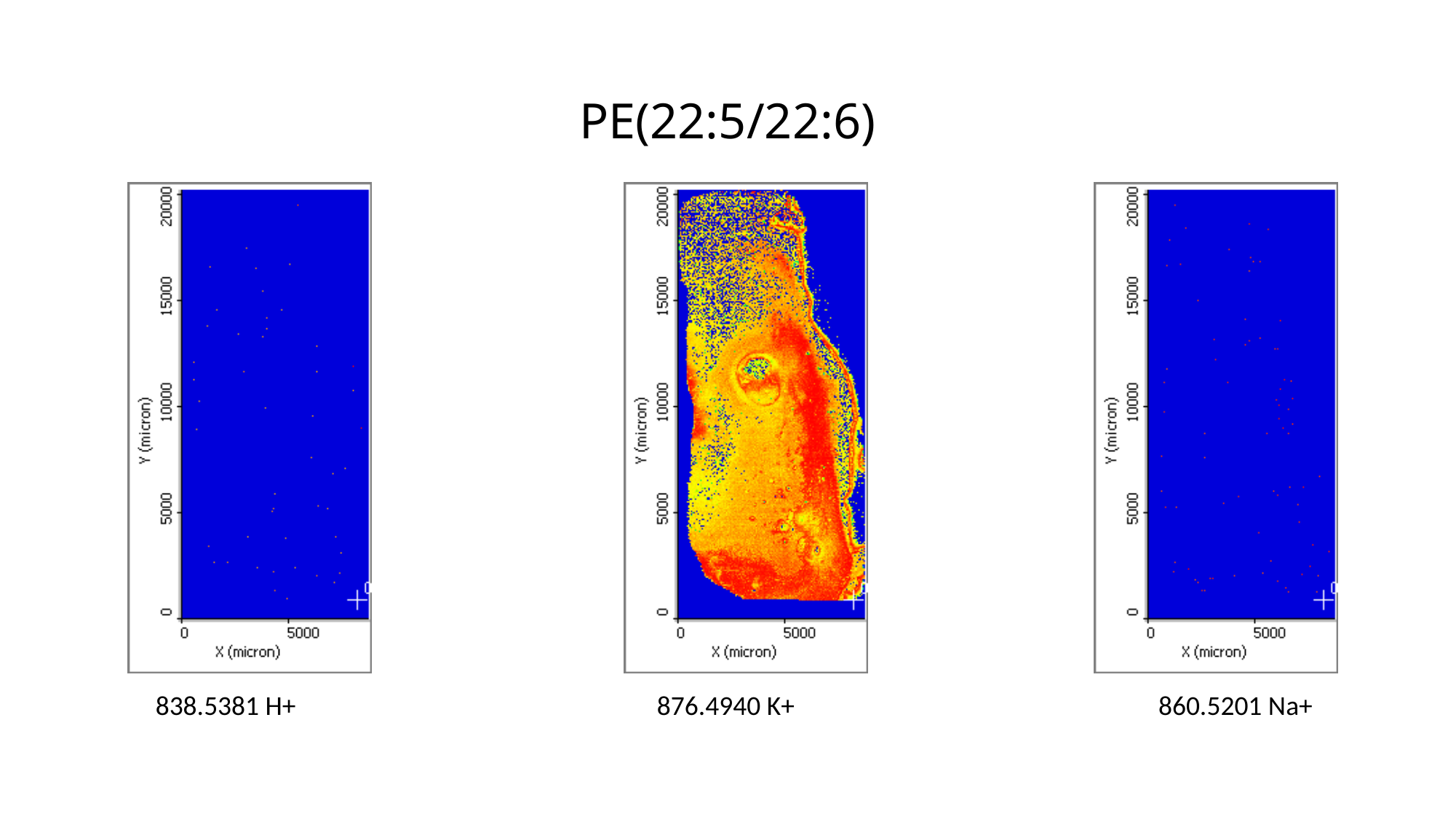

# PE(22:5/22:6)
838.5381 H+
876.4940 K+
860.5201 Na+

### Slide 75
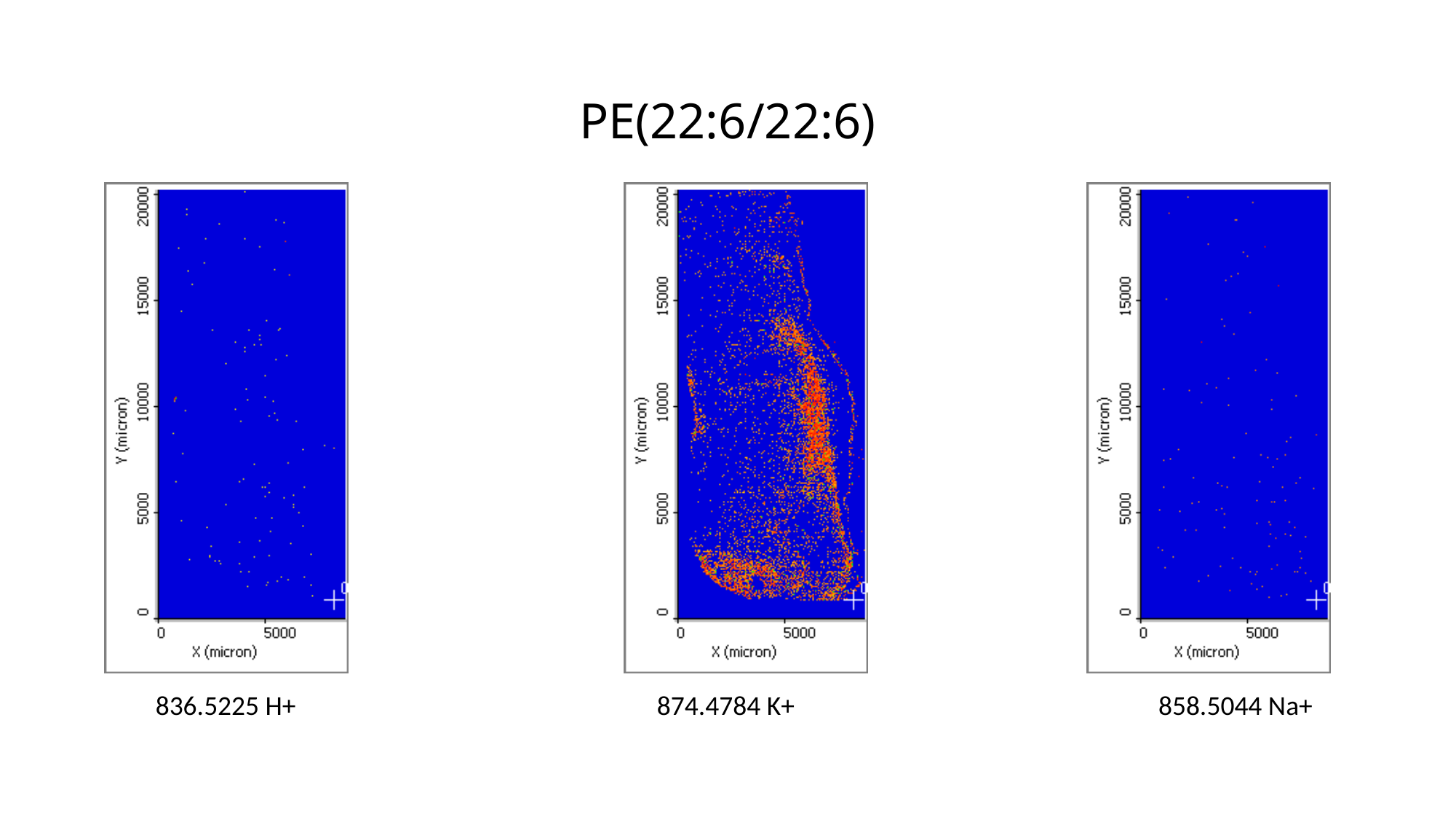

# PE(22:6/22:6)
836.5225 H+
874.4784 K+
858.5044 Na+
